# A novel *Arabidopsis thaliana* protein, ABAP1 INTERACTING PROTEIN 10, mediates crosstalk between the cell cycle and primary metabolism

**DOI:** 10.1101/2024.11.20.624481

**Authors:** Patrícia Montessoro, Joaquin Felipe Roca Paixão, Carinne N M Costa, Laura Ducatti, Letícia P Perdigão Grangeiro, Adriana Flores Fusaro, Helkin F Ballesteros, Vanessa Iurif, Luiz Mors Cabral, Jelmir Craveiro de Andrade, Leticia Tessaro, Wallace de Paula Bernardo, Fernanda Silva Coelho, Janice de Almeida-Engler, Eliemar Campostrini, Carlos Adam Conte-Junior, Adriana Silva Hemerly

## Abstract

Plants have developed a sophisticated regulatory network that coordinates gene expression in meristematic zones in response to environmental conditions. Here, we identified a protein in Arabidopsis (*Arabidopsis thaliana*) that interacts with Armadillo BTB Arabidopsis protein 1 (ABAP1), a negative regulator of the cell cycle in plants. We characterized ABAP1 Interacting Protein (named AIP10) investigating its role in modulating plant development. T-DNA insertion lines with silenced expression of *AIP10* were evaluated phenotypically (morphology, fresh and dry weight), via transcriptomics analyses (RNA-seq and RT-qPCR), physiologically (biochemically, Fluorcam and Li-COR), and metabolically (ATR-FTIR). We showed that AIP10 integrates cell division rates with transcriptional and primary metabolism reprogramming, through its protein interactions with ABAP1 and KIN10, a subunit of SnRK1 (Sucrose non-fermenting-1- related protein kinase 1). ABAP1 levels and activity were reduced in the absence of AIP10, licensing cell cycle progression for longer periods, which culminated in increased rates of cell division that boosted vegetative and reproductive growth. *AIP10* silencing triggered a major transcriptional reprogramming of plant primary metabolism, possibly through SnRK1 regulation. *aip10* mutants showed increased photosynthetic efficiency, as well as boosted carbon fixation, leading to increased biomass, seed productivity, and higher contents of proteins, lipids (triglycerides), and carbohydrates. Finally, we propose that the modulation of *AIP10* expression is part of a mechanism that coordinates higher rates of cell division with better photosynthetic performance and carbon fixation to metabolically meet the plant energy demand, allowing the generation of plants with increased biomass and productivity.

**One-sentence summary:** ABAP1 INTERACTING PROTEIN10 connects the cell cycle and plant metabolism, with its silencing leading to increased cell division, carbon fixation, and metabolite levels in Arabidopsis.

## Introduction

Plants have evolved distinct mechanisms to cope with environmental signals such as light, abiotic stress or pathogen attacks, which influence their architecture and plasticity (Pierik et al., 2021). Developmental plasticity strategies consist in altering morphological and physiological aspects to promote growth, survival and reproduction when confronted with adverse effects (Nawaz et al., 2023). The morphological response is mediated by regulating cell division rates at self-perpetuating meristems (Qi and Zhang, 2020). Cell division and differentiation at both root and shoot apical meristems are coordinated by hormone pathways, receptor kinase-peptide networks and transcription factor signaling (Qi and Zhang, 2020).

A typical plant mitotic cell division is accomplished in a cell cycle consisting of four different phases, G1 (postmitotic interphase), S (DNA synthesis phase), G2 (premitotic interphase), and M (mitosis/cytokinesis). The progression between the different phases is coordinated by key regulators, such as cyclin-dependent kinases (CDKs) complexed to cyclins (CYC), and checkpoint regulators, to ensure the cell cycle is ready to proceed to a new phase (Qi and Zhang, 2020). An important control that integrates the cell cycle with the environment is at the G1/S transition, when cells are licensed to initiate DNA replication, in a step regulated by the pre-replication complex (pre-RC) (Del Pozo et al., 2006). The pre-RC interacts with DNA and/or chromatin marks through the Origin recognition complex (ORC1-to-ORC6), that is assembled as a scaffold for sequential association of Cell Division Cycle 6 (CDC6), Chromatin Licensing and DNA Replication Factor 1 (CDT1) and Mini Chromosome Maintenance (MCM complex:MCM2–7), licensing DNA for replication (Brasil et al., 2017). Positive and negative proliferation signals, that often depend on environmental stimuli, control the response of the pre-RC machinery (Bailis and Forsburg, 2004).

In plants, we have previously characterized Armadillo BTB Arabidopsis protein 1 (ABAP1) as a plant-specific protein that binds directly to ORC1a/b and CDT1a/b in *Arabidopsis thaliana* (Masuda et al., 2008). ABAP1 negatively regulates the assembly of the pre-RC, which reduces DNA replication licensing and cell division rates. Its mechanism of action also involves association with <u>A</u>BAP1 <u>I</u>nteracting <u>P</u>roteins (named AIP) to reduce expression of specific genes, and negatively regulate cell proliferation rates during development (Masuda et al., 2008; Brasil et al., 2015; Cabral et al., 2021). Among the identified proteins interacting with ABAP1, some had unknown functions, such as AIP10. Previous studies evidenced putative orthologs of *AIP10* in *Physcomitrium patens* (*PpSKI1, PpSKI2*) and *Oryza sativa* (*OsHDR1*) that interact with SNF1/AMPK-related protein kinase 1 (SnRK1) homologues, *Pp*Snf1a and *Os*K4 kinases, respectively, to regulate growth, and photoperiodic flowering (Thelander et al., 2007; Sun et al., 2016). SnRK1 is a master regulator of energy homeostasis during sugar starvation, also controlling developmental plasticity and cellular responses to increase resilience under different stress conditions (Baena-González and Lunn, 2020). In Arabidopsis, the catalytic subunits of SnRK1 are encoded by three genes: *SnRK1α1* (KIN10), *SnRK1α2* (KIN11), and *SnRK1α3* (KIN12) (Soto-Burgos and Bassham, 2017).

Here, we describe an ABAP1 interactor that integrates modulation of the cell cycle and plant primary metabolism, adjusting plant plasticity to better respond to environmental signaling. We investigated the role of AIP10 in plant development by characterizing *A. thaliana* mutants with modified *AIP10* expression. Plants with silenced expression of *AIP10* (*aip10-1*) showed higher rates of cell proliferation in the meristems, for longer periods, leading to a vegetative and reproductive increase. AIP10 together with ABAP1 negatively regulate cell division, controlling the expression of ABAP1 target genes and cell cycle progression. RNAseq transcriptomics revealed that *AIP10* silencing leads to a major reprogramming of plant primary metabolism, possibly through SnRK1 regulation. Furthermore, *AIP10* silencing increased CO2 assimilation, as well as carbon fixation, resulting in a higher content of proteins, lipids (triglycerides) and carbohydrates. Finally, we propose that the modulation of *AIP10* expression is part of a mechanism that coordinates higher rates of cell division with better photosynthetic performance and carbon fixation, to metabolically meet its energy demand, allowing the generation of plants with a increase in biomass and gain in productivity.

## Results

### AIP10 interacts with ABAP1, an inhibitor of cell division in plants, and with SnRK1, a master regulator of plant environmental stimuli

The ABAP1 interacting protein 10 (At1g80940) was part of a set of proteins found during a yeast two hybrid (Y2H) screening against an *A. thaliana* cDNA library (Masuda et al., 2008; Brasil et al., 2015), using ABAP1 as a bait. A series of assays were performed to confirm *in vitro* and *in vivo* interaction between AIP10 and ABAP1 (Fig.1). A direct Y2H assay using either AIP10 or ABAP1 as baits (BD) showed interaction between the two proteins (Fig. S1). The same Y2H assay showed that AIP10 did not interact with all preRC members tested (ORC1-6, CDC6 and CDT1, except MCM2–7), neither with ARIA, an ABAP1 homolog. Interactions between ABAP1 and AIP10 were also confirmed *in vitro*, in GST (Gluthatione *S*-transferase) pulldown experiments with GST::ABAP1 and *in vitro* translated AIP10^S35^. As shown in Fig. 1A, AIP10 bound to ABAP1-GST (lane 3), but not to GST alone (lane 2). We next performed an anti-ABAP1 co-immunoprecipitation assay with shoot, root and inflorescence protein extracts of plants expressing *promAIP10::AIP10-YFP* (Fig. 1B). These plants carry a construct containing the full length genomic *AIP10* gene that gives rise to a fluorescently tagged AIP10-YFP C-terminal fusion protein (Tian et al., 2004). A protein blot with antibodies against YFP exhibited a major 50 KDa band at lanes 1, 5 and 9, indicating the *in vivo* interaction between AIP10 and ABAP1 in different plant organs (Fig. 1B). Other observed positive bands may represent isoforms of the AIP10 protein, or phosphorylated forms of the protein (see below, Fig. 1B).

**Figure. 1.**
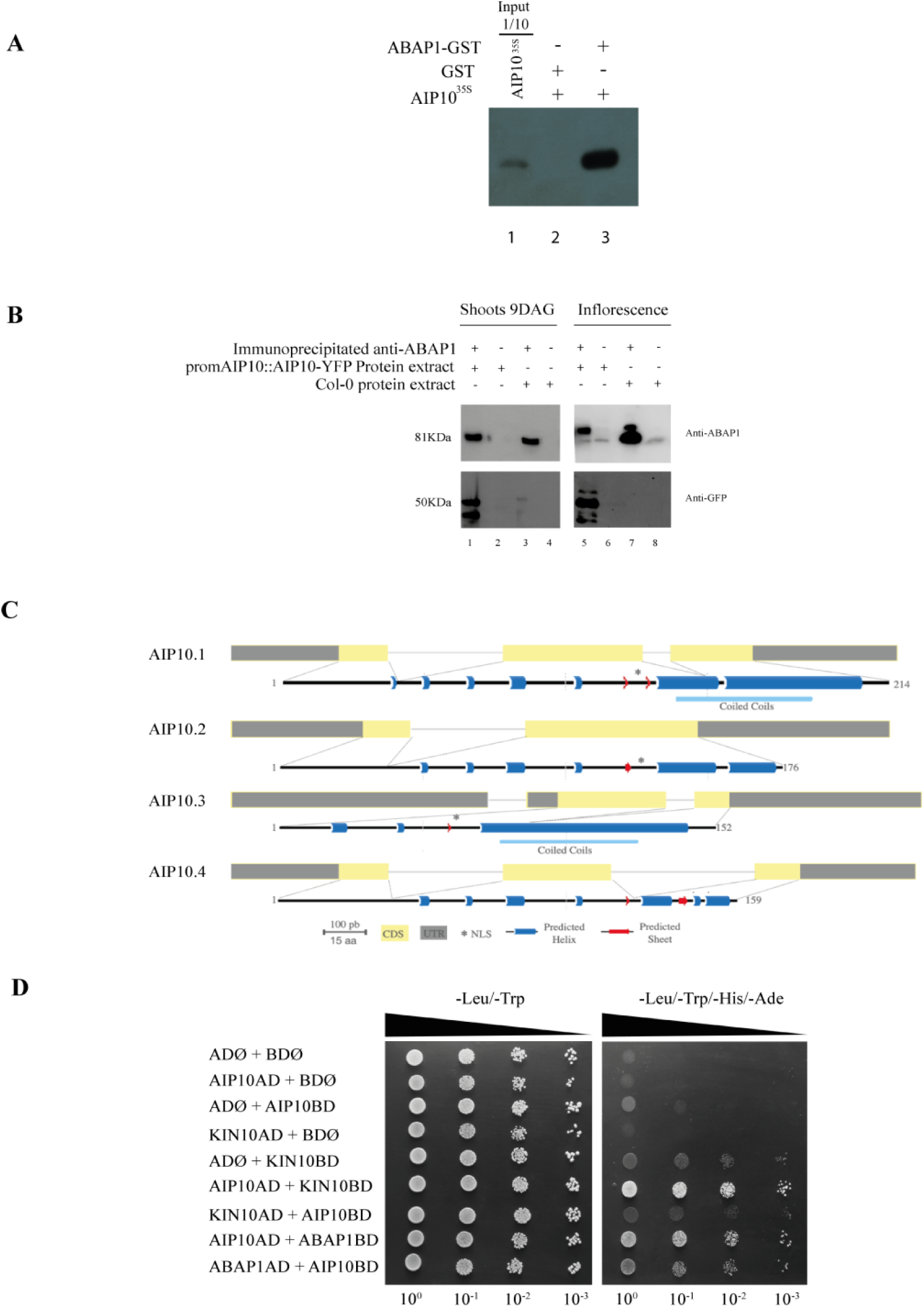
AIP10 interacts with ABAP1 *in vitro* and *in vivo.* **A)** ABAP1-AIP10 interaction through *in vitro* GST-pulldown assay of bacterially expressed recombinant ABAP1-GST fusion tested against *in vitro*-translated AIP10^S35^. ABAP1 and AIP10 interaction is shown in lane 3. **B)** Co-Immunoprecipitation (co-IP) of ABAP1 and protein extracts of plants expressing AIP10-YFP. Reactive bands corresponding to ABAP1 and AIP10-YFP were detected in shoot (lane 1), root (lane 5) and inflorescence (lane 9), by western blot using anti-ABAP1 and anti-GFP antibodies, respectively. The GFP antibody also recognizes the YFP protein. Numbers 1 to 12 refer to lanes in the SDS-PAGE. **C)** Schematic representation of AIP10 splice variants, showing the exons as gray (UTR) and yellow (CDS) boxes. Predicted Nuclear Localization Signal (NLS) is indicated as *, and predicted helix or sheet are indicated in blue or red, respectively. **D)** Yeast two-hybrid assay for protein interaction analysis between AIP10 and KIN10 fused to the AD and BD domains. Empty vectors pDEST22 (AD) and pDEST32 (BD) were used as negative controls, while AIP10 + ABAP1, fused to AD/BD, were used as a positive control. - Leucine/- Tryptophan media selected co-transformation with both BD/AD constructs; - Leucine/-Tryptophan/-Histidine/-Adenine selected for strong protein interaction. AIP10AD showed strong interaction with KIN10BD. GST, Glutathione S-Transferase; YFP, Yellow Fluorescent Protein; GFP, Green Fluorescent Protein; UTR, Untranslated Region; CDS, Coding Sequence; AD, Activation Domain; BD, Binding Domain.

The *AIP10* gene generates four isoforms through alternative splicing (Fig. 1C), in which all isoforms showed high expression levels in young leaves and reproductive organs (Fig. S2). AIP10.1 has 214 amino acids (24 kDa), and AIP10.2, AIP10.3 and AIP10.4 are splice variants with 176 aa (19 kDa), 152 aa (17 kDa) and 159 aa (17 kDa), respectively. AIP10.1, AIP10.2 and AIP10.3 possess a nuclear localization sequence (NLS) and only AIP10.1 and AIP10.3 possess a predicted coiled coil domain on the C-terminal part of the protein (Fig. 1C).

AIP10 is a plant-specific gene distributed all over the plant kingdom, that has at least one copy in all the 136 terrestrial species analyzed (Fig. S3A). ABAP1, besides being distributed all over the plant kingdom, has representatives also in algae, with 511 homologues present in 139 species (Fig S4). The AIP10 protein family is classified in the InterPro database as Heading Date Repressor 1 (HDR1) which regulates flowering time in rice, in a photoperiod-dependent way (Sun et al., 2016). Two homologues have already been described as *PpSKI1/2* (Snf1-related Kinase Interacting protein 1/2) in *Physcomitrium patens* along with *OsHDR1* in *Oryza sativa* (Thelander et al., 2007; Sun et al., 2016). The alignment revealed conserved regions mainly at the C-terminal part of the protein, including an NLS present in all the sequences (red frame), and all sequences presented putative phosphorylation sites (PPS) (Fig. S3B).

A conserved sequence VDVVE**S**MRRI (green frame) adjacent to the NLS was identified, which corresponds to a canonical KIN10 phosphorylation site (MLVFI)X(RKH)XX(**S/T**)XXX(LFIMV) as described by Halford & Hardie (1998) (Fig. S3B). In accordance, *Pp*SKI1/2 and *Os*HDR1 interact with *Pp*Snf1a and *Os*K4 kinases, respectively (Thelander et al., 2007; Sun et al., 2016). To test if the *A. thaliana* AIP10 interact with SnRK1, yeast two-hybrid assays against two catalytic subunits of the kinase, KIN10 and KIN11, was performed (Fig. 1D, Fig. S1). The data showed a direct interaction between AIP10 and KIN10 (Fig. 1D), that what was not seen for KIN11(Fig. S1).

### AIP10 is predominantly expressed in the nuclei of cells from young organs

*A. thaliana* plants expressing *promAIP10::*AIP10-YFP were used to determine the expression of the AIP10 protein in plant tissues during development, as well as its subcellular localization, under the control of its own promoter. Confocal analysis showed that AIP10 is present in different organs and tissues, consistent with the expression data in the eFP Browser at the TAIR platform. During reproduction, AIP10 was detected in reproductive organs such as anthers (Fig. 2A, B) and ovaries (Fig. 2C, D), being present in male and female gametes. AIP10 has a constitutive pattern of expression at different stages of embryogenesis, as shown in globular (Fig. 2E), cordiform (Fig. 2F), torpedo (Fig. 2G) and mature embryo (Fig. 2H). It is present in the nuclei of cells in the cotyledon, in the first pair of leaves (Fig. 2I), in the developing trichomes (Fig. 2J) and in the nuclei of hypocotyl cells (Fig. 2K). In root, constitutive nuclear and perinuclear fluorescence was detected (Fig. 2 L-N). Fluorescence was not observed in the root cap but was present in the root apical meristem (RAM) nuclei (Fig. 2L). AIP10 fluorescence intensity changes at the periphery of nuclei throughout the root (Fig. 2M, N). To investigate the localization of AIP10 in dividing cells, nuclei were stained with DAPI (Fig. 2 O, P). The protein is located at the periphery of the nuclei in the different phases of the cell cycle (Fig. 2O), being absent in mitotic cells (Fig. 2P). Altogether, the data indicate that AIP10 is a protein present in all plant organs and meristems.

**Figure. 2.**
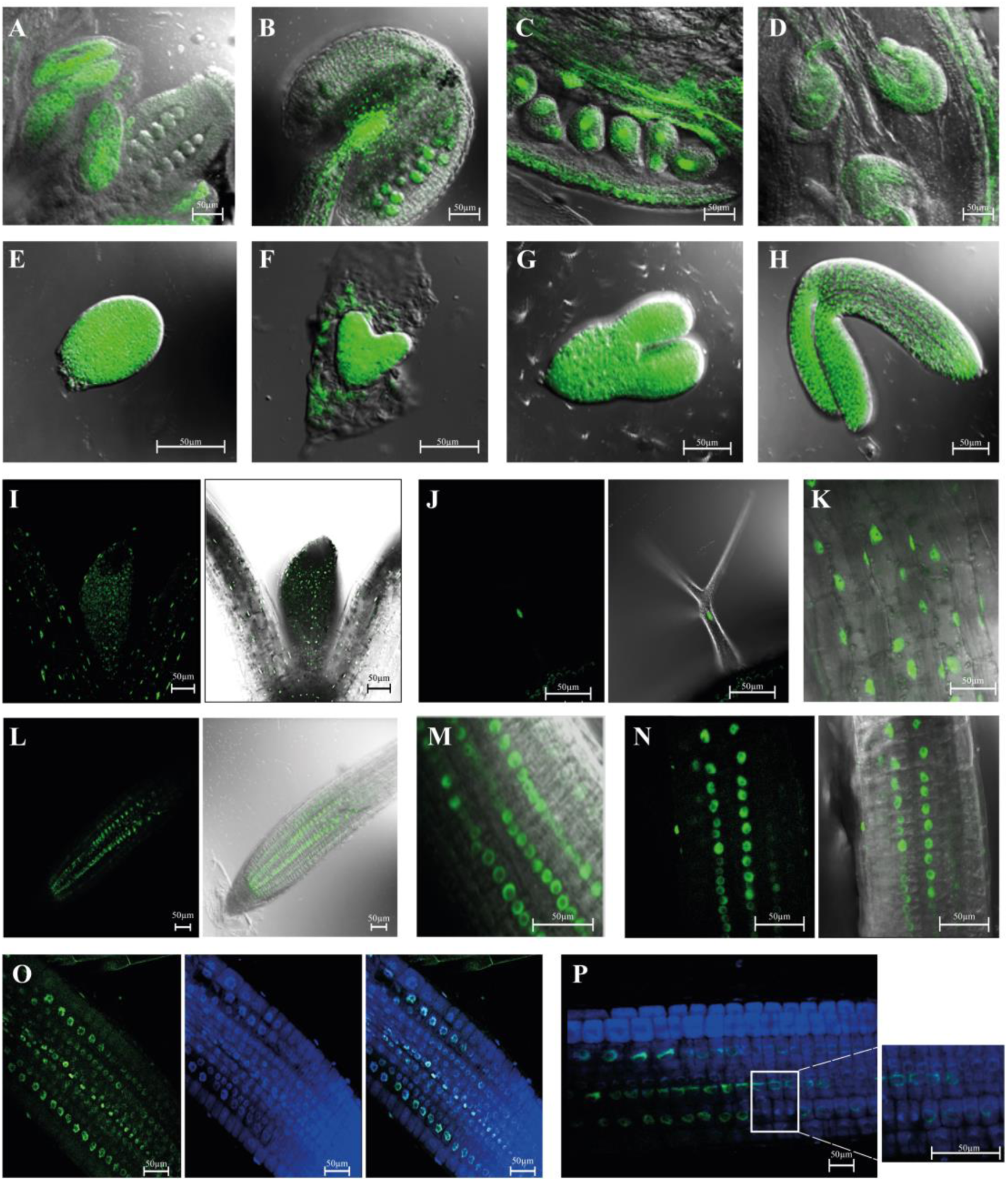
Tissue expression pattern and subcellular localization of fluorescent fusion protein AIP10-YFP, in Arabidopsis expressing *promAIP10::AIP10-YFP.* YFP signal was visualized by epifluorescence confocal microscopy. **A-H, K, M)** Merged images of epifluorescence confocal microscopy and bright-field differential interference contrast; **A, B)** anthers; **C, D)** ovaries; **E)** globular embryo**; F)** cordiform embryo, **G)** torpedo embryo and **H)** mature embryo. **I, J, L, N)** Epifluorescence confocal image in left panel, merged with bright-field differential interference contrast image in right panel; YFP signal in the nuclei of **I)** SAM; **J)** trichome; **K)** hypocotyl, **L)** RAM at 10 DAG; **M)** Nucleus and perinuclear region in root at 5 DAG; **N)** Nucleus and perinuclear region in root at 10 DAG. **O, P)** epifluorescence confocal microscopy images of root cells stained with 4,6-Diamidino-2-phenylindole (DAPI) to visualize the nucleus; **O)** YFP signal from RAM at 5 DAG; **P)** Magnification of RAM at 5 DAG showing that the protein is not present in dividing cells. Scale bars: 50 μm. RAM, Root Apical Meristem; SAM, Shoot Apical Meristem; YFP, Yellow fluorescent protein.

### Modulation of AIP10 levels regulates plant vegetative growth, biomass and productivity

To investigate the role of AIP10 in plant development, *A. thaliana* plants with altered AIP10 transcript levels were characterized. Two T-DNA insertion lines, Salk_022332 and Salk_094618, with the AIP10 gene knockout (*aip10-1*) or knockdown (*aip10-2)* (Fig. S5A, B), respectively, were used for phenotype comparisons.

At the vegetative state, *aip10-1 and aip10-2* plants showed a larger rosette area compared to Col-0 (Fig. 3A, B, D), formed by larger leaves, in significantly higher numbers (Fig. 3E). This was clearly demonstrated in leaf series analysis (Fig.3 C, F). Increased growth in the *aip10* mutants led to a significant increase in fresh and dry weight (Fig.3G, H). Plants with reduced levels of *AIP10* showed an earlier transition from the vegetative to the reproductive stage when compared to Col-0 plants (Fig 3B; I) under long day (LD) conditions (16h light/8h dark). This phenotype was more exacerbated in *aip10- 2* plants (Fig.3B), which corroborates observations in the *hdr1* mutant (AIP10 homolog in rice) that flowered 30 days before Wt under natural and long-day conditions (Sun et al., 2016).

**Figure. 3.**
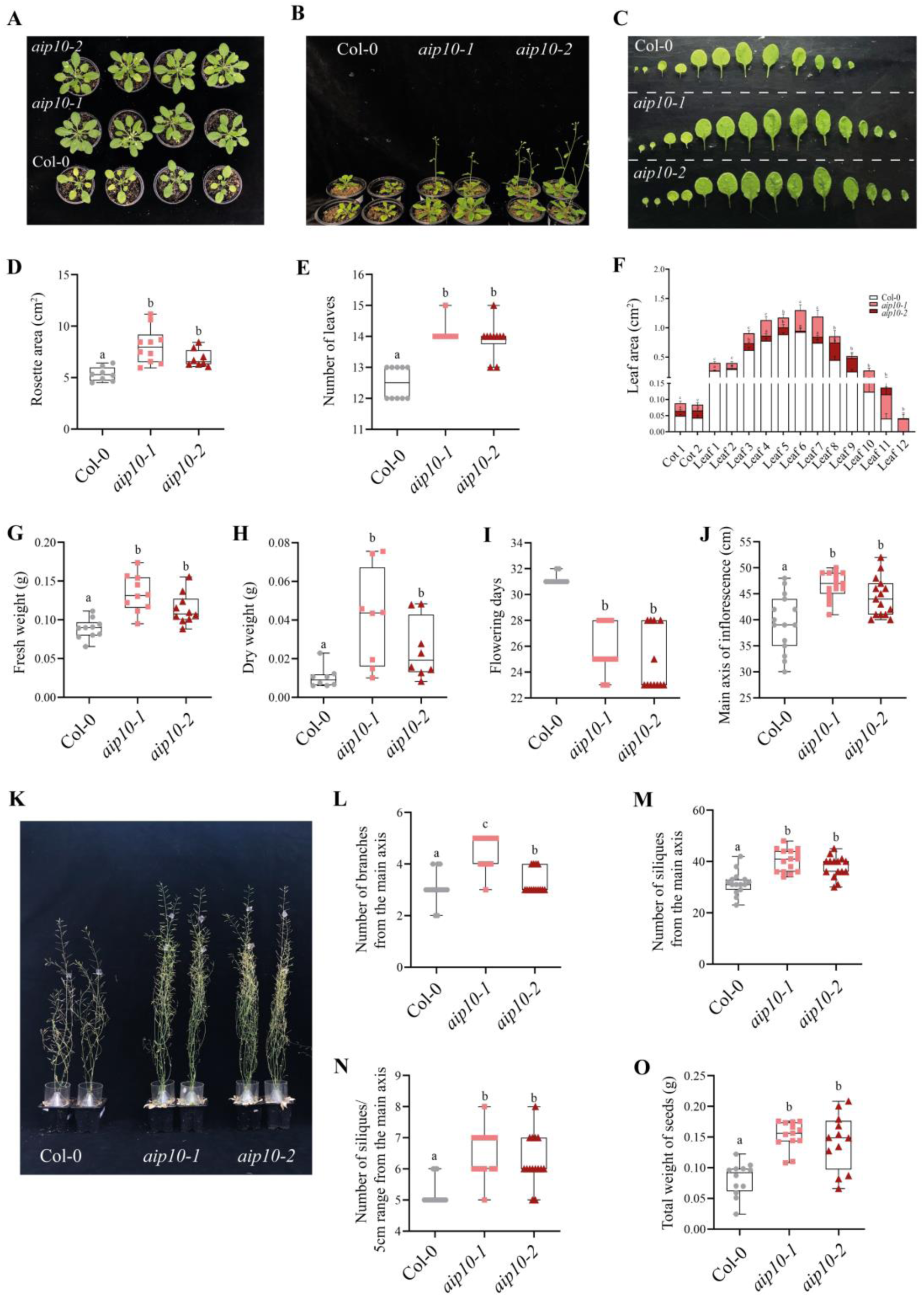
Phenotypic analyzes of AIP10 knock-out (*aip10-1*) and knock-down (*aip10-2*) Arabidopsis plants compared to wild-type Col-0. Experiments carried out, in a photoperiod of 16h/8h at 21°C, cultivated directly in the soil. Representative images of **A)** wild-type Col-0, *aip10-1* and *aip10-2* plants at 30 DAG and **B)** 35 DAG. **C)** Analysis of the leaf series of plants at 20 DAG to assess leaf area. **D)** Rosette area measurement using ImageJ program. **E)** Number of leaves at 32 DAG. **F)** Measurement of leaf area at 22 DAG using ImageJ software. **G)** Whole plant fresh weight and **H)** Whole plant dry weight at 32 DAG. **I)** Flowering days. **J)** Height of the main inflorescence axis at 57 DAG. **K)** Representative images of Col-0, *aip10-1* and *aip10-2* plants at 57 DAG during the reproductive stage. **L)** Number of branches on the main inflorescences axis at 40 DAG. **M)** Total number of siliques in the main inflorescence axis at 57 DAG. **N)** Measurement of the number of siliques at a fixed interval of 5cm on the main inflorescence axis at 57 DAG. **O)** Total seed weight per individual. The box plot graphs show the distribution of data of 10 individuals for biomass and 15 individuals for reproductive analyzes. The box represents the interquartile range (IQR), with the inner line running down the median. The whiskers extend to the maximum and minimum values, including all points within that range. The difference between groups in all analysis was confirmed by One-Way ANOVA (p < 0.05), and different letters indicate statistically different means according to Tukey’s test at 5% probability. DAG, Days After Germination.

At the reproductive stage, *aip10* mutants showed higher main inflorescence axis at the silique ripening stage (57 DAG), with a 16% and 12% significant increase compared to Col-0, respectively (Fig. 3 J-K). *aip10* mutant plants grew and generated the reproductive organs with at least 20% more ramifications and siliques of the main inflorescence axis, than Col-0 (Fig. 3L-N) and produced at least 30% more seeds compared to control plants (Fig. 3O). Furthermore, *aip10* plants showed a delay in senescence, suggesting that meristems remained active for longer periods contributing to the improved plant growth and production of plant organs.

Altogether, the gene expression and mutant phenotype analyses suggested that AIP10 acts throughout development, as it is present in all organs and meristems, and silencing of its expression lead to growth promotion. Next, we investigated molecular and biochemical mechanisms by which downregulation of AIP10 promotes plant development.

### AIP10 is a negative regulator of cell divisions that controls expression of ABAP1 target genes

As AIP10 interacts with ABAP1, a regulator of cell division in plants, and *aip10* mutants showed increased growth, the transcriptional regulation of two direct targets of ABAP1 repression at G1/S transition, *CDT1a* and *CDT1b* genes, were investigated by RT-qPCR. Remarkably, in *aip10-1* plants, *ABAP1* expression was 50% lower in 11 DAG plants and 70% lower in 35 DAG plants. Accordingly, *CDT1a* and *CDT1b*, were increased by 90% and 20%, respectively in 11 DAG and 35DAG (Fig. 4A, B). This data suggests that AIP10 could act in a complex with ABAP1, repressing gene expression. Also, AIP10-ABAP1 interaction could operate in a feedback control mechanism in which AIP10 can somehow transcriptionally regulate *ABAP1* mRNA levels and thus its targets. In addition, *CYCB1;1 and CYCB1;2,* marker genes for cell division, showed around 70% and 15% higher expression, respectively, in the knockout plants at 11 DAG (Fig 4A), endorsing the higher rates of cell division observed in *aip10-1*. In *aip10-2*, the expression profile in 11 DAG leaves was similar as that observed in *aip10-1* plants, except for *CDT1b* and *CYCB1;2* that did not show significant higher expression in the mutant (Fig. 4A). Since *aip10-1* growth phenotypes were more pronounced, compared with *aip10-2*, we can speculate there might be a minimum threshold level of reduction in *AIP10* expression, to activate all the mechanisms by which AIP10 silencing improves plant growth. In roots, the expression of marker genes exhibited the same pattern observed in shoot in Col-0 and *aip10-1* (Fig. S6A).

**Figure. 4.**
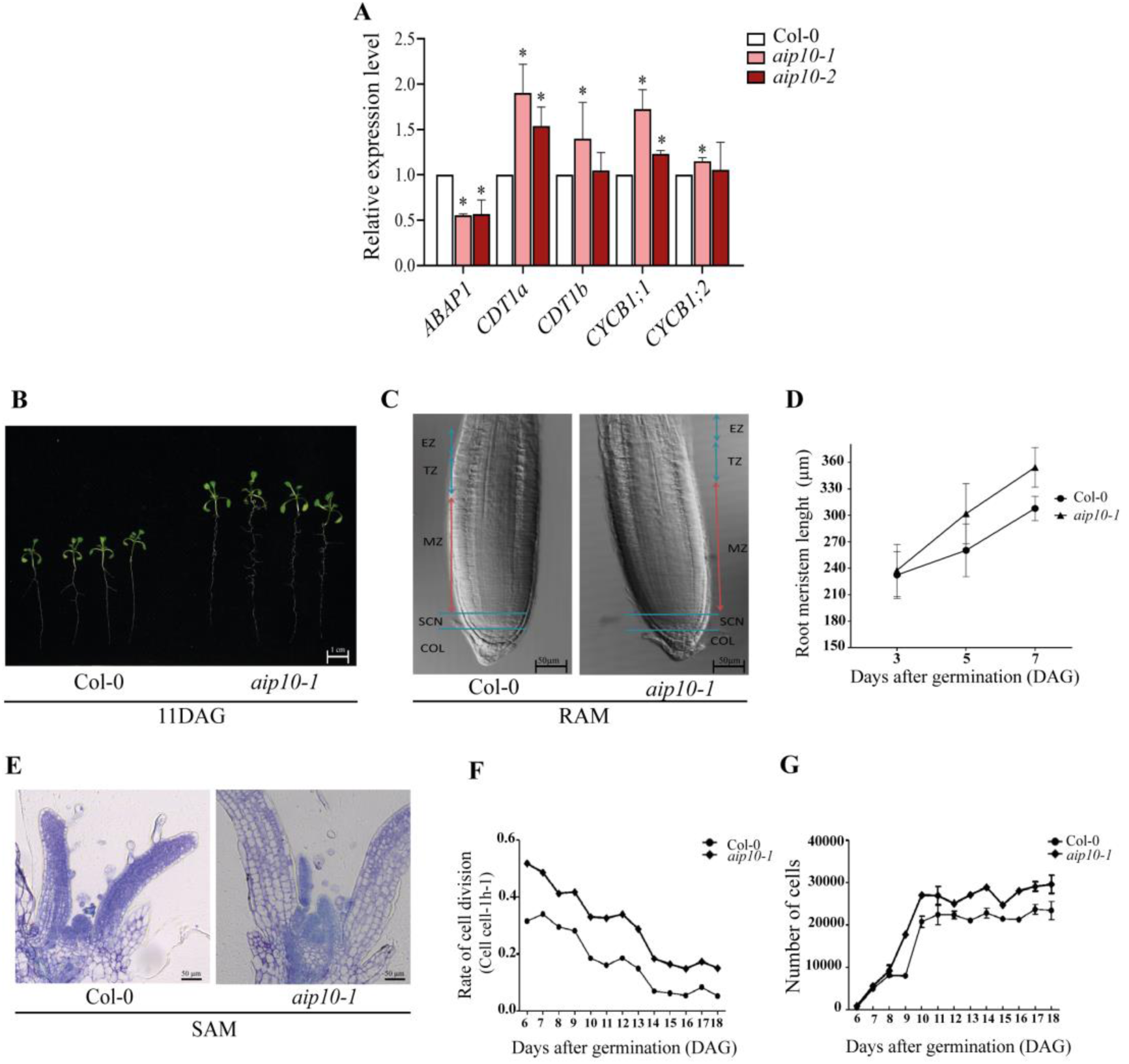
Cell division analyzes in *aip10-1, aip10-2* and wild-type Col-0 plants. **A)** Expression analysis of *ABAP1, CyclinB1;1*, *CyclinB1;2* and the ABAP1 target genes, *CDT1a* and *CDT1b*, in Col-0, *aip10-1* and *aip10-2* plants. Relative mRNA levels were evaluated in both genotypes in shoot of 11DAG seedlings grown *in vitro* in MS medium with 1% sucrose in a photoperiod of 12h/12h to 21°C. Each biological replicate (n = 3) consisted of a set of 10 plants. Data were normalized with the expression of *UBI14* and *GAPDH* as reference genes. Bars indicate standard deviation of biological replicates. Statistical analysis was performed using the Student’s t test (p<0.05). Asterisks (*) indicate significant differences between samples relative to Col-0. **B)** Seedlings at 11 DAG, grown *in vitro* in 1% sucrose MS medium. **C)** Bright-field differential interference contrast of Root Apical Meristem of *aip10-1* and Col-0 plants, 5 DAG (n=5). EZ: elongation zone; TZ: transition zone; MZ: meristematic zone; SCN: Stem cell niche; COL: columella. The arrows show the extension of meristems, 20x magnification. **D)** Mean RAM length of plants at 3, 5 and 7 DAG (n=10) grown *in vitro* in MS medium with 1% sucrose in a photoperiod of 12h/12h to 21°C. **E)** Longitudinal section of the SAM of Col-0 and *aip10-1* plants at 10 DAG. Bars: 0,12 mm. All images are representative of the replicates analyzed. **F, G)** Kinematic analysis of growth of the first leaf pair of Arabidopsis Col-0 and *aip10-1* plants, from 6 to 18 DAG (n=5); **F)** rate of cell division and **G)** number of cells. The line graph corresponds to the monitoring of individuals from 6 to 18 days. Error bars represent averages ± SD of 5 plants.

Next, we evaluated whether AIP10 could act together with ABAP1, modulating meristematic activities through assays using *aip10-1* plants. Anatomical analysis showed increased length of the main root and greater number of lateral roots (Fig. 4B, S6 B, C), and longer Root Meristems (RAM) (Fig. 4C, D) in *aip10-1* plants compared to Col-0. In the Shoot Apical Meristem (SAM), the first and second pair of leaves emerged earlier in *aip10-1* plants, compared to Col-0 (Fig. 4E). These data were corroborated by a kinematic assay of leaf development showing that higher rates of cell divisions were maintained for a longer period in *aip10-1* (Fig 4F), leading to larger leaves (Fig. S6D) with higher cell numbers (Fig. 4G), while cell sizes were the same in *aip10-1* and Col-0 (Fig. S6E). The data indicate that the improved early development of *aip10-1* plants resulted from the boost in cell divisions.

Histochemical analysis illustrated an increased staining in SAM and RAM of *pCDT1a::GUS* x *aip10-1* plants, compared to *pCDT1a::GUS* x Col-0 (control), suggesting a more active *CDT1* promoter (Fig; S7A, B). Also, DNA ploidy levels during vegetative development were assessed by flow cytometry. In both leaf and root, no significant differences in ploidy were observed between Col-0 and *aip10-1* (Fig. S8A-C), which demonstrates that there was no difference in the endoreduplication process in the mutant.

### *AIP10* silencing triggers a major transcriptional reprogramming in pathways with impact on plant primary metabolism and on the response to the environment

To further investigate genetic pathways that could be involved in AIP10 role in development, transcriptomes were generated with 11 DAG and 35 DAG samples of roots and shoots of Col-0 and *aip10-1* plants, that presented the most pronounced growth phenotypes. The selected time points reflected two important stages to investigate the increase in size and productivity of *aip10-1* plants. Differentially expressed genes (DEGs) identified in selected pathways were further investigated by RT-qPCR in both mutants.

Illumina sequencing of 8 libraries at 11 DAG yielded a total of 264.3 million reads and sequencing of 12 libraries at 35 DAG yielded ∼1779 million reads. Differential expression between *aip10-1* and Col-0 was compared, identifying 300 DEGs for 11 DAG roots (157 up/143 down), 323 DEGs for 11 DAG shoots (112 up/211 down), 537 DEGs for 35 DAG roots (223 up/314 down), and 514 DEGs for 35 DAG shoots (114 up/400 down) (Fig. 5A, B).

**Figure. 5.**
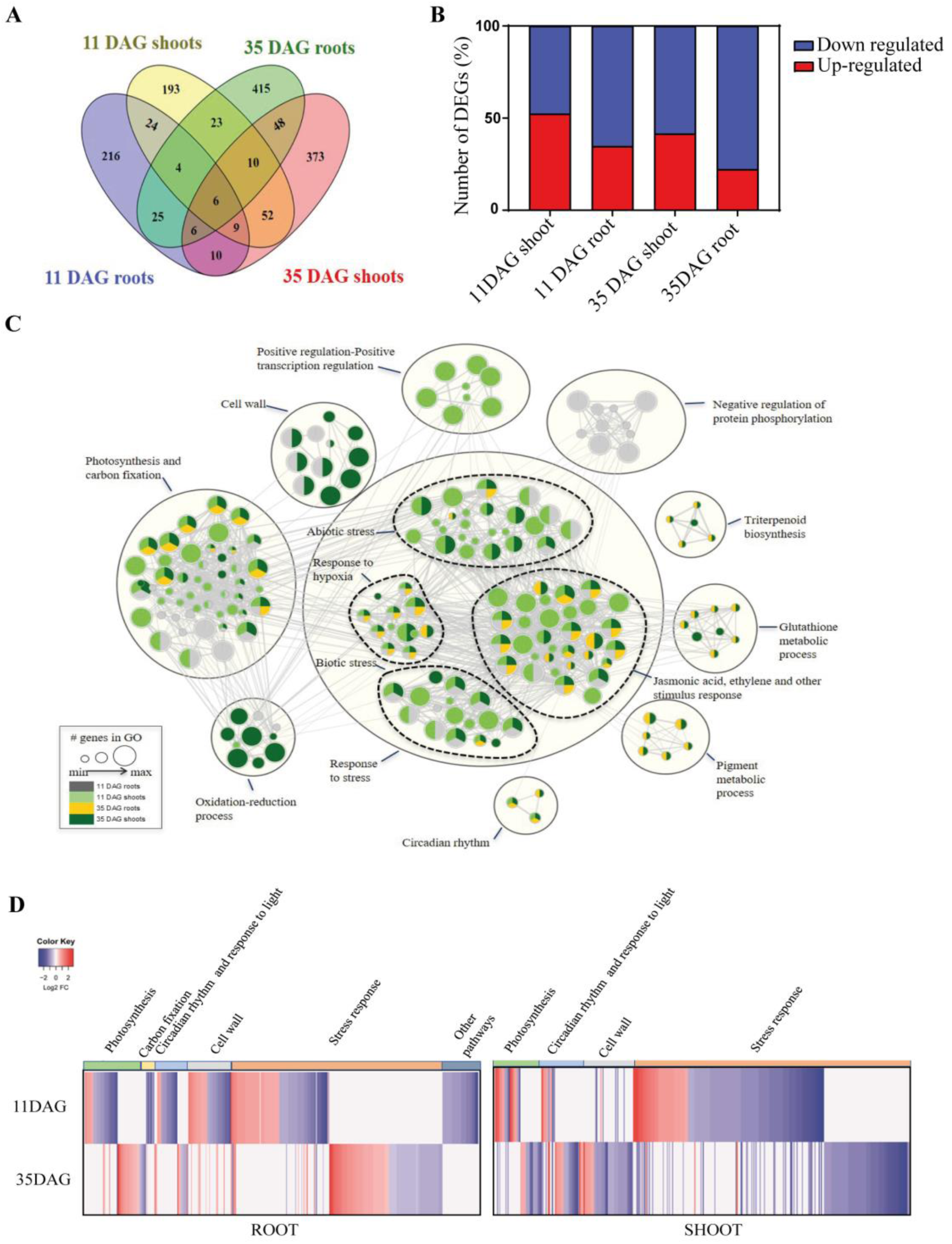
Gene expression reprogramming in the absence of *AIP10*. **A)** Venn diagram showing the total number of differentially expressed genes (DEGs) for each transcriptome comparison, considering *aip10-1* and Col-0 in shoot and root tissues at 11 DAG and 35 DAG from the website https://bioinformatics.psb.ugent.be/webtools/Venn/. **B)** Percentage of the number of DEGs upregulated and downregulated in each comparison: *aip10-1* and Col-0 in shoot and root tissues at 11 DAG and 35 DAG. Upregulated DEGs are shown in red and downregulated are shown in blue. **C)** Enrichment map (Cytoscape Enrichment map) displaying enriched pathways for each timepoint (11 DAG roots, grey; 11 DAG shoots, pale green; 35 DAG roots, yellow; 35 DAG shoots, dark green), regrouped in clusters. Nodes represent gene enrichment sets. Node size corresponds to the number of genes. Edges join gene sets that share common genes. **D)** Heatmap with main clusters for comparisons between *aip10-1* and Col-0 at 11 DAG and 35 DAG, in roots (left panel) and shoots (right panel). Colors represent log2-fold change (violet: downregulated and red: upregulated).

Gene-ranked pathway enrichment analysis with g:Profiler was performed to identify enriched functional categories of the differentially expressed genes (Fig. 5C). The Cytoscape map exhibits nodes that correspond to enrichment terms colored depending on their presence on each *aip10-1* transcriptome timepoint and tissue (11/35 DAG roots/shoots), while edges correspond to shared genes between pathways. Nodes were regrouped into larger clusters to identify major categories of enriched pathways. Photosynthesis and carbon fixation were the second major cluster, followed by cell wall, circadian rhythm and light response. Some clusters were only associated with one point in time, such as induction of transcription factors in 11 DAG root and the repression of protein phosphorylation in 11 DAG shoot. Furthermore, enriched oxidation-reduction process pathways were mainly present in 35 DAG shoot, and triterpenoid biosynthesis, glutathione metabolic process, and pigment metabolic process were DEGs only in 35DAG (Fig. 5C).

Heat maps for the most important clusters were generated and notable patterns were identified (Fig. 5D). Remarkably, the sets of genes linked to clusters, such as photosynthesis, carbon fixation, circadian rhythm, response to light, cell wall, and stress response, were distinct between 11 and 35 DAG, both in roots and shoots, suggesting that *aip10-1* plants differentially remodeled their transcriptional program between the two time points.

### Transcriptional reprogramming in *AIP10* silenced plants might involve SnRK1 regulatory pathway

The functional categories of DEGs in *aip10-1* transcriptomes showed an enrichment for clusters related with plant metabolism. As AIP10 can interact with KIN10 (Fig. 1D), a subunit of SnRK1, we investigated the involvement of this kinase in the transcriptional reprogramming profiles observed in *aip10 mutants*. First, we analyzed the gene expression profile in both mutants of known marker genes of SnRK1: *ASN1* (*ASPARAGINE SYNTHETASE* 1), *SEN1* (*SENESCENCE-ASSOCIATED 1*) and *TPS11* (*TREHALOSE PHOSPHATE SYNTHASE 11)*. Based on the downregulated profile of SnRK1 target genes at 35 DAG, at the transition to the reproductive phase when compared to Col-0 (Fig. 6B), the data suggest that SnRK1 was less active in both mutants at this timepoint. On the other hand, the upregulated profile of the target genes at 11 DAG, at earlier stages of development, suggests that SnRK1 was more active in *aip10-1* plants; however, the expression profile was not so clear in *aip10-2* plants (Fig. 6A).

**Figure. 6.**
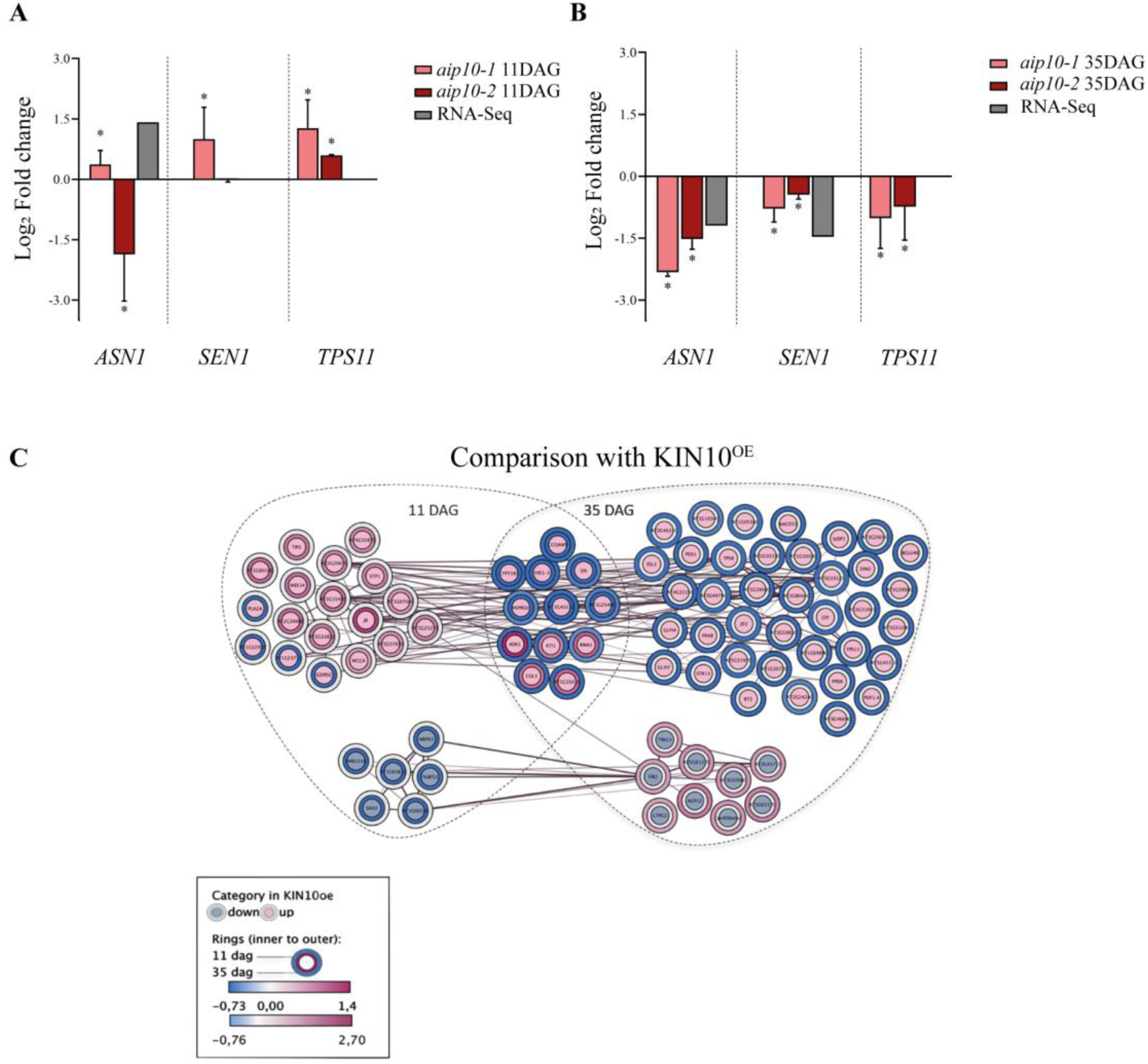
Expression profile analysis of KIN10 target genes in *aip10-1* transcriptomes to indirectly assess SNRK1 activity. **A, B)** Validation by RT-qPCR of selected SnRK1 (KIN10oe) marker genes differentially expressed (DEGs) in *aip10-1* shoot transcriptomes. Relative mRNA levels of *ASN1*, *SEN1* and *TPS11* genes in Col-0, *aip10- 1* and *aip10-2* plants were evaluated in both genotypes, in **A)** shoot of 11 DAG seedlings grown *in vitro* in MS medium with 1% sucrose in a photoperiod of 12h/12h to 21°C and **B)** shoot of 35 DAG plants grown directly in the soil in a photoperiod of 16h/8h to 21°C. Expression levels were normalized by the mRNA levels of the *UBI14* and *GAPDH* genes. Bars represent means ± SD (standard deviation) and * significantly different from Col-0 with p<0.05 with Student’s t test. Each biological replicate (n = 3) consisted of a set of 10 plants at 11 DAG and 5 plants at 35 DAG. The graph bars indicate each biological replicate in log2 fold change data, where value < 0 are repressed and > 0 are induced. **C)** Comparative expression profile analysis of genes (circles) regulated in both KIN10oe/sugar deprivation and *aip10-1* transcriptomes at 11 DAG and/or 35 DAG. Color of inner circles in the map (Cytoscape Omics Visualizer) indicates the expression profile of DEGs in KIN10oe (green: downregulated, magenta: upregulated). An expression heat map for DEGs in *aip10-1* is represented by colors in rings around the inner circle, indicating the log2 fold change (inner ring: 11 DAG and outer ring: 35 DAG). Edges join genes related to a common network or pathway. DAG, Days After Germination.

We further investigated SnRK1 activity in *aip10-1* plants, by comparing transcriptome profile of genes regulated by KIN10 overexpression (KIN10oe) under sugar starvation (Baena-González et al., 2007) with the profiles in 11 DAG and 35 DAG *aip10-1* transcriptomes. SnRK1 activity is increased during sugar starvation due to low levels of trehalose-6-phosphate (T6P), which is used as a signal to indicate a homeostatic sugar imbalance in plants (Baena-González and Lunn, 2020). Figure 6C presents a Map (Cytoscape Omics Visualizer) showing common DEGs between *aip10-1* and KIN10oe transcriptomes. 19 genes that were upregulated, and 6 genes that were downregulated by KIN10oe/sugar starvation, presented a similar differential expression pattern in *aip10-1* at 11DAG, suggesting increased SnRK1 activity at this developmental stage. Contrastingly, at 35 DAG, all 51 genes that were upregulated and 9 genes that were downregulated by KIN10oe/sugar starvation showed an opposite expression pattern in *aip10-1* plants, suggesting that SnRK1 was less active.

SnRK1 and TOR have been shown to exert their roles through large transcriptional reprogramming which affects regulation of nutrient-driven processes mostly in opposite ways (Baena-González et al., 2007; Dong et al., 2015). Therefore, other comparisons were performed between the expression profiles in transcriptomes that were generated: a) in conditions when SnRK1 is active (KINoe/sugar starvation); b) during inhibition of TOR activity (iTOR) (Dong et al., 2015); c) when AIP10 is silenced, at two developmental stages (Fig. S9A). Out of 84 DEGs identified as target genes of SnRK1 (Baena-González et al., 2007) in the *aip10-1* transcriptome, 60 DEGs were also transcriptional targets of TOR (Fig. S9A) (Dong et al., 2015). GO analysis revealed that these genes, common to the three comparisons mentioned above, present biological process terms related to metabolism of carbon and lipids, hypoxia, hormone and stimulus (Fig. S9 B-E). These TOR target genes have been identified through expression profiling analysis of 11 DAG seedlings treated for 24 hours with AZD8055 (AZD), an inhibitor of TOR activity (Dong et al., 2015).

Comparative analyzes between DEGs from the *aip10-1* transcriptomes and iTOR showed that among the set of 1583 genes upregulated by iTOR, 341 DEGs were common to *aip10-1* (Fig. S10). Most DEGs were downregulated in our 35 DAG mutant, suggesting that TOR is activated at this time. GO analysis showed the presence of genes associated with the response to endogenous and external stimuli, hormones and lipids (Fig. S11). From the transcriptional data we can speculate that SnRK1 was possibly more active in early stages of development in parallel with TOR in *aip10-1* at 11 DAG, while TOR appeared to predominate in the transition to the reproductive stage in the mutant when compared to control. However, further functional experiments would be necessary to confirm the involvement of TOR pathway in *aip10-1* plants.

### *AIP10* silencing increases the photosynthesis and CO2 assimilation efficiencies

Next, we further investigated the primary metabolism reprogramming modulated by AIP10 silencing, by combining the phenotyping and transcriptome data, with biochemical and metabolic analyses.

Photosynthesis belongs to the second largest cluster of pathways enriched in *aip10-1* plants (Fig. 5C). Most DEGs related to photosynthesis in 11 DAG shoot of *aip10- 1* were induced (Fig.7), such as genes from photosystems I and II (PSI and PSII), including seven *light-harvesting chlorophyll a/b binding* (*LHCB*) and *PsaD2*. These LHCB proteins serve as the photosystem II antenna complex (Jansson, 1999) and PsaD2 is a component of the PSI reaction center (Ihnatowicz et al., 2004). Three genes from the Chaperone domain *DnaJ* (DJ) superfamily, involved in the stabilization of PSII (Chiu et al., 2013), and *LZF1*, described as a transcriptional regulator that influences chloroplast biogenesis and function (Chang et al., 2008), were also induced. Furthermore, *NDA1*, a type II NAD(P)H dehydrogenase, was also induced and may have a role in stabilization and redox regulation in the mitochondrial respiratory chain (Wallström et al., 2014). These genes were analyzed through RT-qPCR in the mutants, validating their expression profile in plants with total and partial gene silencing (Fig. 8A).

**Figure. 7.**
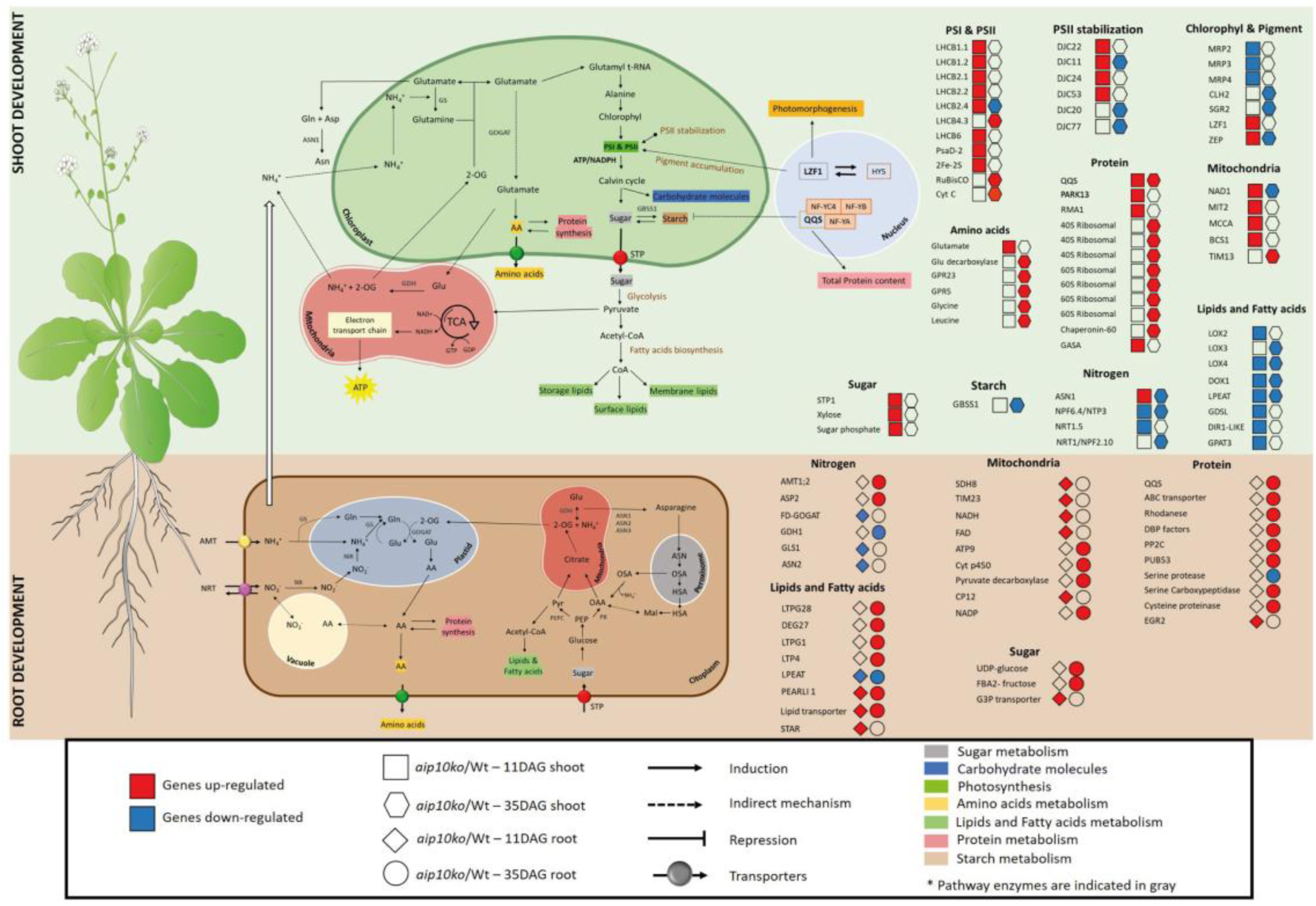
Main genes involved in energy metabolism differentially expressed in shoots and roots of *aip10-1* plants. The schematic representation depicts the expression pattern of genes involved in CO2 uptake and carbon metabolism, photosynthetic pigment synthesis and degradation, as well as genes related to nitrogen, protein, amino acids, starch and sucrose metabolism in both leaves and roots of *aip10-1* plants. The green background refers to the DEGs of the shoot and the brown refers to the DEGs of the root. DEGs in different transcriptomes are represented as follows: squares for 11 DAG shoot, hexagons for 35 DAG shoot, diamond for 11 DAG root and the circle for 35 DAG root. Red color refers to upregulated genes and blue color refers to downregulated genes. The icons present in the figure were highlighted in the footer of the figure. The main products of each route were highlighted with different colors as indicated in the figure. Gray: Sugar metabolism; Blue: Carbohydrate molecules; Fluorescent green: Photosynthesis; Yellow: Amino acid metabolism; Light green: Lipid and fatty acid metabolism; Pink: Proteins; Brown: Starch. DEGs, Differentially Expressed Genes; DAG, Days After Germination.

**Figure. 8.**
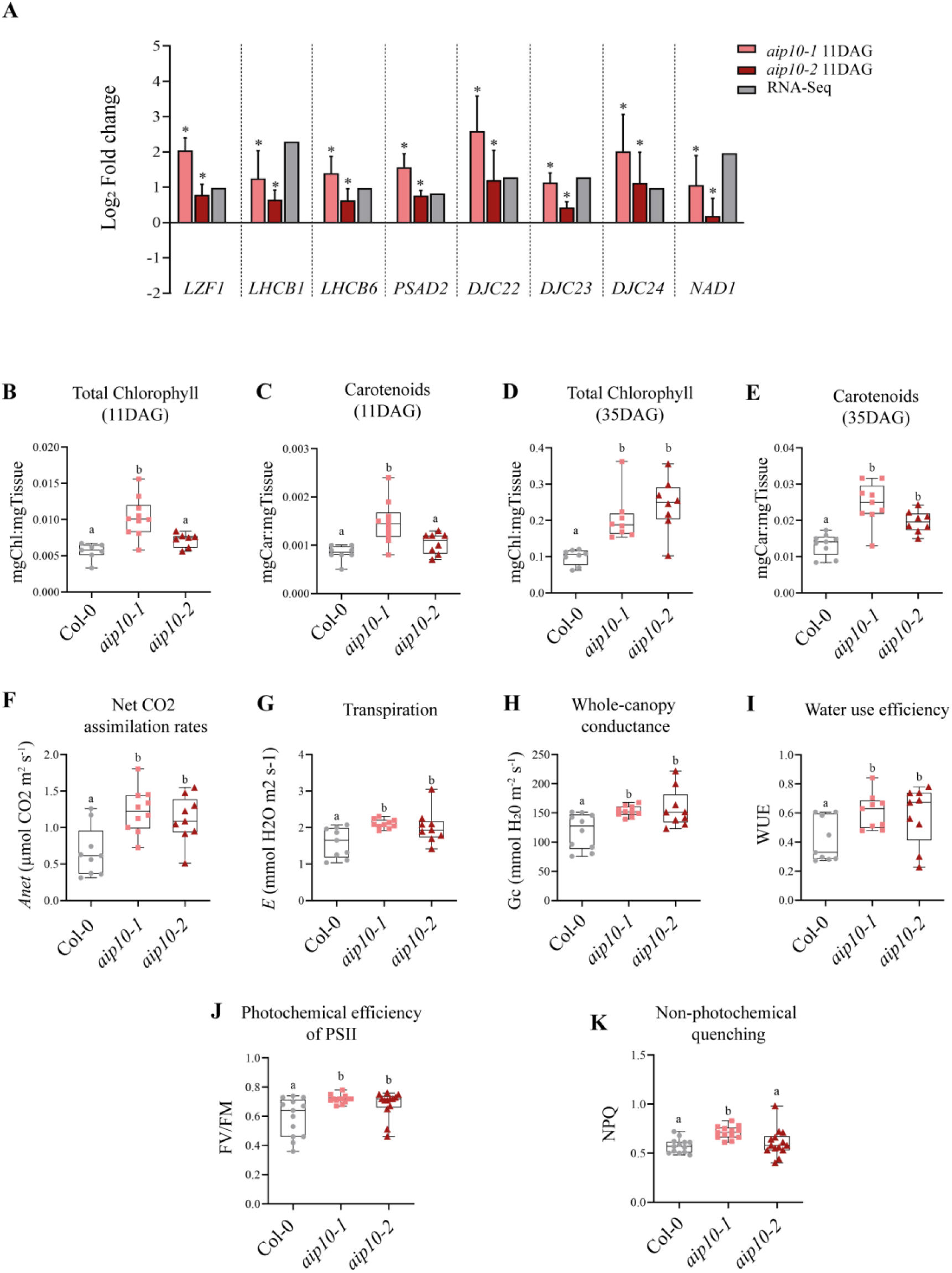
Analysis of photosynthetic genes, chlorophyll content and photosynthesis parameters in *aip10-1* and *aip10-2* plants. **A)** Relative mRNA levels of genes related to photosynthesis (*LZF1*, *LHCB1, LHCB6*, *PSAD2, DJC22, DJC23, DJC24* and *NAD1)* in shoot of 11 DAG seedlings of Col-0, *aip10-1* and *aip10-2* plants grown *in vitro* in MS medium with 1% sucrose in a photoperiod of 12h/12h to 21°C. Expression levels were normalized by the mRNA levels of the *UBI14* and *GAPDH* genes. Bars represent means ± SD (standard deviation) and * significantly different from Col-0 with p<0.05 with Student’s test. Each biological replicate (n = 3) consisted of a set of 10 plants at 11 DAG.RT-qPCR and RNAseq data are presented. The graph bars indicate each biological replicate in log2 fold change data, where value < 0 are repressed and > 0 are induced. **B-K**) Evaluation of photosynthetic physiological parameters. **B, D)** Total chlorophyll (*a*/*b*) and **C, E)** carotenoid content of Col-0, *aip10-1* and *aip10-2* plants grown directly in the soil in a photoperiod of 16h/8h to 21°C at **B,C)** 11 DAG and **D,E)** 35DAG (n=10). **F)** Net CO2 assimilation rate; **G)** Transpiration rate; **H)** Whole-canopy conductance. **I)** Water use efficiency (WUE); measurements were performed with the LICOR-6400 XT Instrument at 22°C in ambient CO2 at 400μmol of photons m^−2^s^-1^ (n=10). (J,K) Chlorophyll fluorescence parameters of Col-0, *aip10-1* and *aip10-2* plants at 11 DAG, including: **J**) Photochemical efficiency of PSII under dark adaptation (Fv/Fm), **K)** non- photochemical quenching (NPQ); measurements were performed with the Fluorcam 800 MF, Photon Systems Instruments (n=15). The box plot graphs show the distribution of data. The box represents the interquartile range (IQR), with the inner line running down the median. The whiskers extend to the maximum and minimum values, including all points within that range. The difference between groups in all analysis was confirmed by One-Way ANOVA (p < 0.05), and different letters indicate statistically different means according to Tukey’s test at 5% probability.

Differently from what was observed at 11 DAG, *aip10-1* plants at 35 DAG showed mainly repressed photosynthesis-related DEGs. This difference is possibly due to the formation of the photosynthetic apparatus, which still occurs at 11 DAG, a period in which most genes are being regulated. Together, these data suggest that silencing *AIP10* appears to improve the formation of the photosynthetic apparatus in the early stages of development, through reprogramming gene expression.

Consistent with RNA-seq data, *AIP10* silencing affected photosynthetic traits, showing a higher content of chlorophyll and carotenoids at 11 and 35 DAG in both mutants (Fig. 8 B-E), when compared to Col-0. Under greenhouse conditions, *aip10-1* and *aip10-2* at 25 DAG showed rates of up to 83% higher of CO2 uptake (*Anet*) (Fig. 8F) and higher transpiration rate (*E*) (Fig. 8G) which remained relatively lower than the gain obtained from photosynthesis. Added to this, *aip10* mutants showed higher stomatal conductance (Gc) (Fig. 8H) and water use efficiency (*WUE*) (Fig. 8I), indicating that these plants capture more CO2 and loose fewer water molecules at each stomatal opening. The measurement of photosystem II efficiency (Fv/Fm) showed that *aip10* mutants had greater photosystem II activity (Fig. 8J), in addition to greater energy dissipation in the form of heat (Fig. 8K).

### *AIP10* silencing modulates the flow of carbon fixation into biomass by improving contents of protein, lipids (triglycerides) and carbohydrates

Carbon fixation belonged to the second major cluster of enriched pathways in *aip10-1* plants (Fig. 5C). We observed an enrichment of pathways such as sugar, protein, starch, nitrogen and lipids (Fig. 7). Among the DEGs induced in *aip10-1* in11 DAG shoot was *STP1* (*Sugar Transporter Protein 1*), which encodes an H+/monosaccharide cotransporter. At 35 DAG, ribosomal genes were induced indicating continuous protein production throughout development. Some lipid-related genes were mainly induced in 11 DAG root and repressed in 11 DAG and 35DAG shoot. Furthermore, *QQS* (*Qua-Quine Starch*), an *A. thaliana* orphan gene that acts as a regulator of starch biosynthesis was upregulated in *aip10-1* at both timepoints. Its overexpression in soybean triggered increased protein content and decreased starch levels (Li and Wurtele, 2015). Besides, there was repression of the *GBSS1* gene (G*ranule-Bound Starch Synthase 1*) involved in starch accumulation, encoding a protein from the UDP-Glycosyltransferase superfamily that is responsible for the biosynthesis of amylose in plants. The data suggest that starch was re-directed to the synthesis of other metabolites in *aip10-1*. The gene expression profiles were validated through RT-qPCR (Fig. 9A, B) in the mutants.

**Figure. 9.**
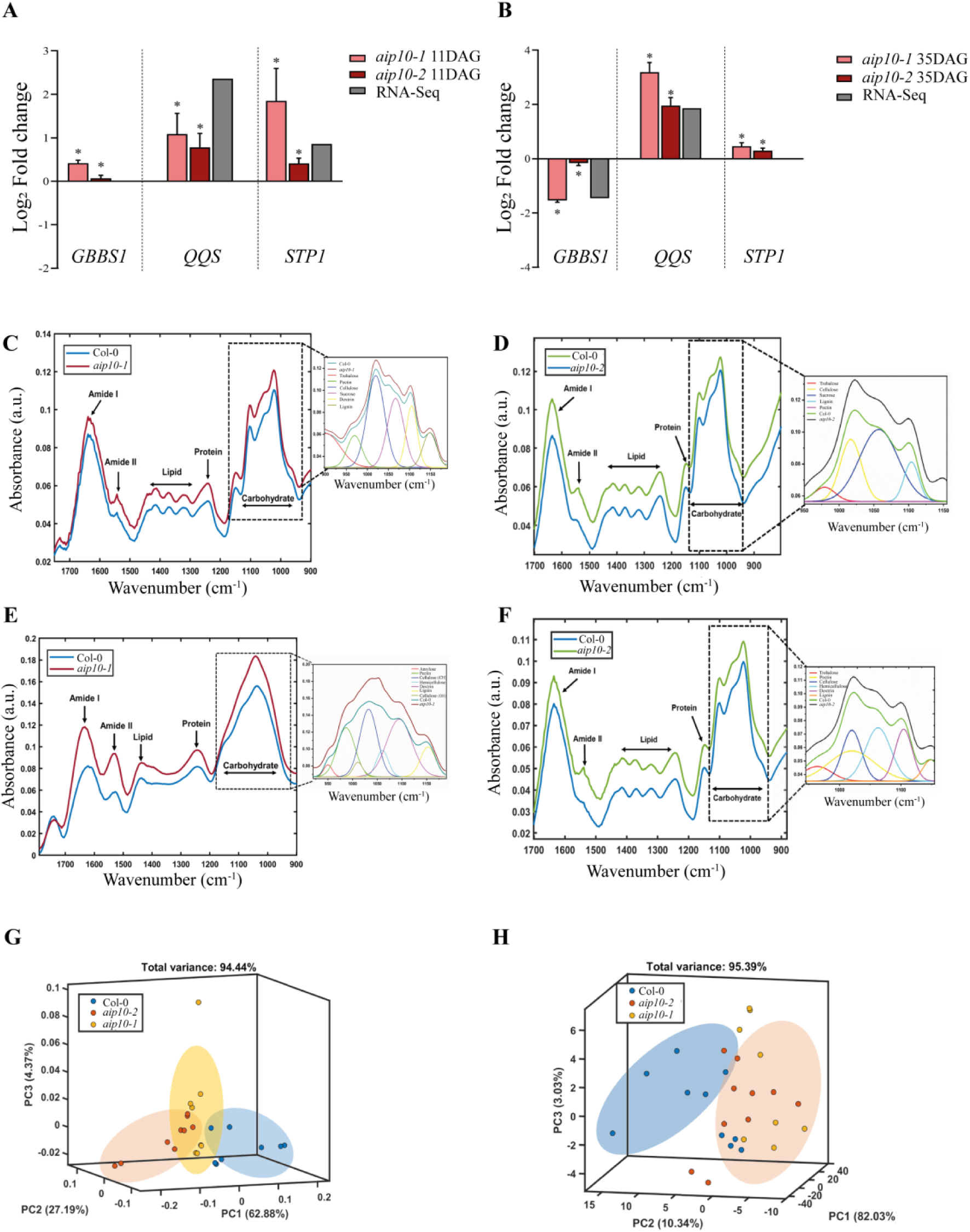
Metabolic profiles of *aip10-1* and *aip10-2* plants. **A-B)** Relative mRNA levels of genes involved in carbon metabolism, differentially expressed in the *aip10-1* transcriptome (*GBSS1*, *QQS* and *STP1* genes), in Col-0, *aip10-1* and *aip10-2* plants. Expression levels were evaluated in both genotypes in **A)** shoot of 11 DAG seedlings grown *in vitro* in MS medium with 1% sucrose in a 12h/12h photoperiod at 21°C and **B)** shoot of35 DAG plants cultivated directly into the soil in a photoperiod of 16h/8h 21°C. Expression levels were normalized by the mRNA levels of the *UBI14* and *GAPDH* genes. Bars represent means ± SD (standard deviation) and * significantly different from Col-0 with p<0.05 with Student’s t test. Each biological replicate (n = 3) consisted of a set of 10 plants at 11 DAG and 5 plants at 35 DAG. RT-qPCR and RNAseq data are presented. The graph bars indicate each biological replicate in the log2 fold change data, where value <0 are repressed and >0 are induced. **C-F)** Metabolic profiling of 35 DAG leaves and mature seeds from Col-0, *aip10-1* and *aip10-2* plants grown directly in soil in a photoperiod of 16h/8h 21°C, by using a Shimadzu IRPrestige-21 attenuated total reflectance infrared spectrometer (ATR-FTIR). Data show average spectra obtained by the ATR-FTIR technique in the region of 900–1700 cm^−1^ which correspond to the regions of most sensitive absorption of the main components. **C)** 35 DAG leaves of Col-0 and *aip10-1* plants. **D)** 35 DAG leaves of Col-0 and *aip10-2* plants. The experiment was carried out on 2 different leaves from 4 individuals of each genotype. The assigned values are normalized by the area of the diamond crystal plate. **E)** Mature seeds obtained from Col-0 and *aip10-1* plants. **F)** Mature seeds obtained from Col-0 and *aip10-2* plants. The experiment was carried out in 3 different macerated pools of 100 seeds each. The assigned values are normalized by the area of the diamond crystal plate. **G, H)** Score plot for the three principal components applied to the seed ATR-FTIR dataset of Col, *aip10-1* and *aip10-2* plants respectively.

The metabolite levels were measured in *aip10* mutants, using ATR-FTIR spectra, that refers to the vibrations between the chemical bonds that can be assigned to distinguish specific functional groups at each wavelength variation (Talari et al., 2017) (Table S4). Within the acquisition range (500 - 4000 cm^-1^), a region called spectral bio-fingerprint was established between 900 – 1800 cm^-1^ (S12A, B; S13D), responsible for providing a large amount of information about the chemical groups present in biomolecules both in leaves and seeds (Santos et al., 2020). The most distinct band regions demonstrated that leaves of *aip10* mutants at 20 DAG (Fig S13A, B) and mature leaves at 35 DAG (Fig. 9C, D) showed approximately 19% higher abundance of proteins, compared to Col-0 plants. The mutants presented higher content of lipids and carbohydrates at 20 DAG (Fig S13A, B) and 35 DAG (Fig. 9C, D). The increased metabolic content in the leaves of the *aip10* mutants might be contributing to its greater biomass. Seeds of mutants also presented higher metabolic contents (Fig. 9E, F), showing a gain of approximately 40% in proteins, 14% in lipids and 18% in carbohydrates (triglycerides). The mutants presented higher levels of reserve polysaccharides. The PC model demonstrated that the mutants cluster together, showing their metabolic similarity in both tissues, that differ from Col-0 (Fig.9G, H; Fig. S13C).

## Discussion

Plant development is shaped by a series of environmental factors, driving to adaptive forms for their survival. In this work, we investigated a protein belonging to the ABAP1 regulatory pathway, called AIP10, which integrates the modulation of the plant’s cell cycle and primary metabolism. Plants with total and partial AIP10 silencing activated cell divisions, leading to organs with increased cell numbers and biomass, that is possibly supported by the transcriptional reprogramming that improved primary metabolism, photosynthesis, and carbon fixation.

### AIP10 is a member of the ABAP1 regulatory network, that interacts with ABAP1 and KIN10, a subunit of SnRK1 kinase

Plant growth is the result of cell division rates and also the balance between cell proliferation, expansion and differentiation (Jones et al., 2017). ABAP1 and the proteins with which it interacts (AIPs) constitute a plant specific pathway for regulating plant cell cycle progression at G1/S transition, an important phase modulated by environmental signals (Masuda et al., 2008). ABAP1 contains eight predicted repeats of the Armadillo β-catenin (ARM) domain in its N-terminal portion, critical for protein-protein interactions (Zhou et al., 2015), and a BTB/POZ (Broad complex/Tram-Track/Bric-a-brac/Poxyvirus and zinc finger) in its C-terminus. Due to the diverse repeats of the ARM domain, ABAP1 can associate with different proteins allowing multiple interactions to occur simultaneously to form different complexes (Masuda et al., 2008).

The characterization of different ABAP1 interactors provided evidence that ABAP1 performs different functions within the cell cycle. When interacting with TCP24, ABAP1 is capable of binding to the CDT1a/b promoters, repressing the expression of both genes (Masuda et al., 2008). The ABAP1 interactor (AIP1), which contains an Agenet/Tudor domain, also interacts with unmodified histones and with LHP1 (LIKE HETEROCHROMATIN PROTEIN 1), being involved in chromatin compaction remodeling, and gene silencing, that regulate flowering through FLT repression (Brasil et al., 2015). ABAP1 also regulates the differentiation of male and female gametophytes, by forming a complex with AtTCP16 in male gametogenesis, and with ADAP in female gametogenesis, which repress the transcription of its target genes, CDT1b and EDA24, respectively (Cabral et al., 2021).

Here we characterized an ABAP1 interactor, AIP10, and provided data on its dual role in regulating cell divisions and in transcriptional reprogramming, through its protein interactions with ABAP1 and KIN10. Our data showed several evidence supporting a joint role of AIP10 and ABAP1, as negative regulators of proliferative cell divisions in all plant meristems. All four AIP10 isoforms were expressed throughout development, indicating a potential role in regulating DNA replication and proliferative cell divisions in various plant meristems. AIP10 directly interacted with ABAP1 and they were found in a complex *in vivo*. AIP10, similarly as ABAP1, was exclusively localized in the nuclei, and both proteins were present in meristems in cycling cells, being absent in mitotic cells (Masuda et al., 2008). AIP10 role in regulating cell division could occur through direct interaction with ABAP1, but also through a feedback mechanism of transcriptional control in which AIP10 could somehow transcriptionally regulate ABAP1 mRNA levels, since AIP10 silenced plants showed reduced ABAP1 mRNA levels. Accordingly, the mRNA levels of two direct targets of ABAP1 repression at G1/S transition, *CDT1a* and *CDT1b* genes, were upregulated in AIP10 silenced plants. Finally, a joint role of AIP10 with ABAP1 could enable a negative regulation of the pre-RC, repressing DNA replication and proliferative cell division. This hypothesis was supported by the kinematics data showing that AIP10 silencing increased rates of cell division and cell numbers in leaves; increased lateral root numbers in roots; and raised expression of cell division marker genes, both in shoots and roots.

We also showed that AIP10 physically interacted with KIN10, but not with KIN11, the catalytic subunits of SnRK1, a metabolic regulator involved in stress responses (Peixoto and Baena-González, 2022). Homologs of AIP10 in bryophyte *P. patens* (SKI1 and SKI2) and rice (HDR1) also showed interaction with proteins homologous to the catalytic subunit of yeast Snf1 kinase and plant SnRK1 (Thelander et al., 2007; Sun et al., 2016). The AIP10.1, AIP10.2 and AIP10.3 isoforms of AIP10 have in the C-terminal portion a nuclear localization site and a putative phosphorylation site initially identified in the two proteins that interact with SnRK1 homologs. In this same region, the AIP10.1 and AIP10.3 isoforms have a coiled-coil conformation that has previously been implicated in interactions with SnRK2s in *Medicago truncatula* and is present in SnRK1a-interacting negative regulators (SKINs) in rice (Nolan et al., 2006; Lin et al., 2014) Furthermore, it was reported that AIP10 can interact with Val2 (VIVIPAROUS- 1/ABSCISIC ACID INSENSITIVE 3-LIKE 2) (Altmann et al., 2020), a transcriptional repressor that recruits the Polycomb repressive complex 2 (PRC2) (Yuan et al., 2021) to catalyze H3 Lys27 trimethylation (H3K27me3) in some of its targets, regulating the expression of development and flowering genes (Gan et al., 2015). These data strengthen that AIP10 has a role in transcriptional regulation, whether at the genetic or epigenetic level. Remarkably, VAL2, as well as AIP1, were shown to interact with LHP1, to regulate flowering (Brasil et al., 2015; Yuan et al., 2021), suggesting a possible connection of these proteins and the ABAP1 network. Also, an epigenetic role of SnRK1 was shown in rice, as the kinase stimulates JMJ705, an H3K27me3 demethylase, to control energy homeostasis (Wang et al., 2021). Altogether, the data support the hypothesis that AIP10 acts in a transcriptional reprogramming hub, in specific developmental contexts.

### AIP10 is involved in integrating cell cycle, transcriptional reprogramming and primary metabolism, modulating plant development

The absence of AIP10 promoted vegetative growth, through increased cell divisions. In addition to the expansion of meristems, the greater biomass of knockout plants was a consequence of the production of more organs, at an accelerated pace and for a longer period of time throughout the plant’s life cycle. The cell cycle is an energetically consuming process dependent on availability of energy metabolites. AIP10 interacts with SnRK1, a central metabolic regulator (Baena-González et al., 2007; Baena-González and Lunn, 2020) that acts in a complex network that includes TOR, a master regulator of development (Margalha et al., 2019) Several genes regulated by the absence of AIP10 converged with the gene expression profiles regulated by SnRK1 and TOR pathways that share partially antagonistic regulatory activities (Margalha et al., 2019). Therefore, we can speculate that the transcriptional reprogramming observed in AIP10 silenced plants could be mediated, at least in part, by SnRK1 and TOR activities. The transcriptional profile data suggested that in the absence of AIP10, TOR was predominantly more active throughout development, thus favoring plant growth, in line with the *aip10-1* phenotypes observed. Accordingly, SnRK1 seemed to be less active in *aip10-1* and *aip10-2* mature plants. However, in young *aip10-1* seedlings, there was evidence of increased SnRK1 activity due to the induction of marker genes, which was not as explicit in *aip10-2*. An increase in SnRK1 levels and activity at the beginning of germination has already been demonstrated (Lu et al., 2007). Even under favorable conditions, fluctuations in sugar levels during seedling establishment activate SnRK1 (Henninger et al., 2022) due to the need for greater energy supply and sugar homeostasis (Peixoto and Baena-González, 2022). Thus, SnRK1 could modify developmental programs according to metabolic state, to adjust plant growth to a specific environment (Margalha et al., 2019). SnRK1 activates photosynthetic genes and regulates growth in response to carbon (Zhang et al., 2009). Therefore, the expression profile of genes related to photosynthetic efficiency in AIP10 silenced plants could be mediated by SnRK1 activities. In parallel, TOR signaling pathway can activate meristems and cell division by controlling E2Fa regulators, while its inhibition by SnRK1 hinders cell cycle progression (Sablowski, 2016). Thus, we can also speculate that activated TOR, in the absence of AIP10, could be contributing to the higher cell division rates in *aip10-1*.

### Implications of the emergence and conservation of AIP10 in the Plantae kingdom and its high potential for biotechnological use in plant adaptation to the environment

Our data showed that silencing a single gene was able to stimulate growth through cell division and transcriptional reprogramming, leading to improved CO2 assimilation, increased biomass and higher metabolic content. A proposed model of the mechanisms that might be acting in the absence of AIP10 is presented in Fig. 10. AIP10 might act in a central hub that links the regulation of cell cycle with the plant primary metabolism, modulating plant development, by association with ABAP1 and SnRK1 (through KIN10) and, possibly, other proteins. AIP10 silencing leads to a transcription reprogramming in different plant pathways. In the absence of AIP10, ABAP1 levels and activity are reduced, releasing transcription repression of pre-RC genes, which allows pre-RC assembly that licenses DNA replication and cell cycle progression, culminating with increased rates of cell division. In parallel, the absence of AIP10 modulates SnRK1 activities, triggering a transcriptional reprogramming of primary plant metabolism, that regulates two major pathways: (i) promotes cell divisions by providing energy, and possibly by stimulating DNA replication mediated by TOR activation; (ii) leads to an increase in photosynthetic efficiency. A coupled regulation of increased cell division rates and carbon sequestration allows a more efficient carbon fixation into root and shoot biomass. Also, absence of AIP10 improves metabolites accumulation, serving as large energy reserve during the vegetative phase, that is metabolized at reproductive phase, contributing to the increase in fruit and seed production, with higher nutritional values.

**Figure. 10.**
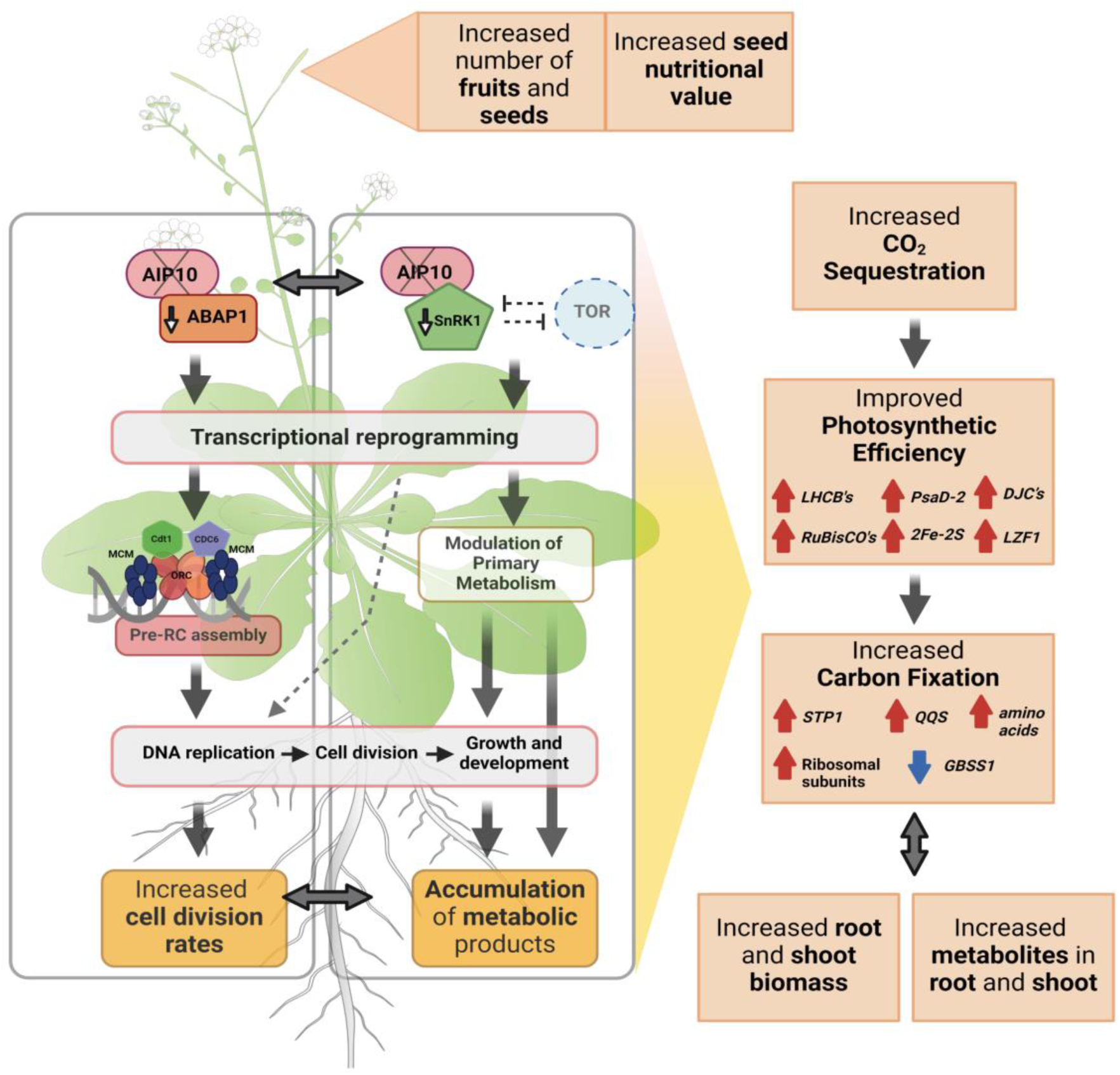
A hypothetical model of mechanisms triggered by the absence of *AIP10*. AIP10 might act by modulating plant development through interaction with ABAP1 and SnRK1 (KIN10), two key regulators of the cell cycle and primary metabolism, respectively (see text for details). In the absence of AIP10, ABAP1 transcriptional levels decrease, preventing inhibition of its targets, *CDT1a/b*, which allows the formation of the pre-RC, licensing DNA to replicate, and cells to progress through the cycle. In parallel, the absence of AIP10 triggers a transcriptional reprogramming of targets of SnRK1, and possibly of TOR pathways, modulating cell division and plant metabolism. These processes lead to an increase in photosynthetic efficiency and results in greater carbon fixation enhanced by an increase in the number of cells, and in the increased accumulation of metabolites. A combination of all these effects contribute not only to real increase in root and shoot biomass and in seed productivity, but also to improved nutritional value of plants, and increased energy source to sustain higher cell division rates. Created with BioRender.com.

Remarkably, plants with full AIP10 silencing show better improvements over plants with partial silencing. This raises the intriguing question of why the absence of AIP10 in *A. thaliana* mutants resulted in promotion of growth and metabolism. *AIP10* gene is found exclusively in the Plantae kingdom, and homologues were identified in all species with sequenced genome available. Previous studies have assigned AIP10 to a set of plant-specific genes that have only one copy in *A. thaliana* and *O. sativa*, likely originated in the chloroplast genome and being transferred horizontally to undergo stronger selection pressure, with a high probability of a critical function that promotes its conservation (Armisén et al., 2008). We can hypothesize that AIP10 is one piece in a larger gene network that might operate with gene expression homeostasis between its members, to fine tune the modulation of plant responses and adaptation to the environment.

Many genes and regulatory pathways evolved in the transition of plants to the terrestrial environment, allowing adaptations to water availability (Fürst-Jansen et al., 2022), such as the emergence of stomata, optimizing the absorption of CO2 and loss of water, and the evolution of the cuticle acting as an extracellular hydrophobic barrier. The *aip10-1* mutant showed several features that suggest greater adaptation to environmental fluctuations as a result of the conquest of the terrestrial environment, such as increased CO2 assimilation and decreased transpiration, enabling better use of water.

Our data demonstrated that modulation of a central hub that integrates the regulation of cell division rates with the plant primary metabolism was an efficient strategy to improve carbon sequestration and fixation into root and shoot biomass, leading to a real increase in plant productivity and nutritional value.

## Materials and Methods

### Plant material and growth conditions

Columbia (Col-0), *aip10-1* (SALK_022332), *aip10-2* (SALK_094618) and *pAIP10*::AIP10-YFP were obtained from the Arabidopsis Biological Resource Center (ABRC) collection. The *pAIP10::*AIP10-YFP line (Tian et al., 2004) harbors a genomic construct of the complete *AIP10* gene (including a 3Kb promoter/5’ UTR region and 1Kb 3’ UTR) encoding an AIP10 protein with C-terminal -YFP fusion.

Lines were confirmed by PCR genotyping using specific primers (Table S1). The *A. thaliana* seeds of *pCDT1a:*:GUS were kindly provided by Dr. Crisanto Gutierrez.

Seeds were sterilized according to Cabral et al., 2021. After 15 days, *in vitro* plants were transferred to a mixture of soil:vermiculite (3:1) and watered with NPK 20-20-20 fertilizer with a final concentration of 0. 6 g/L and cultivated at 22°C with 100 μmol photons m^-2^ s^-1^ light intensity, in 16h light/8h dark cycle for seed formation. Both the mutant and Col-0 (control) plants were cultivated in the same condition and under the same fertilizer treatment. The rosettes were weighed immediately after collection to evaluate fresh biomass. The dried biomass was obtained by drying the material in an oven at 30°C for 7 days.

### *In vitro* and *in vivo* protein interaction assay

The construct pDEST15.ABAP1 (Masuda et al., 2008) was used for expression of ABAP1-GST in *E. coli* BL21 cells, as described by (Chekanova et al., 2000) with some modifications. GST pull-down analyzes were performed following a protocol described by (Tarun & Sachs, 1996). Immunoprecipitation assays and protein gel blots were performed according to the protocols described to Masuda et al. (2008) with antibodies against ABAP1 (1:1000; Covance Corp.) or YFP (anti-GFP, 1:2000; Invitrogen) in blocking buffer. Detection was carried out according to the ECL Western Blotting System according to manufactureŕs instructions (GE-Healthcare). Anti-ABAP1 polyclonal antibody was developed against the peptide antigen GAPIVTQLID (amino acids 28 to 37) by Covance Corp.

### Yeast two-hybrid assay

The yeast two-hybrid interaction assay was performed by using strain PJ694-a (MATa trp1-901 leu2-3 112 ura3-52 his3-200 gal4Δ gal80Δ LYS2::GAL1-HIS3 GAL2-ADE2 met2::GAL7-lacZ). The ABAP1 regulatory network proteins tested in this article were obtained from the laboratory’s clone bank. KIN10 vectors were purchased from Clones/DNA Resources-Other BAC Libraries - TAIR. For the construction of KIN11(A11), its cDNA was amplified by PCR (Suppl. Table 1).

### Microscopy and kinematic analysis

The material was collected at different stages of development and processed according to Cabral et al. (2021). The methodology of kinematics was used to analyze leaf growth of wild type Col-0 and *aip10-1* plants and observed by DIC and confocal microscopy.

Histochemical detection of GUS activity in 5-day-old homozygous pCDT1a::GUS lines, in WT Col-0 and *aip10-1* backgrounds, was done with 5-bromo-4-chloro-3-indolyl β-D-glucuronide according to Cabral et al. (2021). The images were generated in a confocal microscope Leica TCS-SPE, using YFP filters with an excitation laser of 405 nm and the fluorescence was collected between 431 and 532 nm.

### Expression analysis by RT-qPCR

Total RNA was extracted according to Logemann et al., 1987 and treated with RNAse- free DNase I (New England Biolabs®). cDNA was performed using SuperScript™ III Reverse Transcriptase (Invitrogen) and the SYBR Green PCR Master Mix kit (Applied Biosystems) was used for expression analysis. Normalization was done against the average of the reference genes, *Ubiquitin 14* and *GAPDH* using the linear scaling method. The sequences of primers used in the RT-qPCR experiments are listed in Table S1.

### Flow cytometry analysis

Samples consisted of the first pair of leaves at 14 DAG and 20 DAG that were placed in Petri dishes containing 400μl of buffer (hypotonic buffer containing a non-ionic detergent) and chopped with a razor blade to release nuclei. Each macerate was resuspended in the buffer and filtered through a 50 μM filter. Nuclei were incubated in the dark with 1 mg/mL DAPI (4’,6-diamidino-2-phenylindole), and subsequently analyzed with a BRYTE HS Flow Cytometer (Bio-Rad). Analyzes were performed in biological triplicates.

### Photosynthetic analysis

To analyze photosynthetic capacity, the Fluorcam 800 MF, Photon Systems and LI-COR 6400XT equipment (LI-COR Biosciences, USA) were used. To record CO2 assimilation in response to photosynthetic light, measurements were performed in the morning, between 8:00 am and 10:00 am, adjusting the CO2 concentration to 400 ppm, and the photon flux density (PPFD) at 100 μmol m^-2^ s^-1^, with leaf temperature maintained between 21 to 22°C. The flux of CO2 and H2O was converted into photosynthesis (Anet) μmol CO2 m^−2^ s−1 and transpiration (E) mmol H2O m^−2^ s^−1^. Water use efficiency (WUE) was calculated as Anet/E (μmol CO2 mmol H2O^−1^). Canopy conductance (gc) was estimated using the formula gc = (E/DPV) × 101.325.

### Metabolic profiling via ATR-FTIR spectra acquisition

The metabolic profile spectra were obtained using a Shimadzu IRPrestige-21 total attenuated reflectance infrared spectrometer (ATR-FTIR) with a diamond crystal plate (IRIS module – PIKE Technologies). The results were normalized using the crystal surface of the ATR accessory. A pool of 100 macerated seeds was used, enough to cover the entire surface of the crystal of the ATR accessory and the leaves, two distinct regions were divided, and measurements were taken to obtain a spectrum representing the entire sample and three different biological replicates. All data processing was performed in an internal computational routine implemented in Matlab 2021a® (The MathWorks Inc., Natick, USA). The second derivative was also applied and tested with Savitz-Golay smoothing using a second-order polynomial and a 7-point window, which removes not only simple additive shifts but also first-order effects such as baseline shift and spectral noise.

### RNAseq and transcriptome analysis

The RNAseq libraries were constructed with total RNA from two shoot and root repeats of 11 DAG and three shoot and root repeats of 35 DAG by Fasteris Life Sciences SA. 8 libraries (11 DAG) and 12 libraries (35 DAG) were prepared according to the protocol available on the website and sequenced on the Illumina HISeq2500 (11 DAG) (single-end) and NovaSeq 6000 (35 DAG) (paired-end). The raw reads were subjected to control quality (Q30 > 80%) and adapter trimming using Trimmomatic software. Mapping was performed using the TAIR10 genome as a reference using Bowtie software. Expression estimation, normalization and differential expression was performed using Cufflinks software. Libraries were normalized using the FPKM method with significance values at p-value <0.05. The differentially expressed genes (DEGs) were compared between the libraries, based on the difference in expression, generating FoldChange (FC) values classified as induced or repressed based on the Log calculation in base 2 of the FC values comparing the FPKM values of the library of interest against the control. Pathway enrichment analysis was performed using g:Profiler and maps were visualized with EnrichmentMap in the Cytoskape software.

### Sequence alignment and phylogenetic analysis

Multiple protein sequence alignment was performed from available database sequences from species showed in Table S2 and S3 was conducted using CLUSTALW with default parameters. Subsequently, phylogenetic trees were generated using the IQTREE software through Maximum Likelihood Analysis, employing the JTT + Invar (I) + Gamma (G) model with 4 categories and subjected to 1000 repetitions.

### Statistical analysis

Statistical analysis was used following the number of variables. An unpaired Student’s t-test was performed between two samples (* p < 0,05). Tukey’s one-way analysis of variance was used to test the significance level of multiple groups and using P < 0.05 was considered to indicate statistical significance represented in letters.

### Accession numbers

Sequence data from this article can be found in the GenBank/EMBL data libraries under accession numbers SRR27777879 - SRR27777886 (SUB14180857) and SRR27764739 - SRR27764750 (SUB14179990)._The datasets generated for this study can be found in NCBI platform with bioproject number PRJNA1070685.

## Supplementary Data

Supplemental Figure S1. Protein interaction screening by yeast two-hybrid to identify potential AIP10 binding partners.

Supplemental Figure S2. Expression pattern of AIP10 isoforms in different *A. thaliana* tissues.

Supplemental Figure S3. Phylogenetic analysis of AIP10 and its putative orthologs.

Supplemental Figure S4 Phylogenetic analysis of ABAP1 and its putative orthologs.

Supplemental Figure S5 Molecular characterization of *AIP10* mutant lines.

Supplemental Figure S6. Analysis of root and leaf growth parameters in *aip10-1*, compared with wild-type Col-0.

Supplemental Figure S7. Analysis of *CDT1a* expression in meristems of *aip10-1* plants compared to wild-type Col-0.

Supplemental Figure S8. Average nuclear DNA content analyzed by flow cytometry.

Supplemental Figure S9 Comparative analysis of the expression profile of genes differentially expressed in both KIN10^oe^ and *aip10-1*.

Supplemental Figure S10. Comparative analysis of the expression profile of genes regulated by TOR inhibition differentially expressed in *aip10-1*.

Supplemental Figure S11. Functional categorization of genes regulated in both iTOR and *aip10-1*.

Supplemental Figure S12. Loading chart for the first two main components applied to the ATR-FTIR dataset.

Supplemental Figure S13. Metabolic analysis by ATR-FTIR spectrum of *aip10-1* and *aip10-2* plants at 20 DAG, compared with wild-type Col-0.

**Supplemental Table S1** Primers used in this study.

**Supplemental Table S2** AIP10 putative orthologs in other plant species.

**Supplemental Table S3** ABAP1 putative orthologs in other plant species.

**Supplemental Table S4** Attributions for the main vibrational bands identified in the bio- fingerprint region.

## Acknowledgments

We thank Dr. Paulo Cavalcanti Gomes Ferreira (*in memoriam*) for his scientific invaluable contribution to the plant cell cycle field and the discussions of the data in this manuscript.

## Author contributions

The project was designed by A.S.H. The experiments were carried out by P.M, J.F.R.P, C.N.M.C, L.D, L.P, A.F.F, H.F.B, VI, L.M.C, F.S.C and J.A.E. Measurements using the ATR-FTIR equipment were carried out by P.M, J.C.A, LT and CACJ; and measurements with the equipment Li-Cor 6400XT were made by P.M, W.d.P.B and E.C. The manuscript was written by P.M and J.F.R.P, with contributions from the other authors. Reviewing and editing was performed by A.S.H. A.S.H. is the principal investigator. All authors have read and agreed to the published version of the manuscript.

