## Supplemental Figures and Tables for "A novel *Arabidopsis thaliana* protein, ABAP1 INTERACTING PROTEIN 10, mediates crosstalk between the cell cycle and primary metabolism"

The following Supporting Information is available for this article:

**Supplemental Figure S1**. Protein interaction screening by yeast two-hybrid to identify potential AIP10 binding partners. AIP10 protein interaction with ABAP1, ARIA, KIN11 and pre-replication complex components ORC1-6, CDT1 and CDC6 was evaluated. The empty vectors GAL4 AD and GAL4 BD were used as negative controls, while interaction of AIP10 with ABAP1 was used as positive control. Growth of yeast strain on selective media SD-Leucine/- Tryptophan confirmed transformation with both BD/AD constructs. Growth on stringent selective media SD-Leucine/-Tryptophan/-Histidine/-Adenine confirmed strong protein interactions between AIP10 and ABAP1, and indicated that this interaction was mediated by the Armadillo domain of ABAP1. A11 (SnRK1 catalytic subunitKIN11); ARIA (ARM REPEAT PROTEIN INTERACTING WITH ABF2); ORC (ORIGIN RECOGNITION COMPLEX); ARM (ARMADILLO repeat domain of ABAP1); BTB (BTB/POZ domain of ABAP1).

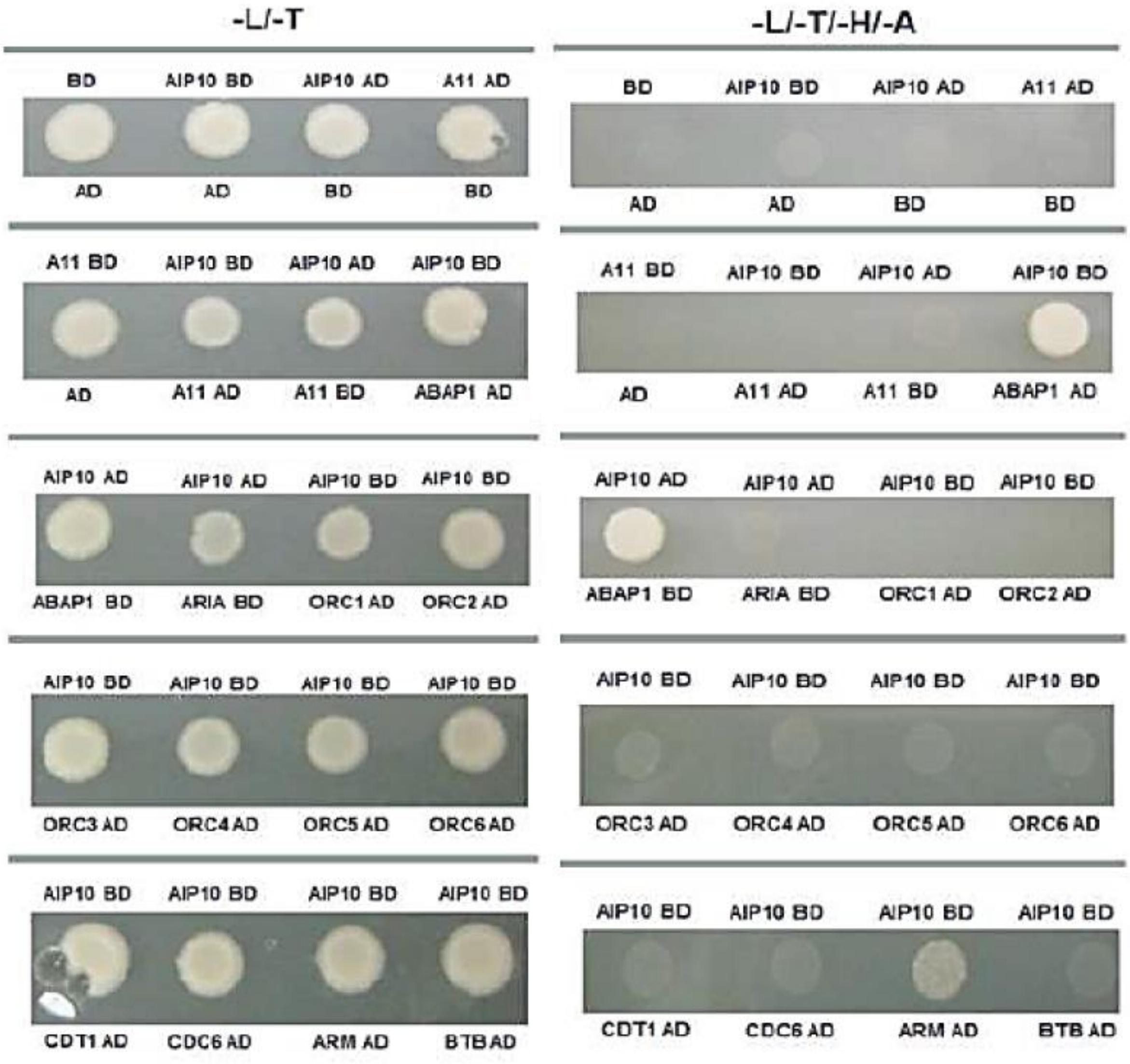

**Supplemental Figure S2**. Expression pattern of *AIP10* isoforms in different *A. thaliana* tissues. Characterization of the expression pattern of *AIP10.1*, *AIP10.2*, *AIP10.3* and *AIP10.4* isoforms in 11DAG roots grown *in vitro*; and young 11DAG seedlings, mature 32DAG rosettes, floral bud, open flowers and siliques of Col-0 plants grown directly *in vivo*, under a 16h/8h light photoperiod. The mRNA levels for each isoform were normalized by the mRNA levels of the constitutive genes *UBI14* and *GAPDH*. Bars indicate mean ± standard deviation. The primers were designed in regions that vary between the isoforms and are individually represented in the diagram in Figure S5.

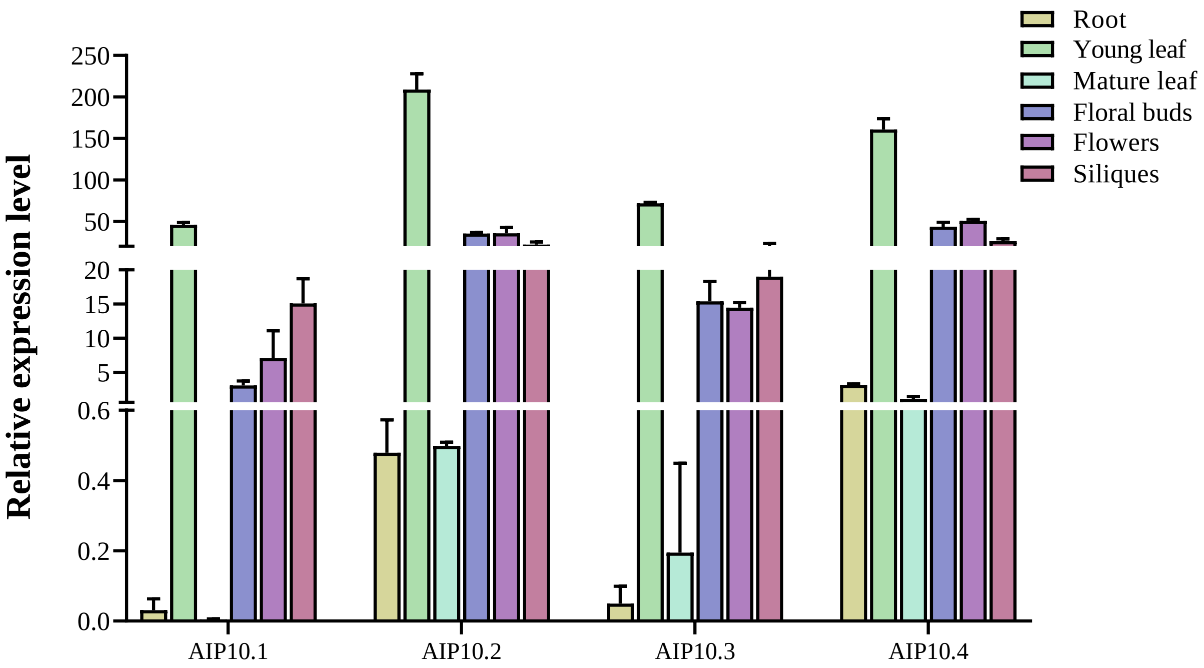

**Supplemental Figure S3.** Phylogenetic analysis of AIP10 and its homologs. **A)** Phylogenetic tree was generated in the IQTREE software using the Maximum Likelihood method using the JTT + Invar (I) + Gamma (G) model with 4 categories subjected to the Bootstrap phylogeny test (1000 repetitions). **B)** Amino acid alignment performed using the Jalview software using the CLUSTAL W method. The image comprises the AIP10 C-terminal region, conserved in all species and which contains the conserved “Coiled coil” domain, a nuclear localization signal (NLS) and a PPS. The protein sequences used in this tree were obtained from the Phytozome, Plaza and Sol Genomics databases.

**A**

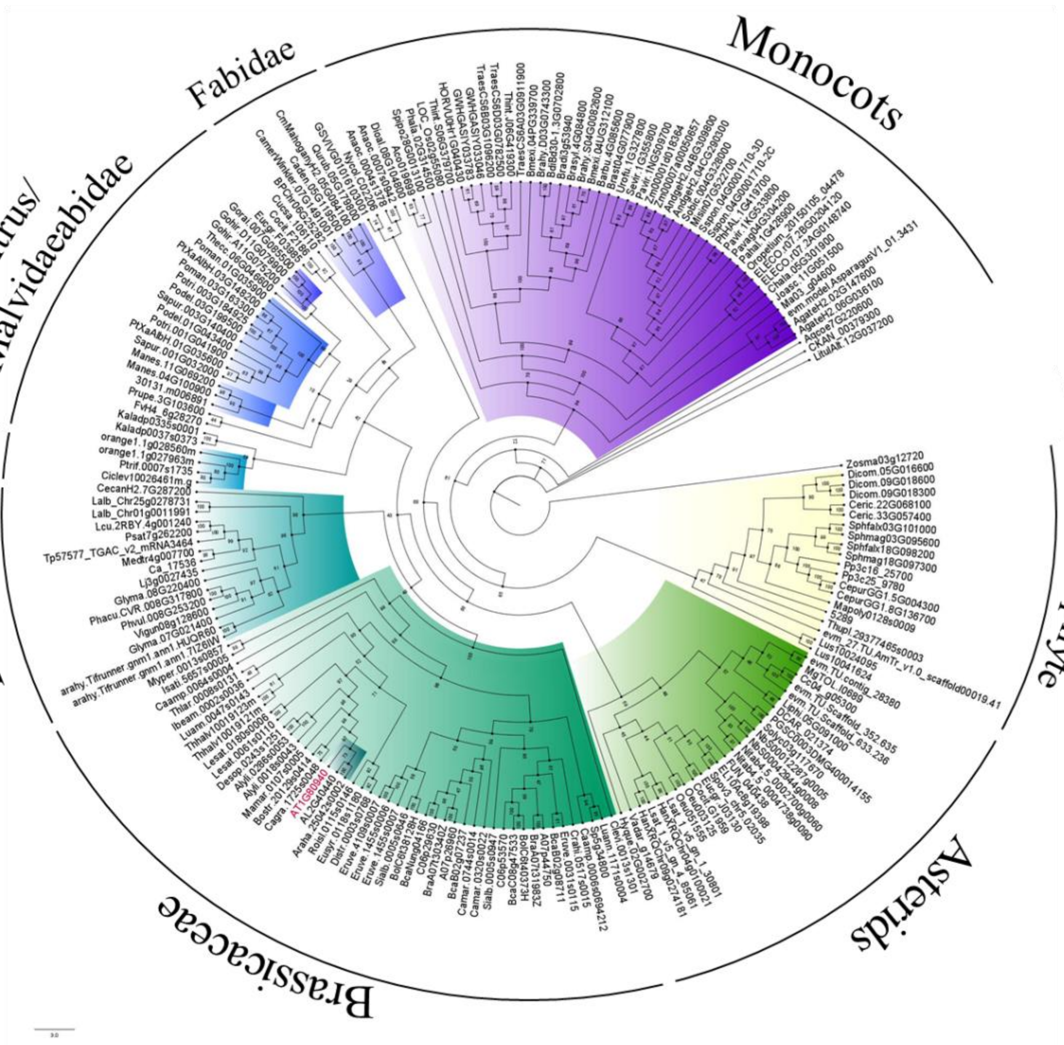

B

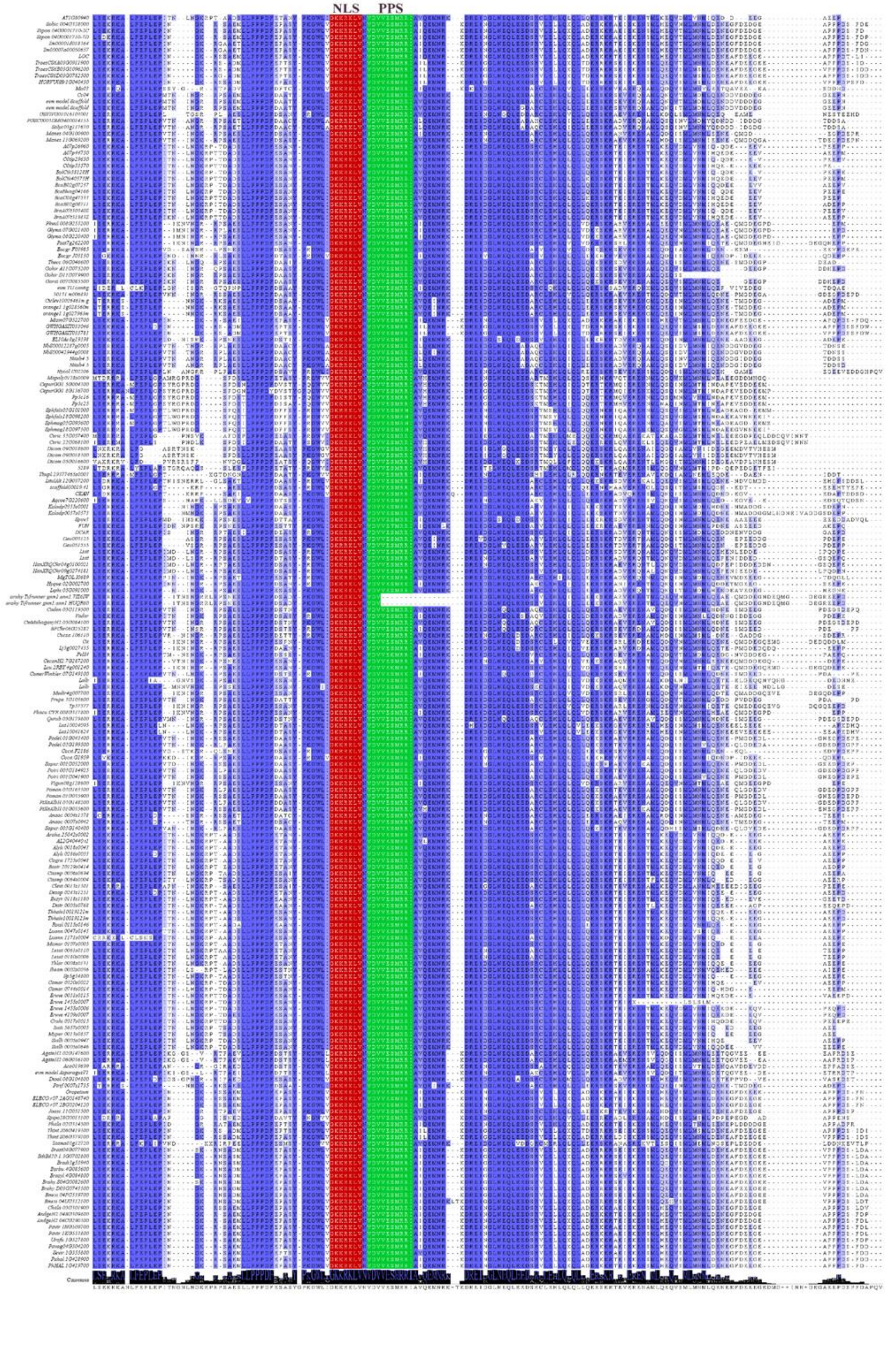

**Supplemental Figure S4.** Phylogenetic analysis of ABAP1 and its homologs. The phylogenetic tree was generated in the IQTREE software using the Maximum Likelihood method using the JTT + Invar (I) + Gamma (G) model with 4 categories subjected to the Bootstrap phylogeny test (1000 repetitions). The protein sequences used were obtained from the Phytozome, Plaza and Sol Genomics databases.

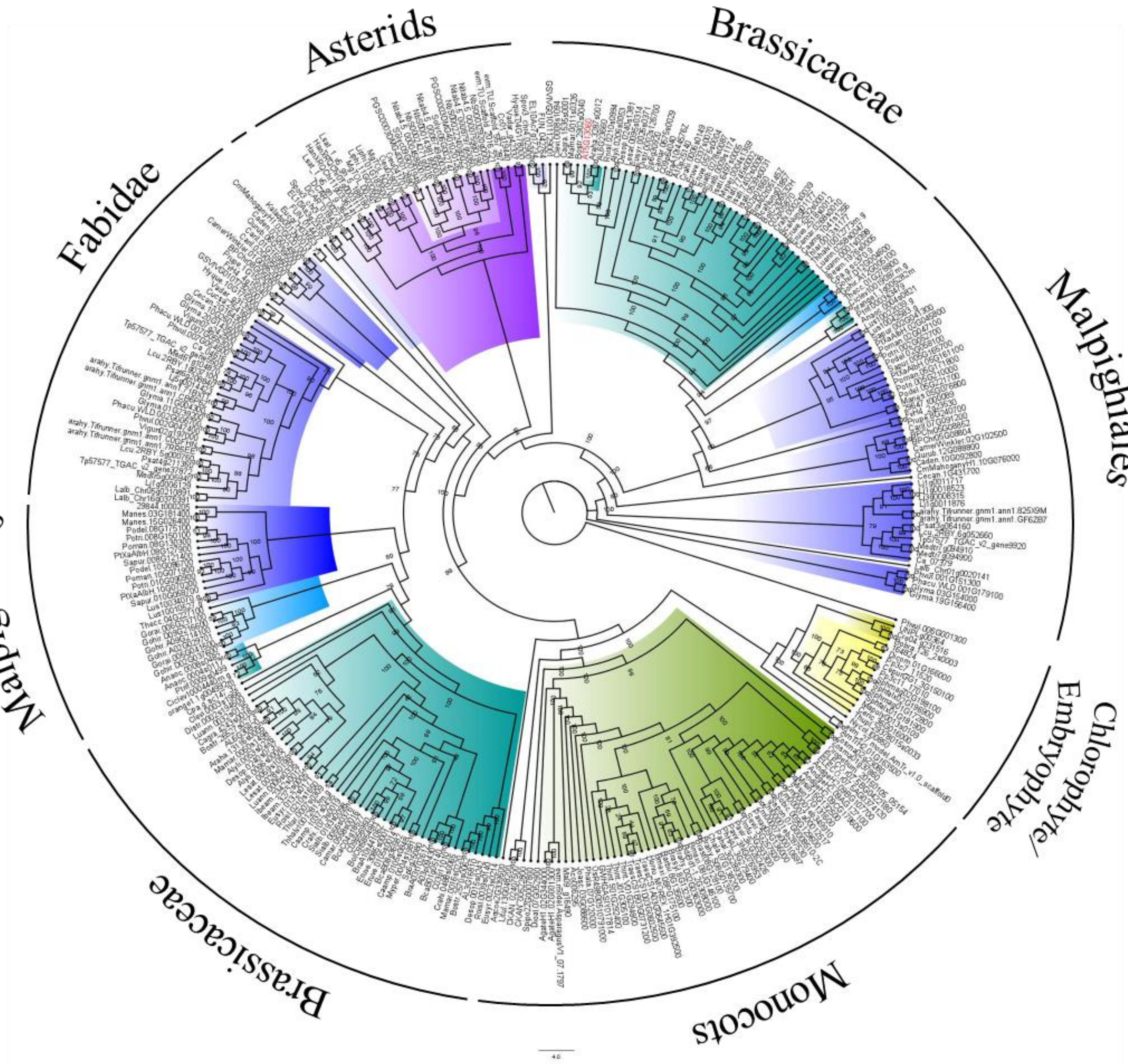

**Supplemental Figure S5.** Molecular characterization of *AIP10* mutant lines. **A)** Map of *AIP10* alternatively spliced isoforms showing the T-DNA insertion site in Salk mutants. Exons are represented by yellow rectangles and gray ones correspond to the 5' and 3' UTR regions. The solid gray line represents the intronic regions and the dotted black line represents the estimated position of the T-DNA in each alternative splicing isoforms of *AIP10*. Specific primers were used for each isoform to evaluate relative expression, as indicated in Figure S2**.** Brown arrows represent primers for isoform AIP10.1, green arrows for AIP10.2, blue arrows for AIP10.3, and pink arrows for AIP10.4. **B)** *AIP10* expression in *aip10-1* (Salk_022332) and *aip10-2* (Salk_094618) seedlings relative to Col-0 was analyzed by qRT-PCR. The positions of the primers are indicated as qRT.Fw and qRT.Rev. Expression levels were normalized by the mRNA levels of the *UBI14* and *GAPDH* genes. Bars represent means ± SD (standard deviation) and * significantly different from Col-0 with p<0.05 with Student's test. (n=3 biological replicates).

**A**

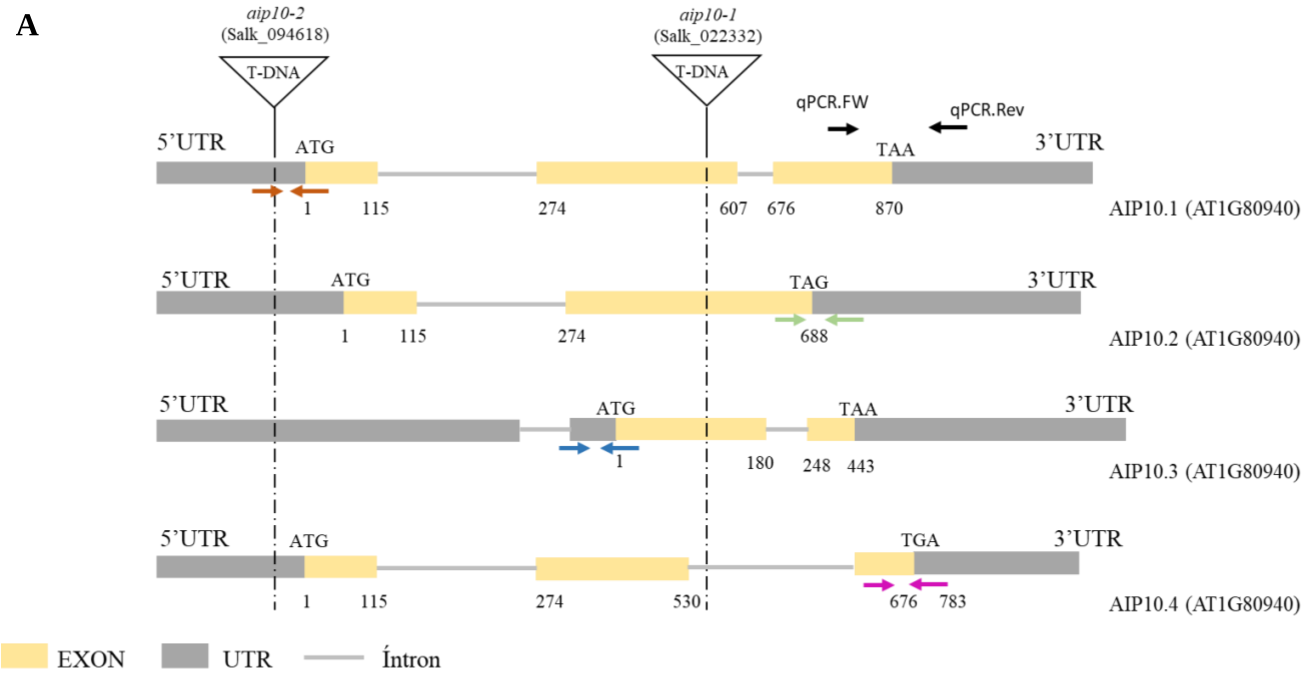


**B**

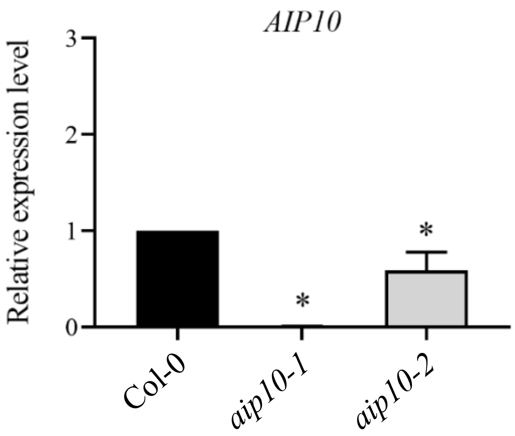

**Supplemental Figure S6.** Analyzes of root and leaf growth parameters in *aip10-1*, compared with wild-type Col-0. **A)** Expression analysis of *ABAP1*, *CyclinB1;1*, *CyclinB1;2* and the ABAP1 target genes, *CDT1a* and *CDT1b,* in Col-0 and *aip10-1*. Relative mRNA levels were evaluated in both genotypes in 11 DAG roots cultivated in vitro in MS medium with 1% sucrose in a 12h/12h photoperiod at 21ºC. **B)** Primary root length (n=10). **C)** Number of lateral roots (n=10). **D)** Leaf area and **E)** Cell area evaluated by kinematic growth analysis of the first leaf pair, from 6 to 18 DAG. Bars represent means ± SD (standard deviation) of the 5 plants.

**A**

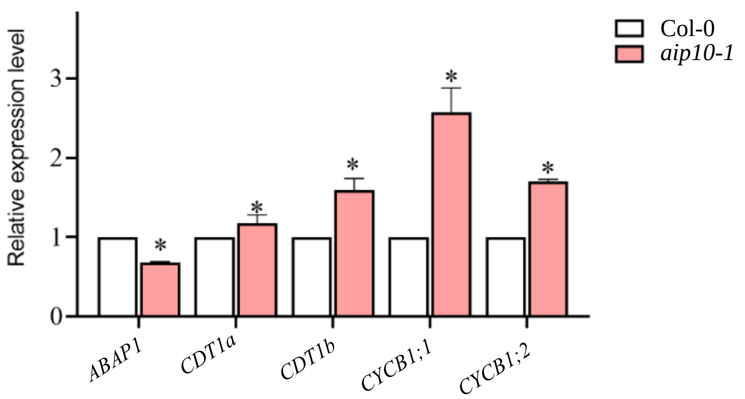


**B**

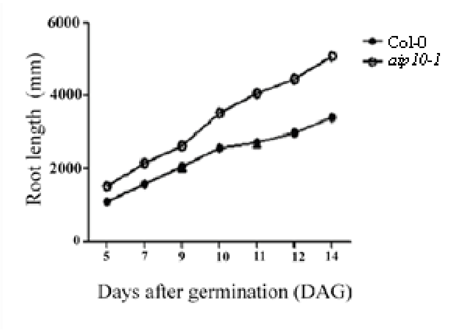


**C**

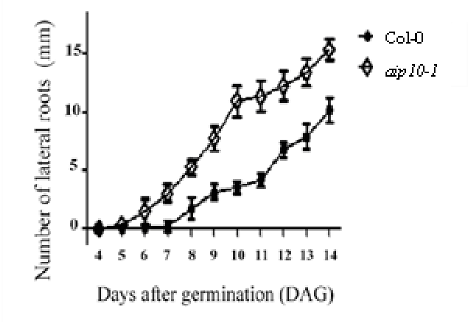


**D**

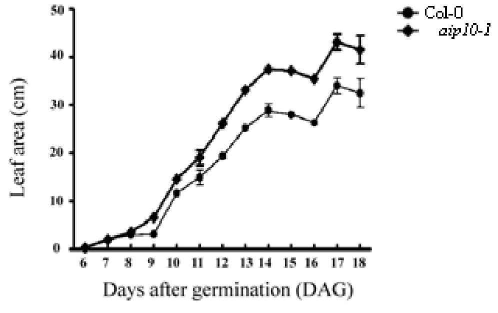


**E**

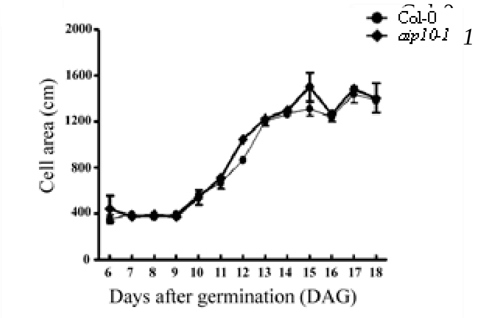

**Supplemental Figure S7.** Analyzes of *CDT1a* expression in meristems of *aip10-1* plants compared to wild-type Col-0. **A)** GUS staining of SAM (left panels) and RAM (right panels) from (A1,A2) Col-0 x *pCDT1a::GUS* and (A3,A4) *aip10-1* x *pCDT1a::GUS* seedlings at 5 DAG; black bars: 0.1mm. **B)** Graph with quantification of expression of *pCDT1a::GUS* in shoots of Col-0 and *aip10-1* seedlings at 5 DAG. The parameter for quantification of GUS expression is represented here as the measurement of GUS staining area of RAM and SAM with the ImageJ software with values given in arbitrary units.

**A**

Col-0 x p*CDT1a::GUS*

*aip10-1* x p*CDT1a::GUS*

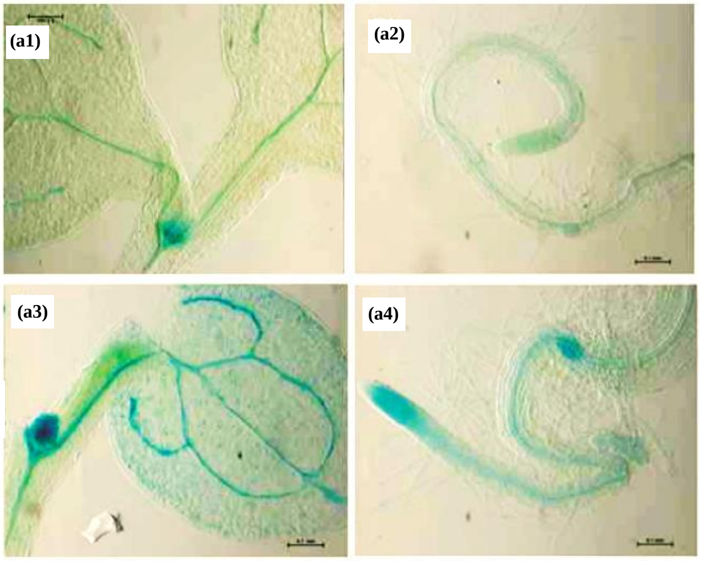


**B**

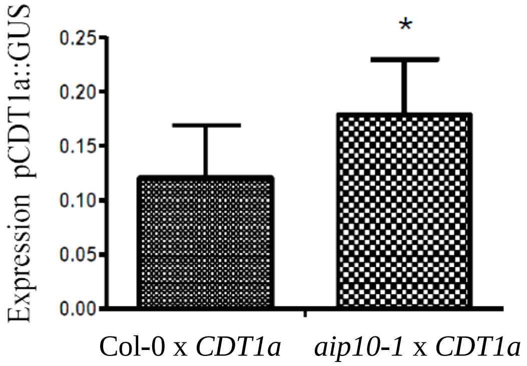

**Supplemental Figure S8.** Average nuclear DNA content analyzed by flow cytometry. The graph shows the ploidy level measured by flow cytometry of the first pair of leaves and roots of Arabidopsis Col-0 and *aip10-1* plants at **A)** 14 DAG and **B)** 20 DAG. **C)** Bar of endoreduplication Index (EI). Bars represent means ± SD (standard deviation) and * significantly different from Col-0 with $p<0.05$ with Student's test. Data were obtained from three biological replicates.

A

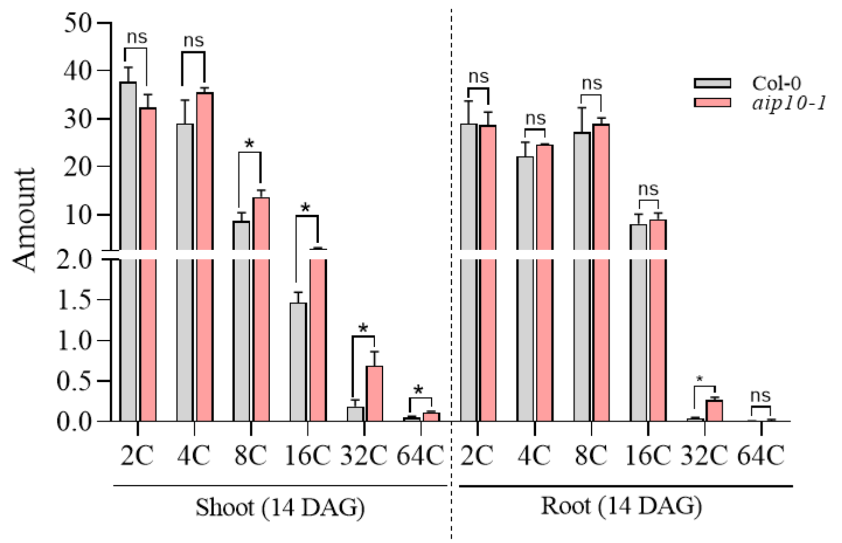


B

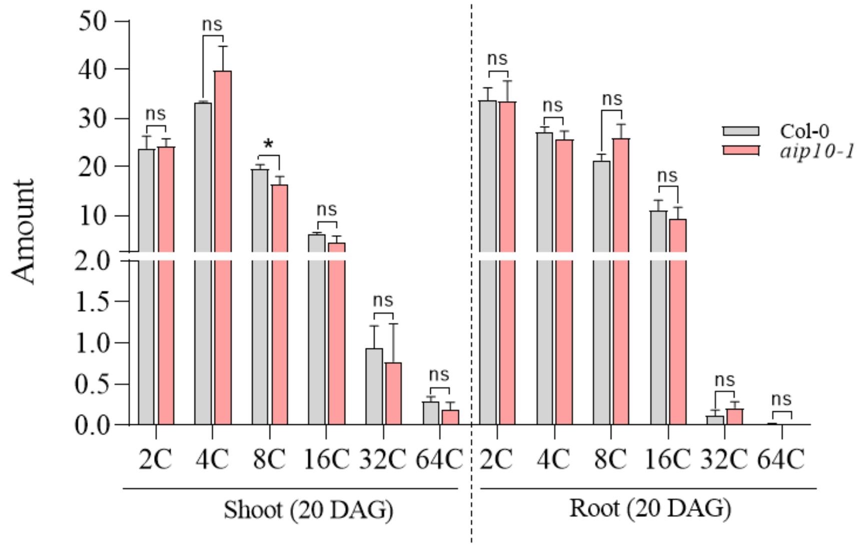

**C**

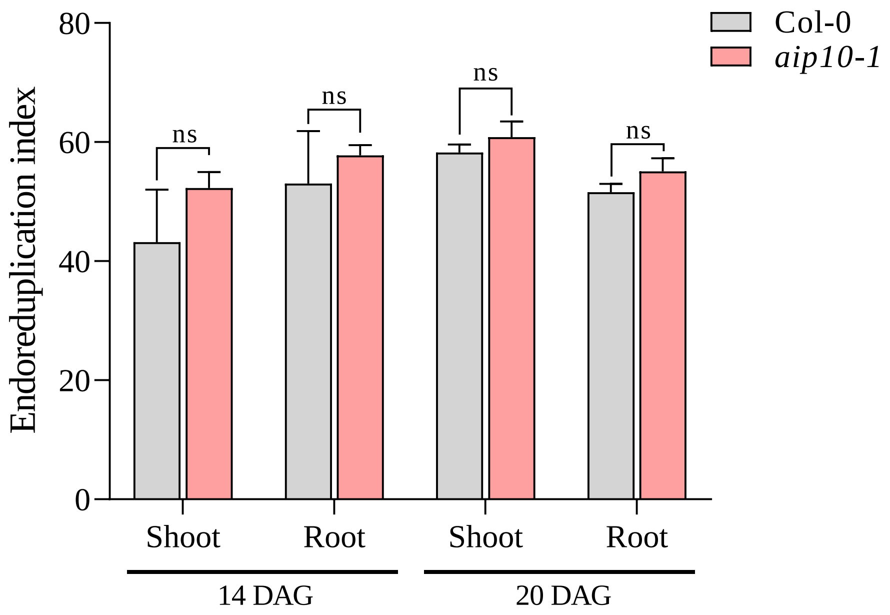
Endoreduplication index
80
60
40
20
0
ns
ns
ns
ns
Col-0
aip10-1
Shoot
Root
Shoot
Root
14 DAG
20 DAG

**Supplemental Figure S9.** Comparative analysis of the expression profile of genes differentially expressed in both KIN10oe and *aip10-1*. **A)** Heatmap of the expression profile of genes regulated in KIN10oe/sugar starvation (lane 1; Baena-González et al., 2007), which were also DEGs in at least one of the *aip10-1* transcriptomes (shoot/root at 11/35 DAG), when compared to Col-0 under normal growth conditions through RNAseq (log2 FC ≥ 0.5 and ≤ - 0.5 with P value <0.05). Heatmaps show color-coded log2 FC values of up-regulated (red) or down-regulated (blue) genes. **B-E)** Functional categorization of DEGs induced or repressed in *aip10-1* transcriptomes of **B)** 11 DAG shoot, **C)** 11 DAG roots, **D)** 35 DAG shoot and **E)** 35 DAG roots, which were common to transcriptomes of KIN10oe/sugar deprivation.

**A**

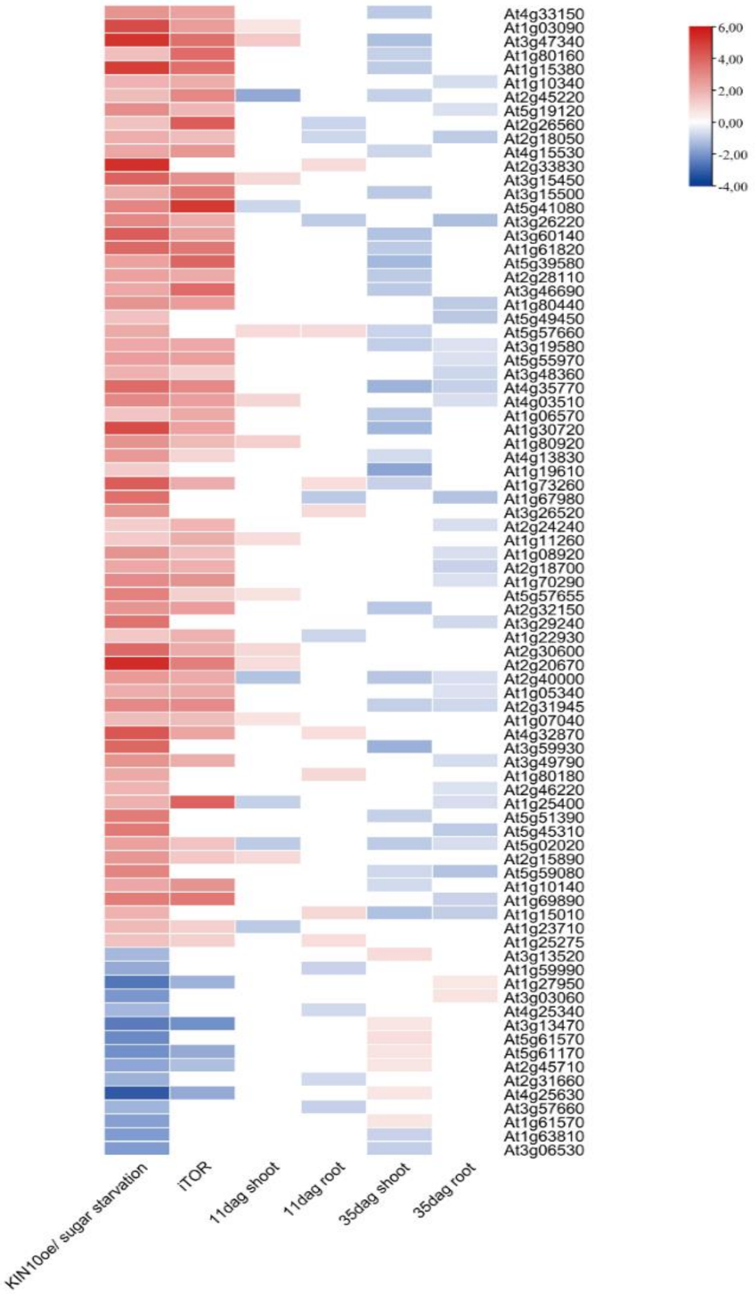

**B**

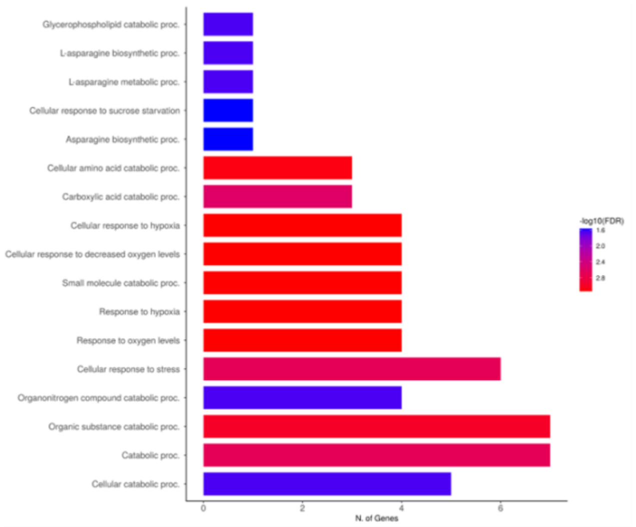
Glycerophospholipid catabolic proc.
L-asparagine biosynthetic proc.
L-asparagine metabolic proc.
Cellular response to sucrose starvation
Asparagine biosynthetic proc.
Cellular amino acid catabolic proc.
Carboxylic acid catabolic proc.
Cellular response to hypoxia
Cellular response to decreased oxygen levels
Small molecule catabolic proc.
Response to hypoxia
Response to oxygen levels
Cellular response to stress
Organonitrogen compound catabolic proc.
Organic substance catabolic proc.
Catabolic proc.
Cellular catabolic proc.
N. of Genes
-log10(FDR)

**C**

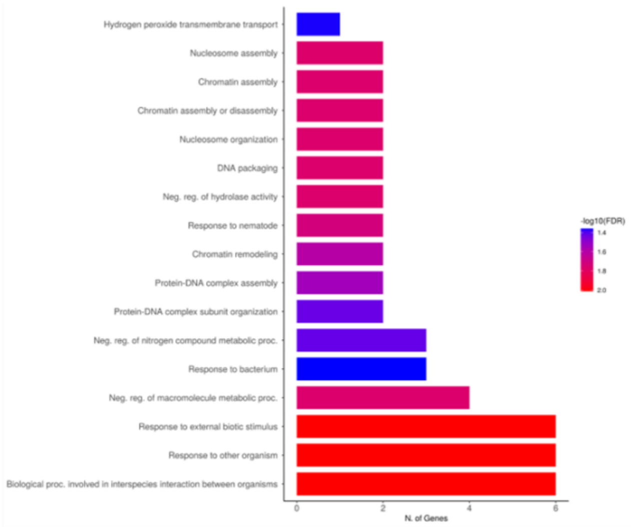
Hydrogen peroxide transmembrane transport
Nucleosome assembly
Chromatin assembly
Chromatin assembly or disassembly
Nucleosome organization
DNA packaging
Neg. reg. of hydrolase activity
Response to nematode
Chromatin remodeling
Protein-DNA complex assembly
Protein-DNA complex subunit organization
Neg. reg. of nitrogen compound metabolic proc.
Response to bacterium
Neg. reg. of macromolecule metabolic proc.
Response to external biotic stimulus
Response to other organism
Biological proc. involved in interspecies interaction between organisms
N. of Genes
-log10(FDR)

**D**

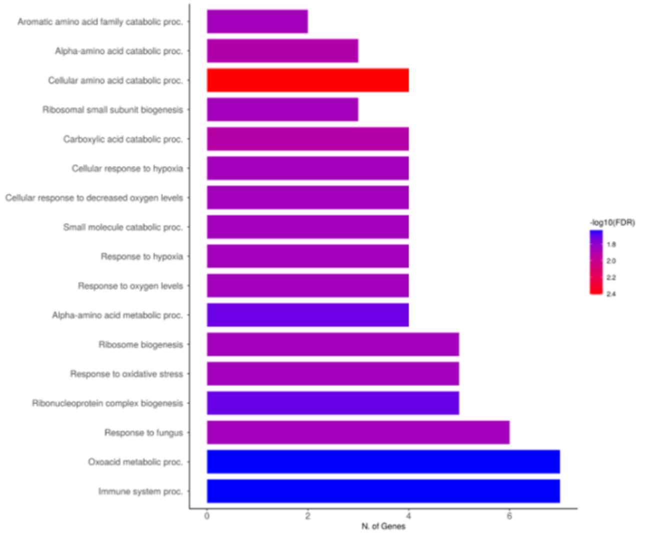
Aromatic amino acid family catabolic proc.
Alpha-amino acid catabolic proc.
Cellular amino acid catabolic proc.
Ribosomal small subunit biogenesis
Carboxylic acid catabolic proc.
Cellular response to hypoxia
Cellular response to decreased oxygen levels
Small molecule catabolic proc.
Response to hypoxia
Response to oxygen levels
Alpha-amino acid metabolic proc.
Ribosome biogenesis
Response to oxidative stress
Ribonucleoprotein complex biogenesis
Response to fungus
Oxoacid metabolic proc.
Immune system proc.
N. of Genes
-log10(FDR)

**E**

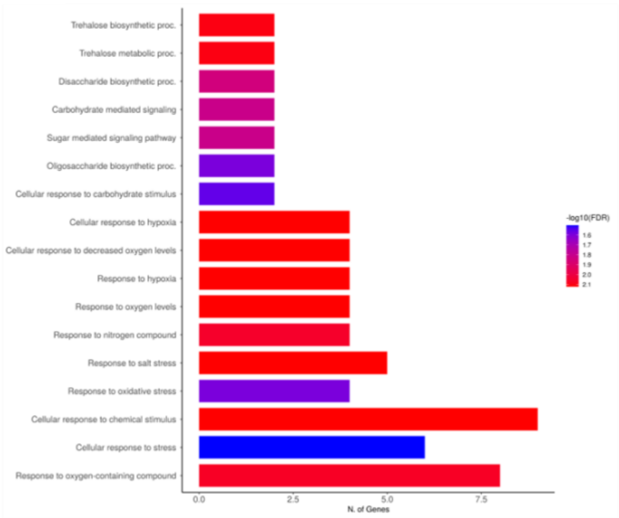
Trehalose biosynthetic proc.
Trehalose metabolic proc.
Disaccharide biosynthetic proc.
Carbohydrate mediated signaling
Sugar mediated signaling pathway
Oligosaccharide biosynthetic proc.
Cellular response to carbohydrate stimulus
Cellular response to hypoxia
Cellular response to decreased oxygen levels
Response to hypoxia
Response to oxygen levels
Response to nitrogen compound
Response to salt stress
Response to oxidative stress
Cellular response to chemical stimulus
Cellular response to stress
Response to oxygen-containing compound
N. of Genes
-log10(FDR)

**Supplemental Figure S10**. Comparative analysis of the expression profile of genes regulated by TOR inhibition differentially expressed in *aip10-1*. Heatmap of the expression profile of genes regulated by TOR inhibition (iTOR) which were also DEGs in the *aip10-1* transcriptomes, when compared to wild-type Col-0 under normal growth conditions, as determined by RNAseq (log2 FC ≥ 0.5 and ≤ - 0.5 with P value <0.05). Heatmaps show color-coded log2 FC values of up-regulated (red) or down-regulated (blue) genes.

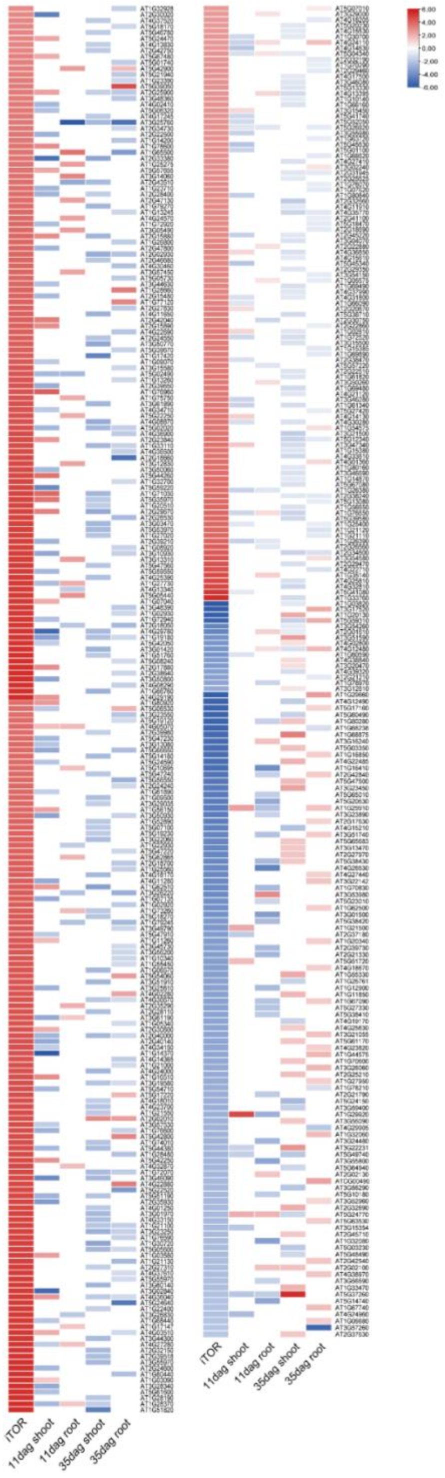

**Supplemental Figure S11.** Functional categorization of genes regulated in both iTOR and *aip10-1*. Analysis of Gene Ontology (GO) categories of differentially expressed genes induced or repressed in the *aip10-1* transcriptomes of **A)** shoot at 11 DAG, **B)** roots at 11 DAG, **C)** shoot at 35 DAG and **D)** roots at 35 DAG, which were common to the iTOR transcriptome as shown in heatmaps of supplementary figure 11.

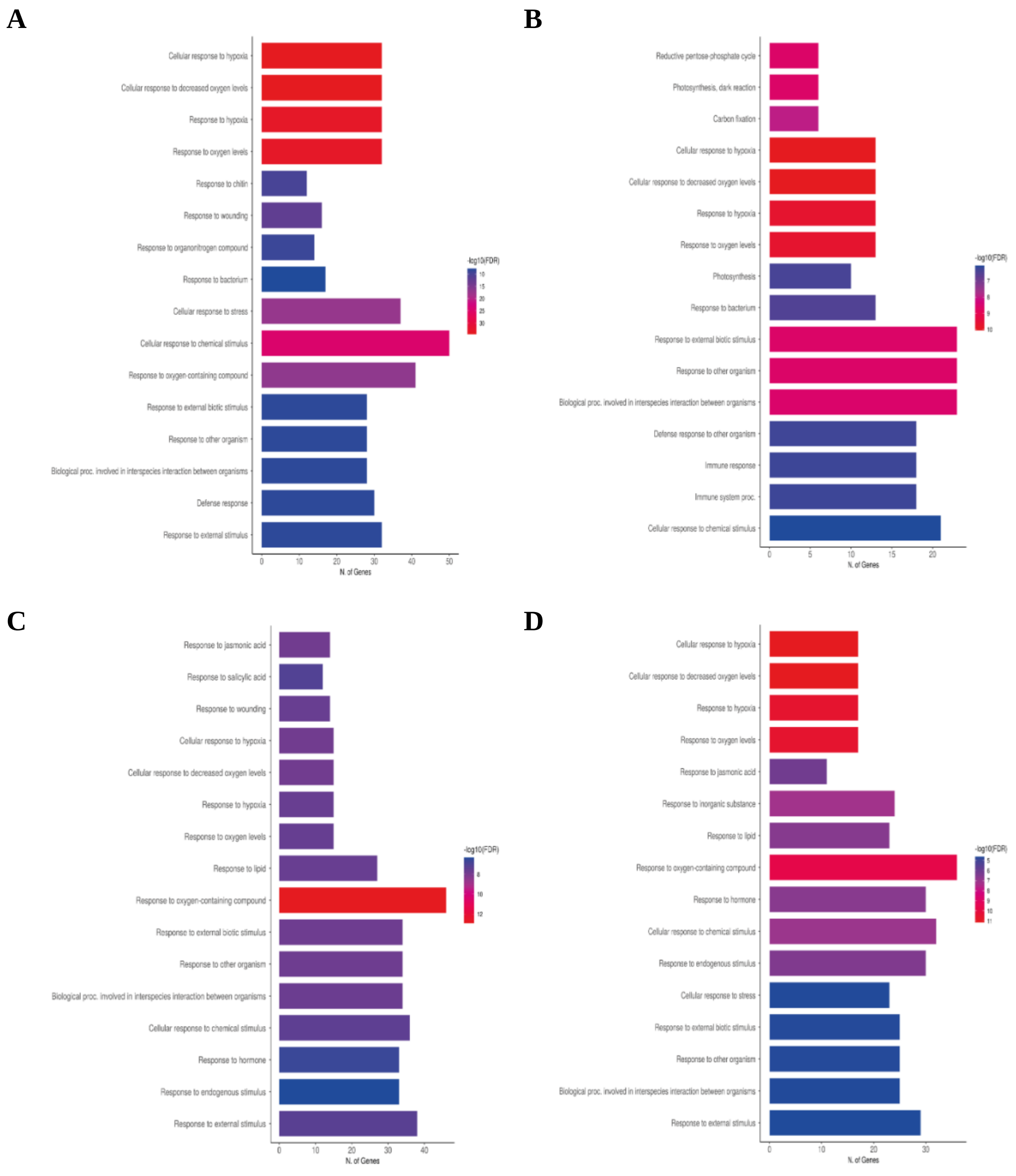

**Supplemental Figure S12.** Loading chart for the first two main components applied to the ATR-FTIR dataset. **A)** Col-0, *aip10-1* and *aip10-2* sheets from 35DAG plants. **B)** Col-0, *aip10-1* and *aip10-2* sheets from seeds.

A

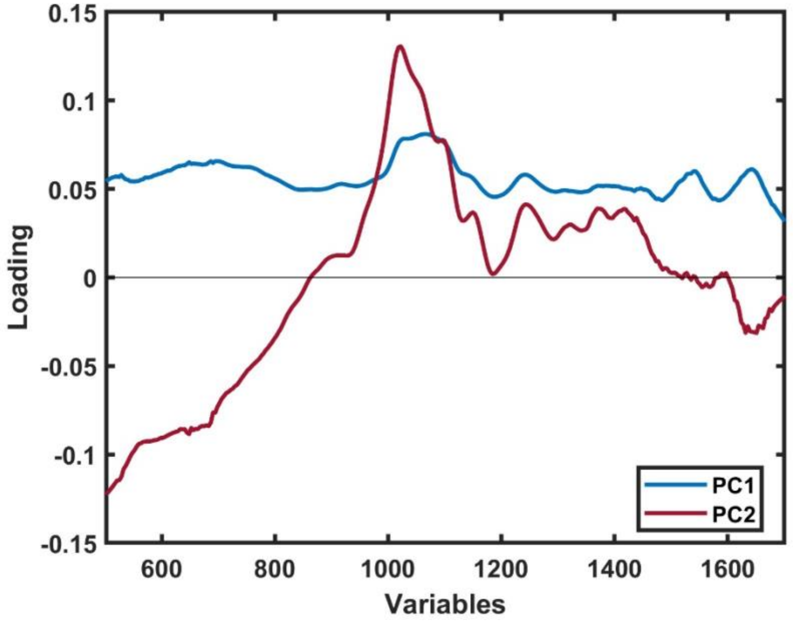


B

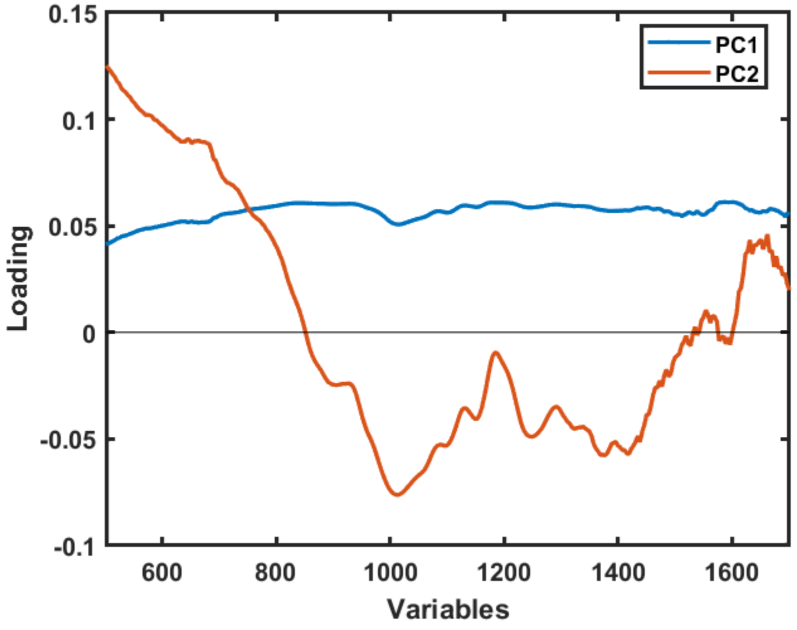

**Supplemental Figure S13.** Metabolic analysis by ATR-FTIR spectrum of *aip10-1* and *aip10-2* plants at 20 DAG, compared with wild-type Col-0. **A)** Average spectra obtained by the FTIR technique in the region of 900–1700 $cm^{-1}$ of Col-0 and *aip10-1* plants at 20 DAG. **B)** Average spectra obtained by the FTIR technique in the region of 900–1700 $cm^{-1}$ of Col-0 and *aip10-2* plants at 20 DAG. **C)** Score plot for three principal components applied to the 20DAG FTIR dataset. **D)** Loading Chart for the first two main components applied to the ATR-FTIR dataset. The experiment was carried out on 2 distinct leaves of 4 individuals of each genotype.

**A**

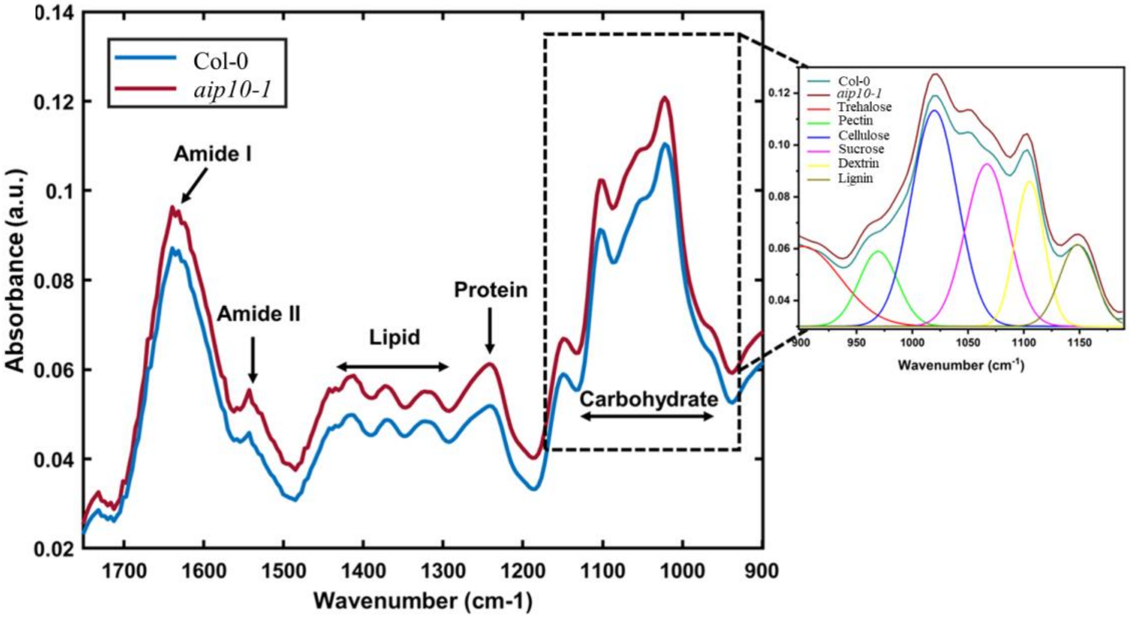


**B**

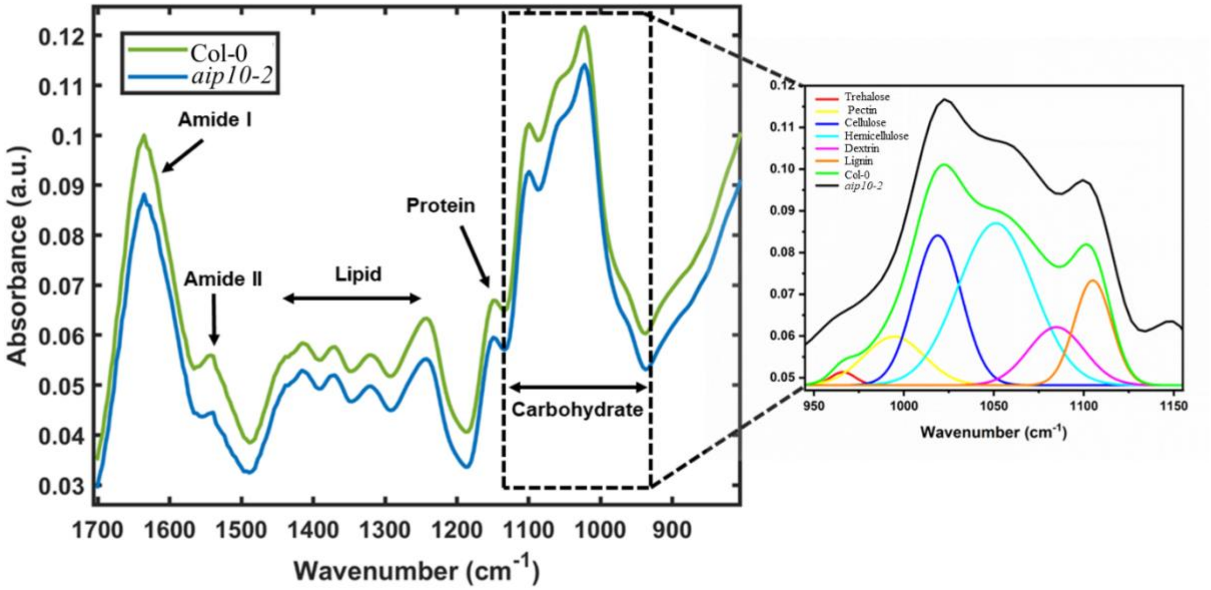

C

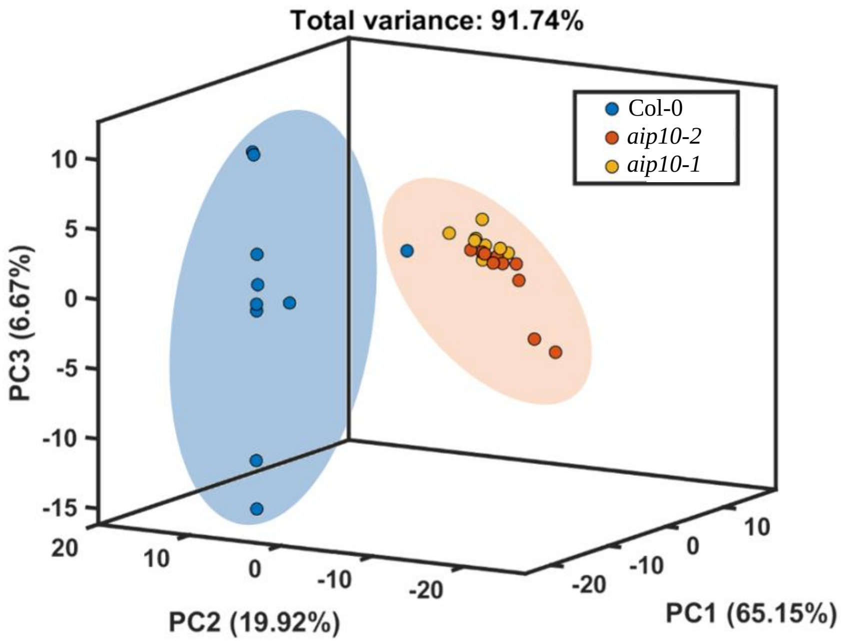
Total variance: 91.74%
Col-0
aip10-2
aip10-1
10
5
0
-5
-10
-15
PC3 (6.67%)
20
10
0
-10
-20
PC2 (19.92%)
-20
-10
0
10
PC1 (65.15%)

D

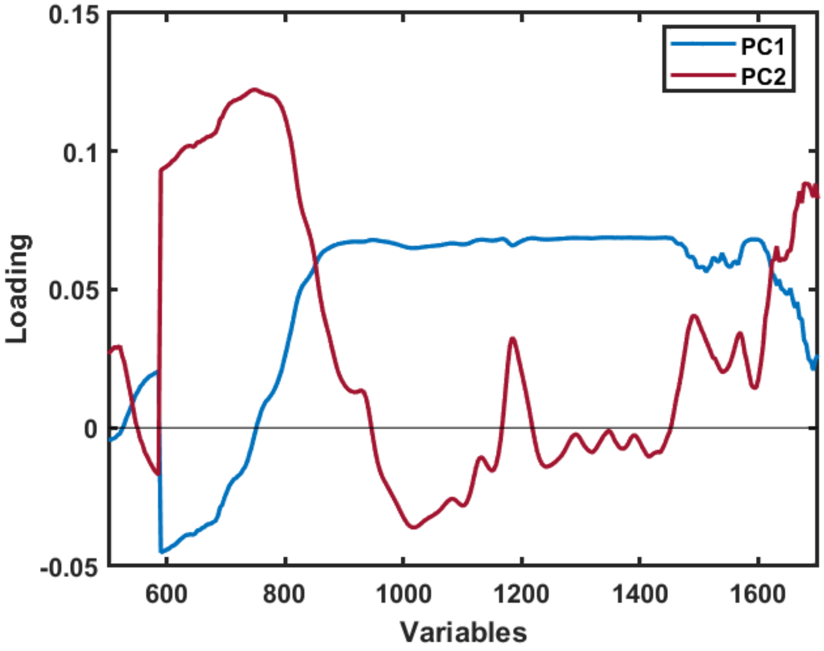
PC1
PC2
0.15
0.1
0.05
0
-0.05
Loading
600
800
1000
1200
1400
1600
Variables

**Supplemental Table S1**. Primers used in this study

| Primer name | Primer Forward Sequence | Primer Reverse Sequence |
|---|---|---|
| **AIP10 Salk Lines:** | | |
| T-DNA border primer (LB1.3) | ATTTTGCCGATTTCGGAAC | |
| SALK_022332 | ACACTCGAAATTCGTGGTCTG | CTTTCGTTTTGATCTGATCCG |
| SALK_094618 | AGCTTTCCTTTTCTCGGACAG | ATATTTTTCTTGGCTGCATGG |
| **Cloning:** | | |
| *KIN11* | GGGGACAAGTTTGTACAAAAAAGCAGGCTTCACAA TGGATCATTCATCAAATAGATTTGGCA | GGGGACCACTTTGTACAAGAAAGCTGGGTCTAAAA CATTCAGATCACACGAAGC |
| **qRT-PCR:** | | |
| *AIP10* | GGAGCTGAAGAACCCTAAGCTAGTT | GGACGAGACAAATCTACAAAAGAATAAA |
| *AIP10.1* | AAGAAGGTTAAAGGTTGCATG | TTTGGCGTTTTGGTTACGACTA |
| *AIP10.2* | GCTGGAGTTTGATTTACGGATTG | TCTGTTCTCTTGCTTCTCTCTTG |
| *AIP10.3* | GCTACCCTTTCTATTTCTGGTCT | GGTCACACCAGAGAAGAACGA |
| *AIP10.4* | CTGGTTGATCGGATCGAGAGATAGA | TCTGTTCTCTTGCTTCTCTCTTG |
| *ABAP1* | TCAGCCTTAAGAAGAGCTTGCA | ACCATAATTGAGAGCTGAGCTTAGTG |
| *Cdt1a* | AATCCGATCACGTCTTGAAGAAG | GAACCACGATCTCAAGAAAGCA |
| *Cdt1b* | AAATGTCGACTGCGAAACAG | AAGTGAAATGTCATGTGAAGTTGCTT |
| *Cyclin B1;1* | CCTCCATTCACTCTCAACAG | CCTGGCAGCTGTGGAATATG |
| *Cyclin B1;2* | TCAGTGCCTTGCTTATTGCTTCC | GTCGGGACTGTCAAATACCATTCG |
| *QQS* | GGTTCATTTTGCCTCACACTTCT | CCCATGATATGACCCTCATTTTG |
| *LHCB1* | TCAAGCCATCGTCACTGGTAAG | ACTGGATCGGCCAAATGGT |
| *LHCB6* | TTGCCATGTTGATCTTTTACTTTGA | CATCATGCCGTTTTCTCACAA |
| *LZF1* | CGAATCAAGGGTTGGGAAA | AGCGGCTAAAAGATAATTACAAATGC |
| *PSAD2* | GGTAAAAATGTTAGTCCCATTGAGGTT | CATCCGCAAGATTTTACATGTAGTG |
| *DJC22* | GGAGCAGGAACCCGTGATTA | ACACATTTGGATGATTTCTTTCGA |
| *DJC23* | AACCGATCAGTGTGGGTAGTGAGTT | GGATTTTATGGCTTCGTATTTACACTAA |
| *DJC24* | GCAGTTACGGTGGACGGAAT | GGACCAACTCGACACATCGA |
| *NAD1* | GACATTAGCCGAATCTGATCCAT | CAAGATAGAGACATAGAGAAGAAGAAGCA |
| *GBSS1* | GCGACGCCGTGAGAATATG | CTTCCCAACGCTTTCTTCAAA |
| *STP1* | GCAGCTCCAAAAAAGCAAGAA | TGTTGCCTTACGGGTTTAATCC |
| *ASN1/DIN6* | AACTTGTCGCCAGATCAAGG | GGAACACGTGCCTCTAGTCC |
| *SEN1* | CAGAGTCGGATCAGGAATGG | ATTTGACCGCTCTCACAACC |
| *TPS11* | TTCAACAGCCTATGGGACAATG | AAAATCTAACGCAAACCCTTTCAA |
| **Housekeeping genes:** | | |
| *GAPDH* | TTGGTGACAACAGGTCAAGCA | AAACTTGTCGCTCAATGCAATC |
| *UBI14* | TCACTGGAAAGACCATTACTCTTGAA | AGCTGTTTTCCAGCGAAGATG |

**Supplemental Table S2.** AIP10 orthologs in other plant species

| Gene ID | Isoforms | Scientific Name | Query Cover | Evalue | Per. Ident | Acc. Len |
|---|---|---|---|---|---|---|
| AT1G80940 | AT1G80940.1 | *Arabidopsis thaliana* | 100% | 9E-159 | 100.00% | 213 |
| | AT1G80940.2 | | 71% | 8E-109 | 98.03% | 175 |
| | AT1G80940.3 | | 70% | 7E-110 | 100.00% | 151 |
| | AT1G80940.4 | | 61% | 2E-91 | 96.15% | 158 |
| Araha.25042s0002 | Araha.25042s0002.1 | *Arabidopsis helleri* | 100% | 1E-139 | 92.52% | 214 |
| AL2G40440 | AL2G40440.t1 | *Arabidopsis lyrata* | 95% | 1E-131 | 93.14% | 214 |
| Aqcoe7G220600 | Aqcoe7G220600.1 | *A. coerulea* | 91% | 8E-69 | 59.61% | 231 |
| | Aqcoe7G220600.2 | | 84% | 8E-69 | 60.10% | 231 |
| | Aqcoe7G220600.3 | | 89% | 1E-64 | 72.14% | 166 |
| | Aqcoe7G220600.4 | | 65% | 1E-64 | 72.14% | 166 |
| AgateH2.02G147600 | AgateH2.02G147600.1 | *Agave tequilana var. Weber's Blue* | 84% | 3E-67 | 60.10% | 209 |
| | AgateH2.02G147600.2 | | 60% | 2E-38 | 52.86% | 158 |
| | AgateH2.02G147600.3 | | 58% | 5E-63 | 77.60% | 172 |
| AgateH2.06G036100 | AgateH2.06G036100.1 | | 89% | 4E-67 | 56.48% | 220 |
| | AgateH2.06G036100.2 | | 60% | 4E-38 | 49.67% | 158 |
| | AgateH2.06G036100.3 | | 89% | 4E-67 | 56.48% | 220 |
| Alyli.0286s0053 | Alyli.0286s0053.1 | *Alyssum linifolium* | 100% | 7E-133 | 89.86% | 216 |
| Alyli.0018s0043 | Alyli.0018s0043.1 | | 95% | 7E-130 | 90.87% | 217 |
| | Alyli.0018s0043.2 | | 95% | 7E-130 | 90.87% | 217 |
| evm_27.TU.AmTr_v1.0_scaffold00019.41 | evm_27.model.AmTr_v1.0_scaffold00019.41 | *Amborella trichopoda* | 86% | 7E-70 | 59.89% | 199 |
| Anaoc.0007s0942 | Anaoc.0007s0942.1 | *Anacardium occidentale* | 92% | 1E-74 | 64.85% | 207 |
| | Anaoc.0007s0942.2 | | 62% | 2E-46 | 61.48% | 171 |
| | Anaoc.0007s0942.3 | | 67% | 2E-64 | 72.11% | 184 |
| Anaoc.0004s1378 | Anaoc.0004s1378.1 | | 64% | 5E-63 | 74.47% | 178 |
| Aco019699 | Aco019699.1 | *Ananas comosus* | 90% | 5E-64 | 57.89% | 211 |
| AndgeH2.04BG309800 | AndgeH2.04BG309800.1 | *Andropogon Gerardi* | 89% | 1E-70 | 61.31% | 204 |
| AndgeH2.04CG290300 | AndgeH2.04CG290300.1 | | 89% | 1E-67 | 60.40% | 207 |
| arahy.Tifrunner.gnm1.ann1.HUQR60 | arahy.Tifrunner.gnm1.ann1.HUQR60.1 | *Arachis hypogaea* | 92% | 2E-71 | 56.65% | 203 |
| arahy.Tifrunner.gnm1.ann1.7IZ6IW | arahy.Tifrunner.gnm1.ann1.7IZ6IW.1 | | 92% | 2E-71 | 56.65% | 203 |
| evm.model.AsparagusV1_01.3431 | evm.model.AsparagusV1_01.3431 | *Asparagus officinalis* | 89% | 8E-69 | 53.21% | 224 |
| EL10Ac8g19398 | EL10Ac8g19398 | *Beta vulgaris* | 90% | 6E-80 | 63.13% | 209 |
| BPChr06G25282 | BPChr06G25282 | *Betula platyphylla* | 80% | 8E-72 | 65.70% | 219 |
| Bostr.20129s0414 | Bostr.20129s0414.1 | *Boechera stricta* | 100% | 9E-139 | 93.09% | 216 |
| | Bostr.20129s0414.2 | | 100% | 9E-139 | 93.09% | 216 |
| Barbu.4G085600 | Barbu.4G085600.1 | *Brachypodium arbuscula* | 89% | 9E-72 | 59.52% | 214 |
| BdiBd30-1.3G0702800 | BdiBd30-1.3G0702800.1 | *Brachypodium distachyon* | 89% | 3E-70 | 59.05% | 214 |
| Bradi3g53940 | Bradi3g53940.1 | | 89% | 4E-69 | 58.57% | 214 |
| Bmexi.04PG339700 | Bmexi.04PG339700.1 | *Brachypodium mexicanum* | 86% | 2E-68 | 57.48% | 214 |
| | Bmexi.04PG339700.2 | | 86% | 2E-68 | 57.48% | 214 |
| | Bmexi.04PG339700.3 | | 86% | 2E-68 | 57.48% | 214 |
| | Bmexi.04PG339700.4 | | 33% | 7E-38 | 84.72% | 128 |
| | Bmexi.04PG339700.5 | | 63% | 3E-68 | 76.98% | 196 |
| Bmexi.04UG312100 | Bmexi.04UG312100.1 | | 89% | 7E-68 | 58.10% | 217 |
| | Bmexi.04UG312100.2 | | 89% | 5E-70 | 58.94% | 214 |
| | Bmexi.04UG312100.3 | | 89% | 5E-70 | 58.94% | 214 |
| Brahy.D03G0743300 | Brahy.D03G0743300.1 | *Brachypodium hybridum* | 84% | 2E-67 | 59.69% | 214 |
| Brahy.S04G0082600 | Brahy.S04G0082600.1 | | 89% | 2E-70 | 59.42% | 214 |
| Brasyl.4G084800 | Brasyl.4G084800.1 | *Brachypodium sylvaticum* | 89% | 1E-71 | 59.05% | 214 |
| Brast04G077900 | Brast04G077900.1 | *Brachypodium stacei* | 89% | 2E-70 | 59.42% | 214 |
| BcaB02g07237 | BcaB02g07237 | *Brassica Carinata* | 100% | 6E-128 | 87.73% | 220 |
| BcaNung04166 | BcaNung04166 | | 100% | 7E-128 | 86.30% | 217 |
| BcaC08g47533 | BcaC08g47533 | | 95% | 1E-122 | 82.79% | 224 |
| BcaB02g08711 | BcaB02g08711 | | 100% | 4E-120 | 83.48% | 224 |
| A07p26960 | A07p26960 | *Brassica napus* | 100% | 1E-128 | 86.30% | 219 |
| C06p29630 | C06p29630 | | 100% | 2E-128 | 86.76% | 218 |
| A07p44750 | A07p44750 | | 95% | 3E-121 | 85.45% | 223 |
| C06p53570 | C06p53570 | | 97% | 2E-120 | 83.49% | 220 |
| BolC6t38128H | BolC6t38128H | *Brassica oleracea* | 100% | 1E-128 | 87.56% | 216 |
| BolC6t40373H | BolC6t40373H | | 95% | 1E-122 | 82.79% | 225 |
| BraA07t30340Z | BraA07t30340Z | *Brassica rapa* | 100% | 1E-128 | 86.30% | 219 |
| BraA07t31983Z | BraA07t31983Z | | 95% | 3E-120 | 84.98% | 223 |
| Camar.0320s0022 | Camar.0320s0022.1 | *Cakile maritima* | 100% | 2E-123 | 84.68% | 220 |
| Camar.0744s0014 | Camar.0744s0014.1 | | 97% | 3E-121 | 84.11% | 209 |
| Cagra.1725s0048 | Cagra.1725s0048.1 | *Capsella grandiflora* | 100% | 6E-139 | 93.09% | 216 |
| | Cagra.1725s0048.2 | | 100% | 6E-139 | 93.09% | 216 |
| evm.TU.contig_28380 | evm.TU.contig_28380.1 | *Carica papaya* | 57% | 3E-56 | 75.41% | 151 |
| Cocit.G1959 | Cocit.G1959.1 | *Carya illinoinensis* | 93% | 1E-71 | 61.19% | 208 |
| Cocit.F2186 | Cocit.F2186.1 | | 90% | 3E-69 | 60.91% | 216 |
| Caden.05G119500 | Caden.05G119500.1 | *Castanea dentata* | 100% | 3E-87 | 62.04% | 220 |
| | Caden.05G119500.2 | | 81% | 1E-75 | 66.47% | 177 |
| | Caden.05G119500.3 | | 81% | 1E-75 | 66.47% | 177 |

| | | | | | | |
|---|---|---|---|---|---|---|
| CmMahoganyH2.05G084100 | CmMahoganyH2.05G084100.1 | *Castanea mollissima Mahogany* | 100% | 3E-87 | 62.04% | 220 |
| | CmMahoganyH2.05G084100.2 | | 81% | 1E-75 | 66.47% | 177 |
| Caamp.0006s0694 | Caamp.0006s0694.1 | *Caulanthus amplexicaulis* | 100% | 7E-128 | 87.10% | 212 |
| Caamp.0064s0004 | Caamp.0064s0004.1 | | 99% | 8E-128 | 87.04% | 216 |
| CepurGG1.5G004300 | CepurGG1.5G004300 | *Ceratodon purpureus GG1* | 61% | 8E-59 | 66.91% | 338 |
| CepurGG1.8G136700 | CepurGG1.8G136700 | | 61% | 9E-56 | 65.22% | 216 |
| Ceric.33G057400 | Ceric.33G057400.1 | *Ceratopteris richardii* | 87% | 2E-60 | 53.23% | 235 |
| | Ceric.33G057400.2 | | 62% | 9E-61 | 67.67% | 165 |
| Ceric.22G068100 | Ceric.22G068100.1 | | 61% | 4E-58 | 68.18% | 233 |
| CecanH2.7G287200 | CecanH2.7G287200.1 | *Cercis canadenses* | 96% | 2E-89 | 63.81% | 212 |
| | CecanH2.7G287200.2 | | 66% | 2E-54 | 63.83% | 148 |
| Chala.05G301900 | Chala.05G301900.1 | *Chasmanthium laxum* | 89% | 5E-67 | 60.00% | 209 |
| Ca_17536 | Ca_17536 | *Cicer arietinum* | 92% | 6E-87 | 64.97% | 199 |
| CKAN_00379800 | CKAN_00379800 | *Cinnamomum kanehirae* | 89% | 2E-68 | 60.62% | 206 |
| Ciclev10026461m.g | Ciclev10026461m.g | *Citrus clementina* | 97% | 2E-75 | 61.61% | 216 |
| orange1.1g028560mg | orange1.1g028560mg | *Citrus sinensis* | 97% | 1E-73 | 60.48% | 207 |
| orange1.1g027963mg | orange1.1g027963m | | 97% | 2E-70 | 61.61% | 216 |
| Clevi.0013s1301 | Clevi.0013s1301.1 | *Cleome violacea* | 95% | 3E-103 | 78.33% | 213 |
| evm.TU.Scaffold_633.236 | evm.model.Scaffold_633.236 | *Coffea arabica* | 92% | 8E-74 | 62.63% | 208 |
| evm.TU.Scaffold_352.635 | evm.model.Scaffold_352.635 | | 92% | 5E-75 | 63.64% | 208 |
| Cc04_g05300 | Cc04_g05300 | *Coffea canephora* | 92% | 5E-75 | 63.64% | 208 |
| CamerWinkler.07G149100 | CamerWinkler.07G149100.1 | *Corylus americana* | 96% | 6E-88 | 66.67% | 213 |
| | CamerWinkler.07G149100.2 | | 75% | 2E-74 | 70.00% | 179 |
| Crahi.0517s0015 | Crahi.0517s0015.1 | *Crambe hispanica* | 95% | 7E-119 | 86.47% | 211 |
| Cucsa.106110 | Cucsa.106110.1 | *Cucumis sativus* | 64% | 3E-74 | 73.38% | 141 |
| DCAR_021374 | DCAR_021374 | *Daucus carota* | 91% | 6E-87 | 65.46% | 206 |
| Desop.0243s1251 | Desop.0243s1251.1 | *Descurainia sophioides* | 95% | 2E-128 | 90.34% | 217 |
| Dioal.08G104800 | Dioal.08G104800.1 | *Dioscorea alata* | 86% | 6E-77 | 63.27% | 210 |
| Dicom.09G018300 | Dicom.09G018300.1 | *Diphasiastrum complanatum* | 61% | 2E-55 | 66.92% | 209 |
| | Dicom.09G018300.2 | | 61% | 2E-55 | 66.92% | 209 |
| Dicom.09G018600 | Dicom.09G018600.1 | | 61% | 2E-55 | 66.92% | 209 |
| Dicom.05G016600 | Dicom.05G016600.1 | | 63% | 3E-49 | 60.74% | 211 |
| | Dicom.05G016600.2 | | 63% | 6E-50 | 60.74% | 149 |
| Distr.0003s0768 | Distr.0003s0768.1 | *Diptychocarpus strictus* | 94% | 8E-124 | 87.25% | 217 |
| ELECO.r07.2BG0204120 | ELECO.r07.2BG0204120.1 | *Eleusine coracana* | 89% | 2E-70 | 60.19% | 209 |
| ELECO.r07.2AG0148740 | ELECO.r07.2AG0148740.1 | | 89% | 2E-70 | 59.51% | 209 |
| Eruve.1455s0006 | Eruve.1455s0006.1 | *Eruca vesicaria* | 97% | 6E-127 | 87.04% | 222 |
| Eruve.4109s0007 | Eruve.4109s0007.1 | | 97% | 4E-126 | 86.57% | 222 |
| | Eruve.4109s0007.2 | | 75% | 4E-100 | 92.59% | 165 |
| Eruve.0031s0115 | Eruve.0031s0115.1 | | 100% | 8E-121 | 84.68% | 222 |
| | Eruve.0031s0115.2 | | 100% | 8E-121 | 84.68% | 222 |
| Eruve.1455s0007 | Eruve.1455s0007.1 | | 86% | 2E-111 | 86.46% | 198 |
| | Eruve.1455s0007.2 | | 62% | 2E-84 | 94.81% | 142 |
| Eucgr.F03985 | Eucgr.F03985 | *Eucalyptus grandis* | 92% | 2E-77 | 59.70% | 212 |
| Eucgr.J03130 | Eucgr.J03130 | | 93% | 6E-72 | 62.56% | 208 |
| Eusyr.0118s1180 | Eusyr.0118s1180.1 | *Euclidium syriacum* | 95% | 8E-126 | 88.89% | 217 |
| Thhalv10019123m | Thhalv10019123m | *Eutrema salsugineum* | 95% | 5E-131 | 89.42% | 217 |
| Thhalv10019121m | Thhalv10019121m | | 95% | 5E-131 | 89.42% | 217 |
| FvH4_6g28270 | FvH4_6g28270.t1 | *Fragaria vesca* | 80% | 4E-72 | 67.43% | 209 |
| | FvH4_6g28270.t2 | | 92% | 2E-82 | 66.16% | 211 |
| Glyma.08G220400 | Glyma.08G220400.1 | *Glycine max* | 95% | 3E-91 | 66.67% | 210 |
| Glyma.07G021400 | Glyma.07G021400.1 | | 100% | 5E-90 | 64.19% | 214 |
| Gohir.A11G075200 | Gohir.A11G075200.1 | *Gossypium hirsutum* | 91% | 5E-88 | 69.39% | 214 |
| Gohir.D11G079900 | Gohir.D11G079900.1 | | 81% | 1E-82 | 73.14% | 195 |
| Gorai.007G085500 | Gorai.007G085500.1 | *Gossypium raimondii* | 87% | 1E-88 | 72.34% | 214 |
| | Gorai.007G085500.2 | | 81% | 1E-82 | 73.14% | 195 |
| | Gorai.007G085500.3 | | 87% | 1E-68 | 61.70% | 194 |
| HanXRQChr04g0100021 | HanXRQChr04g0100021 | *Helianthus annuus* | 91% | 1E-86 | 69.23% | 211 |
| HanXRQChr09g0274181 | HanXRQChr09g0274181 | | 92% | 2E-83 | 65.67% | 205 |
| Hyque.02G002700 | Hyque.02G002700.1 | *Hydrangea quercifolia* | 97% | 6E-95 | 64.32% | 210 |
| | Hyque.02G002700.2 | | 97% | 5E-95 | 64.62% | 209 |
| HORVU0Hr1G040430 | HORVU0Hr1G040430.1 | *Hordem vulgare* | 88% | 1E-65 | 57.94% | 351 |
| | HORVU0Hr1G040430.4 | | 88% | 7E-66 | 57.94% | 345 |
| Ibeam.0002s0036 | Ibeam.0002s0036.1 | *Iberis amara* | 95% | 4E-103 | 80.95% | 214 |
| | Ibeam.0002s0036.2 | | 95% | 2E-99 | 80.00% | 209 |
| Isati.5657s0005 | Isati.5657s0005.1 | *Isatis tinctoria* | 99% | 7E-130 | 87.91% | 214 |
| Joasc.11G051500 | Joasc.11G051500.1 | *Joinvillea ascendens* | 90% | 9E-76 | 61.39% | 204 |
| | Joasc.11G051500.2 | | 90% | 9E-76 | 61.39% | 204 |
| | Joasc.11G051500.3 | | 63% | 8E-72 | 76.98% | 158 |
| | Joasc.11G051500.4 | | 33% | 1E-37 | 84.72% | 96 |
| Kaladp0335s0001 | Kaladp0335s0001.1 | *K. fedtschenkoi* | 87% | 4E-76 | 63.16% | 233 |
| Kaladp0037s0373 | Kaladp0037s0373.1 | | 81% | 5E-74 | 66.29% | 247 |
| | Kaladp0037s0373.2 | | 61% | 7E-73 | 77.04% | 180 |
| | Kaladp0037s0373.3 | | 62% | 2E-71 | 76.47% | 243 |
| | Kaladp0037s0373.4 | | 61% | 7E-73 | 77.04% | 180 |
| | Kaladp0037s0373.5 | | 61% | 7E-73 | 77.04% | 180 |

| | | | | | | |
|---|---|---|---|---|---|---|
| Lsat_1_v5_gn_1_30801 | Lsat_1_v5_gn_1_30801.1 | *Lactuca sativa* | 92% | 1E-89 | 64.79% | 213 |
| | Lsat_1_v5_gn_1_30801.2 | | 95% | 4E-91 | 66.51% | 212 |
| Lsat_1_v5_gn_4_85061 | Lsat_1_v5_gn_4_85061.1 | | 97% | 1E-87 | 61.19% | 216 |
| Lcu.2RBY.4g001240 | Lcu.2RBY.4g001240.1 | *Lens culinaris* | 93% | 3E-82 | 64.50% | 210 |
| Lesat.0180s0006 | Lesat.0180s0006.1 | *Lepidium sativum* | 100% | 4E-130 | 87.33% | 219 |
| | Lesat.0180s0006.2 | | 100% | 1E-132 | 88.94% | 215 |
| | Lesat.0180s0006.3 | | 100% | 1E-132 | 88.94% | 215 |
| Lesat.0061s0110 | Lesat.0061s0110.1 | | 95% | 1E-125 | 87.74% | 220 |
| Liphi.05G091000 | Liphi.05G091000.1 | *Lindenbergia philippensis* | 95% | 4E-81 | 63.85% | 215 |
| | Liphi.05G091000.2 | | 71% | 3E-68 | 73.03% | 164 |
| Lus10041624 | Lus10041624 | *Linum usitatissimum* | 86% | 1E-78 | 64.80% | 236 |
| Lus10024095 | Lus10024095 | | 86% | 3E-77 | 63.27% | 321 |
| LitulAlt.12G037200 | LitulAlt.12G037200 | *Liriodendron tulipifera* | 91% | 4E-75 | 61.46% | 215 |
| Lj3g0027435 | Lj3g0027435.1 | *Lotus japonicus* | 96% | 7E-87 | 61.29% | 215 |
| Luann.0047s0143 | Luann.0047s0143.1 | *Lunaria annua* | 95% | 7E-130 | 90.34% | 216 |
| Luann.1171s0004 | Luann.1171s0004.1 | | 62% | 1E-76 | 89.47% | 152 |
| Lalb_Chr01g0011991 | Lalb_Chr01g0011991 | *Lupinus albus* | 93% | 1E-81 | 61.81% | 206 |
| Lalb_Chr25g0278731 | Lalb_Chr25g0278731 | | 89% | 6E-81 | 65.10% | 206 |
| Mamar.0107s0005 | Mamar.0107s0005.1 | *Malcolmia maritima* | 100% | 6E-135 | 92.49% | 210 |
| Manes.11G069200 | Manes.11G069200 | *Manihot esculenta* | 98% | 7E-90 | 68.25% | 220 |
| Manes.04G100900 | Manes.04G100900 | | 90% | 1E-82 | 67.01% | 202 |
| Mapoly0128s0009 | Mapoly0128s0009 | *Marchantia polymorpha* | 61% | 1E-62 | 69.23% | 243 |
| Medtr4g007700 | Medtr4g007700.1 | *Medicago truncatula* | 93% | 7E-88 | 64.93% | 209 |
| | Medtr4g007700.2 | | 81% | 5E-78 | 62.76% | 195 |
| MgTOL.I0689 | MgTOL.I0689.1 | *Mimulus guttatus* | 91% | 1E-82 | 60.09% | 223 |
| Misin07G522700 | Misin07G522700 | *Miscanthus sinensis* | 89% | 3E-66 | 60.30% | 321 |
| Ma03_g04600 | Ma03_g04600 | *Musa acuminata* | 86% | 6E-70 | 57.87% | 204 |
| Myper.0013s0857 | Myper.0013s0857.1 | *Myagrum perfoliatum* | 99% | 1E-130 | 88.37% | 214 |
| NbS00012287g0005 | NbS00012287g0005 | *Nicotiana benthamiana* | 90% | 4,00E-87 | 63.92% | 209 |
| NbS00042944g0008 | NbS00042944g0008 | | 77% | 1,00E-78 | 66.87% | 184 |
| Nycol.C02206 | Nycol.C02206.1 | *Nymphaea colorata* | 91% | 3,00E-82 | 60.58% | 222 |
| | Nycol.C02206.2 | | 65% | 6,00E-47 | 54.90% | 178 |
| Nitab4.5_0004738g0090 | Nitab4.5_0004738g0090 | *Nicotiana tabacum* | 94% | 9,00E-89 | 63.37% | 215 |
| Nitab4.5_0002700g0060 | Nitab4.5_0002700g0060 | | 90% | 3,00E-88 | 65.80% | 208 |
| Oeu051555 | Oeu051555.1 | *Olea europaea* | 74% | 5E-63 | 67.70% | 156 |
| Oeu003125 | Oeu003125.1 | | 65% | 8E-62 | 73.43% | 265 |
| Oropetium_20150105_04478 | Oropetium_20150105_04478A | *Oropetium thomaeum* | 81% | 4E-63 | 61.17% | 576 |
| LOC_Os02g55080 | LOC_Os02g55080 | *Oryza sativa* | 86% | 5E-70 | 60.70% | 210 |
| Pahal.1G428900 | Pahal.1G428900.1 | *Panicum hallii* | 89% | 2E-70 | 60.29% | 207 |
| PhHAL.1G419700 | PhHAL.1G419700.1 | *Panicum hallii HAL* | 89% | 2E-70 | 60.29% | 207 |
| Pavir.1NG509700 | Pavir.1NG509700.2 | *Panicum virgatum* | 89% | 1E-69 | 59.90% | 207 |
| Pavir.1KG533800 | Pavir.1KG533800.1 | | 89% | 2E-68 | 58.82% | 207 |
| | Pavir.1KG533800.2 | | 59% | 2E-37 | 54.74% | 183 |
| Pavag04G304200 | Pavag04G304200.1 | *Paspalum vaginatum* | 89% | 8E-67 | 59.90% | 207 |
| Phala.02G314500 | Phala.02G314500.1 | *Pharus latifolius* | 89% | 5E-69 | 58.25% | 210 |
| | Phala.02G314500.2 | | 61% | 5,00E-40 | 52.78% | 153 |
| Phacu.CVR.008G317800 | Phacu.CVR.008G317800.1 | *Phaseolus acutifolius* | 95% | 1,00E-90 | 64.56% | 209 |
| Phvul.008G253200 | Phvul.008G253200.1 | *Phaseolus vulgaris* | 95% | 3,00E-91 | 65.05% | 209 |
| Pp3c16_25700 | Pp3c16_25700V3.1 | *Physcomitrium patens* | 61% | 9,00E-62 | 66.91% | 339 |
| | Pp3c16_25700V3.2 | | 61% | 9,00E-62 | 66.91% | 339 |
| Pp3c25_9780 | Pp3c25_9780V3.1 | | 61% | 3,00E-60 | 66.18% | 338 |
| | Pp3c25_9780V3.2 | | 61% | 3,00E-60 | 66.18% | 339 |
| Psat7g262200 | Psat7g262200 | *Pisum sativum* | 93% | 6E-88 | 64.88% | 210 |
| Ptrif.0007s1735 | Ptrif.0007s1735.1 | *Poncirus trifoliata* | 97% | 1E-75 | 61.61% | 216 |
| | Ptrif.0007s1735.2 | | 97% | 7E-74 | 60.48% | 207 |
| | Ptrif.0007s1735.3 | | 66% | 6E-66 | 75.69% | 145 |
| Podel.03G199500 | Podel.03G199500.1 | *Populus deltoides* | 95% | 5E-88 | 68.29% | 220 |
| Podel.01G043400 | Podel.01G043400.1 | | 95% | 1E-86 | 66.83% | 221 |
| | Podel.01G043400.2 | | 90% | 1E-75 | 63.21% | 207 |
| | Podel.01G043400.3 | | 63% | 3E-72 | 80.00% | 152 |
| Poman.03G163300 | Poman.03G163300.1 | *Populus nigra x maximowiczii* | 95% | 1E-88 | 68.78% | 221 |
| | Poman.03G163300.2 | | 90% | 7E-78 | 65.28% | 207 |
| | Poman.03G163300.3 | | 63% | 2E-72 | 80.74% | 152 |
| Poman.01G035900 | Poman.01G035900.1 | | 91% | 5E-87 | 70.26% | 214 |
| | Poman.01G035900.2 | | 90% | 7E-78 | 65.28% | 207 |

Table S3. ABAP1 orthologs in other plant species

| Gene ID | Isoforms | Scientific Name | Query Cover | E value | Per. Ident | Acc. Len |
|---|---|---|---|---|---|---|
| AT5G13060 | AT5G13060.1 | Arabidopsis thaliana | 100% | 0 | 100% | 737 |
| Sobic.009G129600 | Sobic.009G129600.1 | Sorghum bicolor | 88% | 0.0 | 60.79% | 745 |
| Sspon.07G0008510-2C | Sspon.07G0008510-2C | Saccharum spontaneum | 79% | 0.0 | 58.38% | 622 |
| Zm00001eb287530 | Zm00001eb287530 | Zea mays B73 | 88% | 0.0 | 59.69% | 724 |
| Zm00001eb349910 | Zm00001eb349910 | | 87% | 0.0 | 60.74% | 748 |
| OsR498G0510791000 | OsR498G0510791000.01 | Oryza sativa | 89% | 0.0 | 60.76% | 745 |
| TraesCS1A03G0645500 | TraesCS1A03G0645500 | Triticum aestivum | 88% | 0.0 | 60.37% | 730 |
| TraesCS1B03G0731200 | TraesCS1B03G0731200 | | 89% | 0.0 | 60.15% | 742 |
| TraesCS1D03G0602500 | TraesCS1D03G0602500 | | 89% | 0.0 | 60.15% | 776 |
| Horvu_MOREX_1H01G392500 | Horvu_MOREX_1H01G392500 | Hordeum vulgare | 89% | 0.0 | 59.67% | 742 |
| Ma09_g16490 | Ma09_g16490.1 | Musa acuminata | 89% | 0.0 | 61.93% | 726 |
| Cc01_g19440 | Cc01_g19440.1 | Coffea canephora | 95% | 0.0 | 60.85% | 712 |
| evm.TU.Scaffold_557.287 | evm.model.Scaffold_557.287 | Coffea arabica | 95% | 0.0 | 58.96% | 692 |
| evm.TU.Scaffold_2016.221 | evm.model.Scaffold_2016.221 | | 95% | 0.0 | 57.94% | 717 |
| GSVIVG01011129001 | GSVIVG01011129001 | Vitis vinífera | 93% | 0.0 | 59.43% | 713 |
| GSVIVG01017925001 | GSVIVG01017925001 | | 89% | 0.0 | 63.47% | 705 |
| PGSC0003DMG400012880 | PGSC0003DMT400033530 | Solanum tuberosum | 95% | 0.0 | 59.83% | 708 |
| PGSC0003DMG401020126 | PGSC0003DMT400051860 | | 95% | 0.0 | 59.86% | 709 |
| Solyc09G001948 | Solyc09G001948.1 | Solanum lycopersicum | 95% | 0.0 | 59.92% | 708 |
| Solyc06G002747 | Solyc06T002747.1 | | 95% | 0.0 | 50.86% | 734 |
| | Solyc06T002747.2 | | 95% | 0.0 | 51.88% | 720 |
| Manes.03G181400 | Manes.03G181400.1 | Manihot esculenta | 87% | 0.0 | 63.79% | 708 |
| Manes.05G076800 | Manes.05G076800.4 | | 91% | 0.0 | 62.61% | 692 |
| Manes.15G026400 | Manes.15G026400.3 | | 94% | 0.0 | 61.17% | 707 |
| A03p09430 | A03p09430 | Brassica napus | 92% | 0.0 | 59.33% | 716 |
| C09p56950 | C09p56950 | | 88% | 0.0 | 60.09% | 1266 |
| C03p11010 | C03p11010 | | 82% | 0.0 | 50.49% | 554 |
| A10p25140 | A10p25140 | | 96% | 0.0 | 85.09% | 707 |
| C09p64040 | C09p64040 | | 96% | 0.0 | 79.18% | 755 |
| C02p05870 | C02p05870 | | 99% | 0.0 | 81.66% | 729 |
| A02p01660 | A02p01660 | | 96% | 0.0 | 80.54% | 756 |
| BolC2t06532H | BolC2t06532H | Brassica oleracea | 99% | 0.0 | 81.79% | 729 |
| BolC9t58549H | BolC9t58549H | | 88% | 0.0 | 60.09% | 1267 |
| BraA02t05145Z | BraA02t05145Z | Brassica rapa | 96% | 0.0 | 83.15% | 733 |
| BraA03t10261Z | BraA03t10261Z | | 94% | 0.0 | 58.36% | 716 |
| BraA10t44106Z | BraA10t44106Z | | 88% | 0.0 | 61.14% | 1296 |
| BraA10t44576Z | BraA10t44576Z | | 96% | 0.0 | 84.67% | 707 |
| BcaB01g05140 | BcaB01g05140 | Brassica carinata | 94% | 0.0 | 56.01% | 700 |
| BcaB08g36582 | BcaB08g36582 | | 94% | 0.0 | 58.83% | 711 |
| BcaC04g18577 | BcaC04g18577 | | 88% | 0.0 | 62.02% | 715 |
| BcaNung01177 | BcaNung01177 | | 99% | 0.0 | 83.09% | 737 |
| BcaNung01857 | BcaNung01857 | | 99% | 0.0 | 81.52% | 729 |
| Phvul.001G151300 | Phvul.001G151300.1 | Phaseolus vulgaris | 88% | 0.0 | 53.50% | 704 |
| | Phvul.002G047400.3 | | 89% | 0.0 | 62.22% | 707 |
| | Phvul.002G047400.4 | | 0,89 | 0.0 | 62.22% | 707 |
| Phvul.006G001300 | Phvul.006G001300.1 | | 87% | 4E-176 | 43.82% | 699 |
| Phvul.007G054500 | Phvul.007G054500.1 | | 94% | 0.0 | 60.40% | 706 |
| Glyma.01G239200 | Glyma.01G239200.1 | Glycine max | 92% | 0.0 | 60.70% | 707 |
| | Glyma.01G239200.3 | | 92% | 0.0 | 60.70% | 706 |
| Glyma.03G154000 | Glyma.03G154000.1 | | 89% | 0.0 | 53.76% | 705 |
| Glyma.10G250100 | Glyma.10G250100.2 | | 89% | 0.0 | 61.88% | 707 |
| | Glyma.10G250100.4 | | 89% | 0.0 | 61.88% | 708 |
| Glyma.11G004300 | Glyma.11G004300.1 | | 92% | 0.0 | 61.08% | 708 |
| | Glyma.11G004300.3 | | 92% | 0.0 | 61.08% | 709 |
| Glyma.19G156400 | Glyma.19G156400.1 | | 88% | 0.0 | 54.67% | 714 |
| | Glyma.19G156400.2 | | 88% | 0.0 | 55.41% | 704 |
| Glyma.20G143500 | Glyma.20G143500.1 | | 89% | 0.0 | 61.72% | 707 |
| | Glyma.20G143500.2 | | 89% | 0.0 | 60.06% | 686 |
| Psat6g208840 | Psat6g208840.1 | Pisum sativum | 94% | 0.0 | 59.80% | 704 |
| Psat4g211360 | Psat4g211360.1 | | 88% | 0.0 | 61.98% | 703 |
| Psat3g064160 | Psat3g064160.1 | | 93% | 0.0 | 56.31% | 735 |
| Eucgr.H03553 | Eucgr.H03553 | Eucalyptus grandis | 89% | 0.0 | 62.18% | 712 |
| Thecc.01G278800 | Thecc.01G278800.1 | Theobroma cacao | 94% | 0.0 | 58.98% | 705 |
| Thecc.04G227100 | Thecc.04G227100.1 | | 89% | 0.0 | 63.17% | 704 |

| | | | | | | |
|---|---|---|---|---|---|---|
| Gorai.005G041500 | Gorai.005G041500.1 | Gossypium raimondii | 92% | 0.0 | 61.08% | 705 |
| Gorai.006G237100 | Gorai.006G237100.1 | | 89% | 0.0 | 61.95% | 704 |
| | Gorai.006G237100.2 | | 89% | 0.0 | 59.82% | 681 |
| | Gorai.006G237100.3 | | 85% | 0.0 | 63.13% | 687 |
| | Gorai.006G237100.4 | | 86% | 0.0 | 67.00% | 513 |
| | Gorai.006G237100.5 | | 86% | 0.0 | 62.60% | 697 |
| Lsat_1_v5_gn_5_188180 | Lsat_1_v5_gn_5_188180.1 | Lactuca sativa | 95% | 0.0 | 61.16% | 757 |
| | Lsat_1_v5_gn_5_188180.2 | | 95% | 0.0 | 61.16% | 757 |
| Lsat_1_v5_gn_9_3640 | Lsat_1_v5_gn_9_3640.3 | | 94% | 0.0 | 59.86% | 726 |
| Lsat_1_v5_gn_3_84821 | Lsat_1_v5_gn_3_84821.1 | | 37% | 1E-54 | 53.97% | 239 |
| Cpa.g.sc147.32 | Cpa.t.sc147.32 | Carica papaya | 92% | 0.0 | 60.47% | 715 |
| Cpa.g.sc370.9 | Cpa.t.sc370.9 | | 93% | 0.0 | 61.99% | 704 |
| 29844.t000205 | 29844.m003359 | Ricinus communis | 89% | 0.0 | 63.13% | 719 |
| 29647.t000089 | 29647.m002081 | | 84% | 0.0 | 60.09% | 598 |
| Ciclev10004440m.g | Ciclev10004440m | Citrus clementina | 89% | 0.0 | 62.86% | 717 |
| Ciclev10019087m.g | Ciclev10019087m | | 94% | 0.0 | 61.69% | 704 |
| orange1.1g005282m.g | orange1.1g005282m | Citrus sinensis | 94% | 0.0 | 61.69% | 704 |
| orange1.1g004992m.g | orange1.1g005044m | | 89% | 0.0 | 62.58% | 720 |
| | orange1.1g004992m | | 89% | 0.0 | 62.58% | 720 |
| | orange1.1g005088m | | 89% | 0.0 | 62.25% | 715 |
| | orange1.1g005144m | | 89% | 0.0 | 62.25% | 712 |
| | orange1.1g007101m | | 75% | 0.0 | 64.80% | 618 |
| | orange1.1g007104m | | 75% | 0.0 | 64.80% | 618 |
| | orange1.1g008781m | | 66% | 0.0 | 63.21% | 554 |
| | orange1.1g008940m | | 86% | 0.0 | 67.67% | 548 |
| | orange1.1g010291m | | 86% | 0.0 | 67.59% | 513 |
| | orange1.1g015851m | | 49% | 5E-123 | 62.78% | 399 |
| Misin17G131600 | Misin17G131600.1 | Miscanthus sinensis | 88% | 0.0 | 61.01% | 732 |
| | Misin17G131600.2 | | 88% | 0.0 | 60.31% | 717 |
| Misin16G132500 | Misin16G132500.1 | | 88% | 0.0 | 60.55% | 744 |
| EL10Ac7g17234 | EL10Ac7g17234.1 | Beta vulgaris | 94% | 0.0 | 56.13% | 744 |
| EL10As3g23353 | EL10As3g23353.1 | | 88% | 0.0 | 58.74% | 714 |
| Bobra.126_2s0003 | Bobra.126_2s0003.1 | Botryococcus braunii | 94% | 0.0 | 47.89% | 707 |
| Cre04.g231516 | Cre04.g231516.t1.1 | Chlamydomonas reinhardtii | 94% | 0.0 | 47.89% | 707 |
| | Cre04.g231516.t2.1 | | 84% | 4E-179 | 45.17% | 763 |
| UNPLg00364 | UNPLg00364.t1 | Chromochloris zofingiensis | 83% | 0.0 | 47.10% | 677 |
| Araha.3287s0012 | Araha.3287s0012.1 | Arabidopsis halleri | 85% | 0.0 | 93.51% | 634 |
| Araha.13116s0005 | Araha.13116s0005.1 | | 94% | 0.0 | 60.51% | 711 |
| | Araha.13116s0005.2 | | 94% | 0.0 | 60.46% | 711 |
| Bostr.2902s0040 | Bostr.2902s0040.1.p | Boecherastricta | 100% | 0.0 | 91.47% | 739 |
| Bostr.26527s0273 | Bostr.26527s0273.1 | | 94% | 0.0 | 60.63% | 710 |
| Bostr.2570s0062 | Bostr.2570s0062.1 | | 85% | 0.0 | 60.14% | 439 |
| Mamar.0011s0326 | Mamar.0011s0326.1 | Malcolmia maritima | 100% | 0.0 | 89.99% | 739 |
| Mamar.0050s1498 | Mamar.0050s1498.1 | | 94% | 0.0 | 60.46% | 710 |
| Mamar.0016s0220 | Mamar.0016s0220.1 | | 94% | 0.0 | 55.71% | 698 |
| Alyli.0206s0053 | Alyli.0206s0053.1 | Alyssum linifolium | 96% | 0.0 | 88.89% | 707 |
| Alyli.0096s0006 | Alyli.0096s0006.1 | | 94% | 0.0 | 59.77% | 710 |
| Alyli.0053s0226 | Alyli.0053s0226.1 | | 94% | 0.0 | 59.77% | 710 |
| | Alyli.0053s0226.2 | | 94% | 0.0 | 59.83% | 711 |
| Thlar.0013s1177 | Thlar.0013s1177.1 | Thlaspi arvense | 96% | 0.0 | 85.65% | 710 |
| Thlar.0007s0309 | Thlar.0007s0309.1 | | 94% | 0.0 | 60.83% | 710 |
| AL6G23660 | AL6G23660.t1 | Arabidopsis lyrata | 96% | 0.0 | 92.72% | 713 |
| AL6G30820 | AL6G30820.t1 | | 94% | 0.0 | 60.80% | 710 |
| | AL6G30820.t2 | | 94% | 0.0 | 60.85% | 712 |
| AL3G16810 | AL3G16810.t1 | | 88% | 0.0 | 57.25% | 694 |
| | AL3G16810.t2 | | 88% | 0.0 | 57.25% | 694 |
| Aqcoe2G339500 | Aqcoe2G339500.1 | Aquilegia coerulea | 89% | 0.0 | 57.87% | 757 |
| AgateH1.02G001300 | AgateH1.02G001300.1 | Agave tequilana var. Weber's Blue | 89% | 0.0 | 58.90% | 744 |
| | AgateH1.02G001300.2 | | 89% | 0.0 | 58.90% | 744 |
| | AgateH1.02G001300.2 | | 89% | 0.0 | 61.67% | 714 |
| | AgateH1.02G001300.3 | | 89% | 0.0 | 58.32% | 739 |
| | AgateH1.02G001300.4 | | 89% | 0.0 | 58.03% | 737 |
| AgateH1.02G344000 | AgateH1.02G344000.1 | | 89% | 0.0 | 60.82% | 712 |
| Alyli.0206s0053 | Alyli.0206s0053.1 | Alyssum linifolium | 96% | 0.0 | 88.89% | 707 |
| Alyli.0096s0006 | Alyli.0096s0006.1 | | 94% | 0.0 | 59.77% | 710 |
| Alyli.0053s0226 | Alyli.0053s0226.1 | | 94% | 0.0 | 59.77% | 710 |
| | Alyli.0053s0226.2 | | 94% | 0.0 | 59.83% | 711 |
| evm_27.model.AmTr_v1.0_scaffold00025.384 | evm_27.model.AmTr_v1.0_scaffold00025.38 | Amborella trichopoda | 87% | 0.0 | 59.63% | 717 |
| AmTrH2.01G163500 | AmTrH2.01G163500.1 | | 87% | 0.0 | 59.63% | 717 |
| Anaoc.0004s0117 | Anaoc.0004s0117.7 | Anacardium occidentale | 77% | 0.0 | 70.98% | 451 |
| | Anaoc.0004s0117.1 | | 89% | 0.0 | 62.71% | 714 |
| | Anaoc.0004s0117.2 | | 86% | 0.0 | 67.79% | 513 |
| | Anaoc.0004s0117.3 | | 86% | 0.0 | 67.79% | 513 |
| | Anaoc.0004s0117.4 | | 86% | 0.0 | 67.79% | 513 |
| | Anaoc.0004s0117.5 | | 77% | 0.0 | 70.98% | 452 |
| | Anaoc.0004s0117.6 | | 77% | 0.0 | 70.98% | 452 |
| | Anaoc.0004s0117.7 | | 77% | 0.0 | 70.98% | 452 |
| | Anaoc.0004s0117.8 | | 89% | 0.0 | 62.71% | 714 |
| | Anaoc.0004s0117.9 | | 80% | 0.0 | 64.53% | 653 |
| | Anaoc.0004s0117.10 | | 75% | 0.0 | 64.09% | 617 |
| | Anaoc.0004s0117.11 | | 89% | 0.0 | 62.71% | 714 |
| | Anaoc.0004s0117.12 | | 80% | 0.0 | 64.53% | 653 |
| | Anaoc.0004s0117.13 | | 65% | 0.0 | 63.43% | 579 |
| Anaoc.0004s0821 | Anaoc.0004s0821.1 | | 85% | 0.0 | 65.34% | 587 |
| Anaoc.0008s0482 | Anaoc.0008s0482.1 | | 89% | 0.0 | 62.97% | 715 |
| Aco006295 | Aco006295.1 | Ananas comosus | 90% | 0.0 | 59.31% | 706 |

| | | | | | | |
|---|---|---|---|---|---|---|
| AndgeH2.09AG127100 | AndgeH2.09AG127100.2 | Andropogon Gerardi | 0% | 0 | 65.00% | 0 |
| | AndgeH2.09AG127100.1 | | 71% | 0.0 | 62.62% | 615 |
| | AndgeH2.09AG127100.3 | | 0% | 0 | 63.00% | 0 |
| AndgeH2.09BG107700 | AndgeH2.09BG107700.7 | | 0% | 0 | 64.00% | 0 |
| | AndgeH2.09BG107700.6 | | 0% | 0 | 64.00% | 0 |
| | AndgeH2.09BG107700.4 | | 0% | 0 | 63.00% | 0 |
| | AndgeH2.09BG107700.5 | | 0% | 0 | 64.00% | 0 |
| | AndgeH2.09BG107700.2 | | 0% | 0 | 63.00% | 0 |
| | AndgeH2.09BG107700.1 | | 87% | 0.0 | 63.94% | 593 |
| | AndgeH2.09BG107700.3 | | 0% | 0 | 63.00% | 0 |
| AndgeH2.09CG179600 | AndgeH2.09CG179600.1 | | 89% | 0.0 | 60.73% | 736 |
| arahy.Tifrunner.gnm1.ann1.7RR6EE | arahy.Tifrunner.gnm1.ann1.7RR6EE.1 | Arachis hypogaea | 62% | 0.0 | 64.43% | 437 |
| | arahy.Tifrunner.gnm1.ann1.7RR6EE.2 | | 0% | 0 | 0,64 | 0 |
| arahy.Tifrunner.gnm1.ann1.C0GFHN | arahy.Tifrunner.gnm1.ann1.C0GFHN.1 | | 88% | 0.0 | 59.02% | 685 |
| arahy.Tifrunner.gnm1.ann1.CB6P6Z | arahy.Tifrunner.gnm1.ann1.CB6P6Z.1 | | 94% | 0.0 | 57.38% | 724 |
| arahy.Tifrunner.gnm1.ann1.71BXTT | arahy.Tifrunner.gnm1.ann1.71BXTT.1 | | 94% | 0.0 | 56.83% | 730 |
| arahy.Tifrunner.gnm1.ann1.825X9M | arahy.Tifrunner.gnm1.ann1.825X9M.1 | | 93% | 0.0 | 56.80% | 699 |
| arahy.Tifrunner.gnm1.ann1.GF6ZB7 | arahy.Tifrunner.gnm1.ann1.GF6ZB7.1 | | 93% | 0.0 | 56.94% | 699 |
| | arahy.Tifrunner.gnm1.ann1.GF6ZB7.2 | | 93% | 0.0 | 56.94% | 699 |
| evm.TU.AsparagusV1_07.1797 | evm.model.AsparagusV1_07.1797 | Asparagus officinalis | 94% | 0.0 | 57.79% | 714 |
| BPChr05G08852 | BPChr05G08852 | Betula platyphylla | 84% | 0.0 | 64.84% | 715 |
| BPChr05G08804 | BPChr05G08804 | | 91% | 0.0 | 61.98% | 712 |
| BPChr08G05060 | BPChr08G05060 | | 84% | 0.0 | 51.90% | 656 |
| Bostr.2902s0040 | Bostr.2902s0040.1 | Boechera stricta | 100% | 0.0 | 91.47% | 739 |
| Bostr.26527s0273 | Bostr.26527s0273.1 | | 94% | 0.0 | 60.63% | 710 |
| Bostr.2570s0062 | Bostr.2570s0062.1 | | 85% | 0.0 | 60.14% | 439 |
| Barbu.8G135600 | Barbu.8G135600.1 | Brachypodium arbuscula | 88% | 0.0 | 60.06% | 740 |
| Brahy.S08G0135700 | Brahy.S08G0135700.1 | Brachypodium hybridum | 89% | 0.0 | 59.85% | 734 |
| Brahy.D02G0363000 | Brahy.D02G0363000.1 | | 88% | 0.0 | 60.21% | 787 |
| Brasyl.8G132300 | Brasyl.8G132300.1 | Brachypodium sylvaticum | 88% | 0.0 | 60.06% | 739 |
| | Brasyl.8G132300.2 | | 84% | 0.0 | 61.46% | 718 |
| | Brasyl.8G132300.3 | | 84% | 0.0 | 61.08% | 722 |
| Brast08G128100 | Brast08G128100.1 | Brachypodium stacei | 88% | 0.0 | 60.21% | 739 |
| Brmexi.08PG175100 | Brmexi.08PG175100.1 | Brachypodium mexicanum | 84% | 0.0 | 60.95% | 773 |
| | Brmexi.08PG175100.2 | | 88% | 0.0 | 60.06% | 734 |
| | Brmexi.08PG175100.3 | | 88% | 0.0 | 59.91% | 732 |
| | Brmexi.08PG175100.4 | | 88% | 0.0 | 58.99% | 725 |
| | Brmexi.08PG175100.5 | | 88% | 0.0 | 58.20% | 755 |
| Brmexi.08UG146100 | Brmexi.08UG146100.1 | | 88% | 0.0 | 56.87% | 759 |
| | Brmexi.08UG146100.2 | | 88% | 0.0 | 59.60% | 731 |
| Camar.0268s0010 | Camar.0268s0010.1 | Cakile maritima | 95% | 0.0 | 84.41% | 741 |
| Camar.0675s0029 | Camar.0675s0029.1 | | 96% | 0.0 | 85.21% | 710 |
| Camar.0558s0016 | Camar.0558s0016.1 | | 94% | 0.0 | 60.00% | 712 |
| Cagra.1535s0001 | Cagra.1535s0001.1 | Capsella grandiflora | 95% | 0.0 | 83.45% | 716 |
| Cagra.4243s0018 | Cagra.4243s0018.1 | | 94% | 0.0 | 60.83% | 712 |
| Cagra.4243s0018 | Cagra.4243s0018.2 | | 94% | 0.0 | 60.77% | 711 |
| Caril.07G091200 | Caril.07G091200.1 | Carya illinoinensis | 89% | 0.0 | 62.48% | 717 |
| Caril.01G159600 | Caril.01G159600.1 | | 83% | 0.0 | 64.72% | 760 |
| | Caril.01G159600.2 | | 83% | 0.0 | 64.72% | 760 |
| | Caril.01G159600.3 | | 94% | 0.0 | 60.60% | 705 |
| | Caril.01G159600.4 | | 94% | 0.0 | 60.60% | 705 |
| | Caril.01G159600.5 | | 91% | 0.0 | 61.37% | 683 |
| Caril.02G095600 | Caril.02G095600.1 | | 87% | 0.0 | 60.80% | 730 |
| | Caril.02G095600.2 | | 87% | 0.0 | 63.33% | 704 |
| | Caril.02G095600.3 | | 87% | 0.0 | 63.33% | 704 |
| | Caril.02G095600.4 | | 87% | 0.0 | 63.33% | 704 |
| | Caril.02G095600.5 | | 87% | 0.0 | 63.33% | 704 |
| | Caril.02G095600.6 | | 87% | 0.0 | 61.02% | 681 |
| | Caril.02G095600.7 | | 87% | 0.0 | 62.94% | 708 |
| Caden.10G092800 | Caden.10G092800.1 | Castanea dentata | 93% | 0.0 | 62.03% | 708 |
| Caden.02G024900 | Caden.02G024900.1 | | 88% | 0.0 | 63.68% | 735 |
| CmMahoganyH1.10G076000 | CmMahoganyH1.10G076000.1 | Castanea mollissima Mahogany | 93% | 0.0 | 62.03% | 708 |
| | CmMahoganyH1.10G076000.2 | | 93% | 0.0 | 61.60% | 703 |
| CmMahoganyH1.02G020700 | CmMahoganyH1.02G020700.1 | | 88% | 0.0 | 63.68% | 735 |
| Caamp.1041s1256 | Caamp.1041s1256.1 | Caulanthus amplexicaulis | 99% | 0.0 | 86.18% | 736 |
| Caamp.1037s0435 | Caamp.1037s0435.1 | | 94% | 0.0 | 59.83% | 711 |
| | Caamp.1037s0435.2 | | 94% | 0.0 | 59.77% | 711 |
| Caamp.0105s0848 | Caamp.0105s0848.1 | | 94% | 0.0 | 59.69% | 711 |
| | Caamp.0105s0848.2 | | 94% | 0.0 | 59.63% | 711 |
| CepurGG1.3G150100 | CepurGG1.3G150100.1 | Ceratodon purpureus GG1 | 95% | 0.0 | 54.41% | 722 |
| | CepurGG1.3G150100.2 | | 95% | 0.0 | 56.03% | 702 |
| | CepurGG1.3G150100.3 | | 91% | 0.0 | 56.87% | 680 |
| | CepurGG1.3G150100.4 | | 95% | 0.0 | 55.79% | 705 |
| | CepurGG1.3G150100.5 | | 95% | 0.0 | 54.63% | 720 |
| | CepurGG1.3G150100.6 | | 81% | 0.0 | 57.62% | 596 |
| | CepurGG1.3G150100.7 | | 81% | 0.0 | 54.21% | 634 |
| | CepurGG1.3G150100.8 | | 81% | 0.0 | 57.90% | 593 |
| | CepurGG1.3G150100.9 | | 78% | 0.0 | 55.95% | 590 |
| | CepurGG1.3G150100.10 | | 75% | 0.0 | 53.01% | 584 |
| Ceric.36G016000 | Ceric.36G016000.1 | Ceratopteris richardii | 92% | 0.0 | 58.33% | 711 |
| | Ceric.36G016000.2 | | 92% | 0.0 | 58.42% | 711 |
| Cecan.1G431700 | Cecan.1G431700.1 | Cercis canadenses | 95% | 0.0 | 61.69% | 742 |
| | Cecan.1G431700.2 | | 93% | 0.0 | 62.50% | 711 |
| Cecan.2G164300 | Cecan.2G164300.1 | | 94% | 0.0 | 61.01% | 706 |
| | Cecan.2G164300.2 | | 94% | 0.0 | 61.01% | 707 |
| | Cecan.2G164300.3 | | 94% | 0.0 | 61.01% | 707 |
| | Cecan.2G164300.4 | | 94% | 0.0 | 61.01% | 707 |
| | Cecan.2G164300.5 | | 94% | 0.0 | 60.43% | 706 |

| | | | | | | |
|---|---|---|---|---|---|---|
| Chala.07G138200 | Chala.07G138200.1 | Chasmanthium laxum | 88% | 0.0 | 60.88% | 750 |
| Ca_04616 | Ca_04616 | Cicer arietinum | 89% | 0.0 | 61.52% | 705 |
| Ca_07379 | Ca_07379 | | 92% | 0.0 | 57.72% | 732 |
| CKAN_00522800 | CKAN_00522800 | Cinnamomum kanehirae | 94% | 0.0 | 59.68% | 703 |
| CKAN_02407500 | CKAN_02407500 | | 87% | 0.0 | 63.51% | 704 |
| Clevi.0008s1094 | Clevi.0008s1094.1 | Cleome violacea | 96% | 0.0 | 69.15% | 696 |
| Clevi.0003s1966 | Clevi.0003s1966.1 | | 94% | 0.0 | 60.11% | 711 |
| | Clevi.0003s1966.2 | | 94% | 0.0 | 60.11% | 711 |
| CamerWinkler.02G102500 | CamerWinkler.02G102500.1 | Corylus americana | 95% | 0.0 | 60.73% | 710 |
| CamerWinkler.01G086200 | CamerWinkler.01G086200.1 | | 88% | 0.0 | 63.72% | 704 |
| | CamerWinkler.01G086200.2 | | 88% | 0.0 | 61.43% | 682 |
| | CamerWinkler.01G086200.3 | | 88% | 0.0 | 61.43% | 682 |
| | CamerWinkler.01G086200.4 | | 88% | 0.0 | 63.72% | 705 |
| Crahi.0006s0149 | Crahi.0006s0149.1 | Crambe hispanica | 96% | 0.0 | 85.69% | 729 |
| Crahi.0002s0039 | Crahi.0002s0039.1 | | 99% | 0.0 | 84.22% | 728 |
| Crahi.1073s0019 | Crahi.1073s0019.1 | | 94% | 0.0 | 59.94% | 710 |
| Crahi.0488s0065 | Crahi.0488s0065.1 | | 92% | 0.0 | 58.10% | 708 |
| Cucsa.254160 | Cucsa.254160.1 | Cucumis sativus | 88% | 0.0 | 62.80% | 703 |
| | Cucsa.254160.2 | | 86% | 0.0 | 67.13% | 530 |
| DCAR_002731 | DCAR_002731 | Daucus carota | 95% | 0.0 | 58.11% | 703 |
| DCAR_018420 | DCAR_018420 | | 95% | 0.0 | 58.17% | 696 |
| Desop.0240s1381 | Desop.0240s1381.1 | Descurainia sophioides | 96% | 0.0 | 89.17% | 707 |
| Desop.0240s0737 | Desop.0240s0737.1 | | 94% | 0.0 | 59.77% | 710 |
| Desop.0015s0242 | Desop.0015s0242.1 | | 91% | 0.0 | 54.22% | 634 |
| Dioal.07G052900 | Dioal.07G052900.1 | Dioscorea alata | 88% | 0.0 | 62.02% | 718 |
| | Dioal.07G052900.2 | | 88% | 0.0 | 62.02% | 718 |
| Dicom.01G166000 | Dicom.01G166000.1 | Diphasiastrum complanatum | 98% | 0.0 | 54.68% | 708 |
| | Dicom.01G166000.2 | | 93% | 0.0 | 56.52% | 705 |
| | Dicom.01G166000.3 | | 93% | 0.0 | 56.52% | 705 |
| | Dicom.01G166000.5 | | 93% | 0.0 | 56.52% | 705 |
| | Dicom.01G166000.4 | | 93% | 0.0 | 56.52% | 705 |
| | Dicom.01G166000.6 | | 93% | 0.0 | 56.52% | 705 |
| | Dicom.01G166000.7 | | 93% | 0.0 | 56.44% | 706 |
| | Dicom.01G166000.8 | | 93% | 0.0 | 56.44% | 706 |
| Eruve.0575s0004 | Eruve.0575s0004.1 | Eruca vesicaria | 96% | 0.0 | 85.79% | 707 |
| Eruve.0337s0070 | Eruve.0337s0070.1 | | 96% | 0.0 | 85.65% | 707 |
| Eruve.2518s0012 | Eruve.2518s0012.1 | | 99% | 0.0 | 81.72% | 749 |
| Eruve.2532s0001 | Eruve.2532s0001.1 | | 96% | 0.0 | 83.47% | 716 |
| Eruve.3085s0001 | Eruve.3085s0001.1 | | 88% | 0.0 | 62.23% | 717 |
| Eruve.0105s0014 | Eruve.0105s0014.1 | | 88% | 0.0 | 62.23% | 717 |
| Distr.0002s126700 | Distr.0002s126700.1 | Diptychocarpus strictus | 95% | 0.0 | 85.51% | 703 |
| Distr.0002s194500 | Distr.0002s194500.1 | | 94% | 0.0 | 60.68% | 711 |
| | Distr.0002s194500.2 | | 94% | 0.0 | 60.63% | 711 |
| ELECO.r07.5BG0422180 | ELECO.r07.5BG0422180.1 | Eleusine coracana | 85% | 0.0 | 61.89% | 727 |
| ELECO.r07.5AG0374120 | ELECO.r07.5AG0374120.1 | | 85% | 0.0 | 61.57% | 727 |
| Eusyr.0026s0371 | Eusyr.0026s0371.1 | Euclidium syriacum | 96% | 0.0 | 83.83% | 711 |
| Eusyr.0132s0171 | Eusyr.0132s0171.1 | | 94% | 0.0 | 60.06% | 710 |
| Eusyr.0023s0214 | Eusyr.0023s0214.1 | | 88% | 0.0 | 57.32% | 697 |
| Thhalv10012773m.g | Thhalv10012773m | Eutrema salsugineum | 99% | 0.0 | 84.01% | 742 |
| Thhalv10012813m.g | Thhalv10012813m | | 94% | 0.0 | 59.69% | 710 |
| FvH4_4g15040 | FvH4_4g15040.t1 | Fragaria vesca | 85% | 0.0 | 61.99% | 639 |
| | FvH4_4g15040.t2 | | 85% | 0.0 | 61.99% | 770 |
| | FvH4_4g15040.t3 | | 86% | 0.0 | 66.19% | 575 |
| | FvH4_4g15040.t4 | | 94% | 0.0 | 60.60% | 704 |
| | FvH4_4g15040.t5 | | 80% | 0.0 | 61.62% | 705 |
| | FvH4_4g15040.t6 | | 94% | 0.0 | 59.43% | 711 |
| | FvH4_4g15040.t7 | | 91% | 0.0 | 61.52% | 719 |
| FvH4_2g23530 | FvH4_2g23530.t1 | | 90% | 0.0 | 61.50% | 708 |
| | FvH4_2g23530.t2 | | 60% | 1,0E-178 | 62.47% | 513 |
| | FvH4_2g23530.t3 | | 86% | 0.0 | 62.07% | 674 |
| | FvH4_2g23530.t4 | | 80% | 0.0 | 62.92% | 707 |
| | FvH4_2g23530.t5 | | 75% | 0.0 | 62.66% | 652 |
| | FvH4_2g23530.t6 | | 42% | 2,0E-113 | 61.02% | 373 |
| HanXRQChr11g0336141 | HanXRQChr11g0336141 | Helianthus annuus | 95% | 0.0 | 59.46% | 704 |
| HanXRQChr01g0003111 | HanXRQChr01g0003111 | | 95% | 0.0 | 61.31% | 707 |
| Hyque.04G178000 | Hyque.04G178000.1 | Hydrangea quercifolia | 95% | 0.0 | 60.85% | 707 |
| Hyque.10G175000 | Hyque.10G175000.1 | | 94% | 0.0 | 61.00% | 705 |
| Ibeam.3272s0005 | Ibeam.3272s0005.1 | Iberis amara | 94% | 0.0 | 59.60% | 711 |
| | Ibeam.3272s0005.2 | | 94% | 0.0 | 59.55% | 710 |
| Ibeam.2342s0017 | Ibeam.2342s0017.1 | | 94% | 0.0 | 59.55% | 710 |
| Ibeam.1976s0005 | Ibeam.1976s0005.1 | | 95% | 0.0 | 85.15% | 711 |
| Isati.6497s0005 | Isati.6497s0005.1 | Isatis tinctoria | 96% | 0.0 | 86.06% | 707 |
| Isati.7402s0006 | Isati.7402s0006.1 | | 87% | 0.0 | 90.24% | 543 |
| Isati.0539s0031 | Isati.0539s0031.1 | | 96% | 0.0 | 85.92% | 707 |
| Isati.7745s0002 | Isati.7745s0002.1 | | 96% | 0.0 | 85.77% | 707 |
| Joasc.03G088600 | Joasc.03G088600.1 | Joinvillea ascendens | 88% | 0.0 | 61.89% | 733 |
| Kaladp0013s0013 | Kaladp0013s0013.1 | Kalanchoe fedtschenkoi | 94% | 0.0 | 57.59% | 707 |
| Lcu.2RBY.6g052660 | Lcu.2RBY.6g052660.1 | Lens culinaris | 89% | 0.0 | 55.62% | 686 |
| Lcu.2RBY.1g037100 | Lcu.2RBY.1g037100.1 | | 94% | 0.0 | 60.09% | 704 |
| Lcu.2RBY.5g000760 | Lcu.2RBY.5g000760.1 | | 88% | 0.0 | 57.86% | 760 |
| Lesat.0028s0199 | Lesat.0028s0199.1 | Lepidium sativum | 94% | 0.0 | 60.34% | 712 |
| | Lesat.0028s0199.2 | | 94% | 0.0 | 60.28% | 711 |
| Lesat.0059s1052 | Lesat.0059s1052.1 | | 94% | 0.0 | 60.00% | 711 |
| Lesat.0095s0314 | Lesat.0095s0314.1 | | 96% | 0.0 | 87.18% | 710 |
| Liphi.01G183500 | Liphi.01G183500.1 | Lindenbergia philippensis | 95% | 0.0 | 59.44% | 718 |
| | Liphi.01G183500.2 | | 95% | 0.0 | 58.87% | 713 |
| Liphi.11G032600 | Liphi.11G032600.1 | | 93% | 0.0 | 59.68% | 710 |
| | Liphi.11G032600.2 | | 93% | 0.0 | 59.77% | 709 |
| Lus10010527.g | Lus10010527 | Linum usitatissimum | 87% | 0.0 | 63.93% | 717 |
| Lus10022039.g | Lus10022039 | | 94% | 0.0 | 52.57% | 723 |
| Lus10034070.g | Lus10034070 | | 87% | 0.0 | 62.95% | 884 |
| Lus10042583.g | Lus10042583 | | 94% | 0.0 | 53.49% | 709 |
| Lj1g0006129 | Lj1g0006129.3 | Lotus japonicus | 87% | 0.0 | 59.61% | 726 |
| Lj1g0011717 | Lj1g0011717.1 | | 78% | 0.0 | 50.77% | 680 |
| Lj1g0011876 | Lj1g0011876.1 | | 93% | 0.0 | 54.94% | 699 |
| Lj1g0018523 | Lj1g0018523.1 | | 86% | 0.0 | 52.17% | 732 |
| Lj3g0008315 | Lj3g0008315.1 | | 87% | 0.0 | 48.93% | 613 |
| Lj5g0014423 | Lj5g0014423.1 | | 89% | 0.0 | 62.03% | 706 |

| | | | | | | |
|---|---|---|---|---|---|---|
| Lalb_Chr01g0020141 | Lalb_Chr01g0020141 | Lupinus albus | 93% | 0.0 | 54.83% | 690 |
| Lalb_Chr05g0210891 | Lalb_Chr05g0210891 | | 88% | 0.0 | 61.13% | 694 |
| Lalb_Chr16g0376391 | Lalb_Chr16g0376391 | | 89% | 0.0 | 60.55% | 687 |
| Mamar.0011s0826 | Mamar.0011s0826.1 | Malcolmia maritima | 100% | 0.0 | 89.99% | 739 |
| Mamar.0050s1498 | Mamar.0050s1498.1 | | 94% | 0.0 | 60.46% | 710 |
| Mamar.0016s0220 | Mamar.0016s0220.1 | | 94% | 0.0 | 55.71% | 698 |
| Mapoly0013s0109 | Mapoly0013s0109.1 | Marchantia polymorpha | 92% | 0.0 | 54.47% | 751 |
| Medtr1g104870 | Medtr1g104870.1 | Medicago truncatula | 94% | 0.0 | 59.94% | 704 |
| Medtr5g005940 | Medtr5g005940.1 | | 78% | 0.0 | 61.45% | 601 |
| | Medtr5g005940.2 | | 78% | 0.0 | 61.45% | 696 |
| Medtr7g094900 | Medtr7g094900.1 | | 83% | 0.0 | 61.26% | 716 |
| Medtr7g094910 | Medtr7g094910.1 | | 84% | 0.0 | 60.61% | 697 |
| | Medtr7g094910.2 | | 75% | 0.0 | 62.46% | 683 |
| MgTOL.I00297 | MgTOL.I00297.1 | Mimulus guttatus | 91% | 0.0 | 59.12% | 698 |
| MgTOL.N2070 | MgTOL.N2070.1 | | 94% | 0.0 | 60.40% | 715 |
| Myper.0004s1359 | Myper.0004s1359.1 | Myagrum perfoliatum | 99% | 0.0 | 84.08% | 736 |
| Myper.0004s1981 | Myper.0004s1981.1 | | 94% | 0.0 | 60.20% | 710 |
| Nycol.K00850 | Nycol.K00850.1 | Nymphaea colorata | 93% | 0.0 | 56.73% | 698 |
| Oeu041702 | Oeu041702.3 | Olea europaea | 89% | 0.0 | 63.69% | 716 |
| Oeu028070 | Oeu028070.1 | | 89% | 0.0 | 60.58% | 751 |
| Oropetium_20150105_05154 | Oropetium_20150105_05154A | Oropetium thomaeum | 88% | 3E-116 | 70.45% | 587 |
| PhHAL.3G279400 | PhHAL.3G279400.1 | Panicum hallii | 88% | 0.0 | 60.15% | 742 |
| Podel.08G175100 | Podel.08G175100.1 | Populus deltoides | 87% | 0.0 | 63.46% | 718 |
| Podel.10G086700 | Podel.10G086700.1 | | 87% | 0.0 | 63.00% | 717 |
| Podel.05G221700 | Podel.05G221700.1 | | 89% | 0.0 | 62.33% | 698 |
| Podel.02G058100 | Podel.02G058100.1 | | 88% | 0.0 | 61.37% | 704 |
| Ptrif.0009s0499 | Ptrif.0009s0499.1 | Poncirus trifoliata | 89% | 0.0 | 62.71% | 717 |
| Ptrif.0003s3479 | Ptrif.0003s3479.1 | | 94% | 0.0 | 61.69% | 704 |
| Pp3c7_11520 | Pp3c7_11520V3.1 | Physcomitrium patens | 69% | 0.0 | 61.13% | 551 |
| | Pp3c7_11520V3.2 | | 86% | 0.0 | 60.78% | 674 |
| | Pp3c7_11520V3.3 | | 93% | 0.0 | 56.44% | 709 |
| | Pp3c7_11520V3.4 | | 93% | 0.0 | 56.69% | 706 |
| | Pp3c7_11520V3.5 | | 93% | 0.0 | 56.69% | 706 |
| Pp3c11_17010 | Pp3c11_17010V3.1 | | 90% | 0.0 | 57.31% | 709 |
| | Pp3c11_17010V3.2 | | 90% | 0.0 | 55.73% | 728 |
| | Pp3c11_17010V3.3 | | 90% | 0.0 | 55.49% | 731 |
| | Pp3c11_17010V3.4 | | 90% | 0.0 | 57.31% | 709 |
| | Pp3c11_17010V3.5 | | 90% | 0.0 | 57.57% | 706 |
| Phacu.WLD.002G006700 | Phacu.WLD.002G006700.1 | Phaseolus acutifolius | 89% | 0.0 | 62.22% | 707 |
| | Phacu.WLD.002G006700.1 | | 89% | 0.0 | 62.22% | 707 |
| | Phacu.WLD.002G006700.2 | | 86% | 0.0 | 63.49% | 694 |
| Phacu.WLD.007G055800 | Phacu.WLD.007G055800.1 | | 94% | 0.0 | 60.11% | 707 |
| | Phacu.WLD.007G055800.2 | | 90% | 0.0 | 61.07% | 714 |
| Phacu.WLD.001G179100 | Phacu.WLD.001G179100.1 | | 88% | 0.0 | 52.75% | 706 |
| | Phacu.WLD.001G179100.2 | | 79% | 0.0 | 54.30% | 577 |
| Phlat.07G120000 | Phlat.07G120000.1 | Pharus latifolius | 88% | 0.0 | 60.82% | 731 |
| Pavag09G136000 | Pavag09G136000.1 | Paspalum vaginatum | 88% | 0.0 | 61.31% | 741 |
| Pavir.3KG262426 | Pavir.3KG262426.1 | Panicum virgatum | 84% | 0.0 | 60.93% | 741 |
| | Pavir.3KG262426.2 | | 84% | 0.0 | 60.93% | 739 |
| | Pavir.3KG262426.3 | | 75% | 0.0 | 63.65% | 685 |
| | Pavir.3KG262426.3 | | 75% | 0.0 | 63.65% | 716 |
| | Pavir.3KG262426.4 | | 71% | 0.0 | 62.62% | 662 |
| Pavir.3NG076958 | Pavir.3NG076958.1 | | 85% | 0.0 | 61.60% | 741 |
| | Pavir.3NG076958.2 | | 85% | 0.0 | 61.60% | 739 |
| | Pavir.3NG076958.3 | | 85% | 0.0 | 60.34% | 726 |
| | Pavir.3NG076958.4 | | 83% | 0.0 | 62.08% | 722 |
| Poman.08G130800 | Poman.08G130800.1 | Populus nigra x maximowiczii | 87% | 0.0 | 63.76% | 717 |
| Poman.05G171800 | Poman.05G171800.1 | | 89% | 0.0 | 62.63% | 708 |
| | Poman.05G171800.2 | | 86% | 0.0 | 63.58% | 680 |
| Poman.10G071900 | Poman.10G071900.1 | | 87% | 0.0 | 63.00% | 717 |
| Poman.02G047100 | Poman.02G047100.1 | | 88% | 0.0 | 61.22% | 704 |
| | Poman.02G047100.2 | | 71% | 0.0 | 62.48% | 648 |
| | Poman.02G047100.3 | | 71% | 0.0 | 62.48% | 648 |
| | Poman.02G047100.4 | | 79% | 0.0 | 63.61% | 736 |
| PtXaAlbH.05G161100 | PtXaAlbH.05G161100.1 | | 89% | 0.0 | 62.31% | 696 |
| PtXaAlbH.10G070200 | PtXaAlbH.10G070200.1 | | 87% | 0.0 | 61.78% | 725 |
| | PtXaAlbH.10G070200.2 | Populus tremula x Populus alba | 87% | 0.0 | 62.54% | 717 |
| | PtXaAlbH.10G070200.3 | | 87% | 0.0 | 62.54% | 716 |
| | PtXaAlbH.10G070200.4 | | 87% | 0.0 | 61.62% | 709 |
| | PtXaAlbH.10G070200.5 | | 87% | 0.0 | 62.54% | 715 |
| | PtXaAlbH.10G070200.6 | | 87% | 0.0 | 62.23% | 713 |
| PtXaAlbH.02G045800 | PtXaAlbH.02G045800.1 | | 88% | 0.0 | 61.19% | 706 |
| | PtXaAlbH.02G045800.2 | | 88% | 0.0 | 61.37% | 796 |
| PtXaAlbH.08G127900 | PtXaAlbH.08G127900.1 | | 94% | 0.0 | 60.08% | 717 |
| | PtXaAlbH.08G127900.1 | | 94% | 0.0 | 60.08% | 717 |
| | PtXaAlbH.08G127900.2 | | 94% | 0.0 | 60.08% | 717 |
| | PtXaAlbH.08G127900.3 | | 94% | 0.0 | 60.08% | 717 |
| | PtXaAlbH.08G127900.4 | | 94% | 0.0 | 60.08% | 717 |
| | PtXaAlbH.08G127900.5 | | 94% | 0.0 | 60.08% | 717 |
| Potri.008G150100 | Potri.008G150100.1 | Populus trichocarpa | 87% | 0.0 | 63.76% | 717 |
| | Potri.008G150100.2 | | 84% | 0.0 | 64.71% | 704 |
| | Potri.008G150100.1 | | 87% | 0.0 | 63.76% | 717 |
| | Potri.008G150100.2 | | 84% | 0.0 | 64.71% | 689 |
| | Potri.008G150100.3 | | 79% | 0.0 | 65.58% | 665 |
| | Potri.008G150100.4 | | 86% | 0.0 | 68.38% | 529 |
| | Potri.008G150100.5 | | 79% | 0.0 | 64.98% | 661 |
| | Potri.008G150100.6 | | 84% | 0.0 | 64.56% | 713 |
| | Potri.008G150100.6 | | 84% | 0.0 | 64.56% | 704 |
| Potri.005G210000 | Potri.005G210000.1 | | 89% | 0.0 | 62.33% | 698 |
| Potri.002G052700 | Potri.002G052700.2 | | 88% | 0.0 | 61.37% | 704 |
| Potri.010G090900 | Potri.010G090900.1 | | 87% | 0.0 | 62.10% | 713 |
| | Potri.010G090900.2 | | 87% | 0.0 | 62.84% | 713 |
| | Potri.010G090900.3 | | 87% | 0.0 | 61.31% | 781 |
| | Potri.010G090900.4 | | 72% | 0.0 | 62.94% | 648 |
| | Potri.010G090900.7 | | 77% | 0.0 | 62.71% | 709 |
| | Potri.010G090900.8 | | 72% | 0.0 | 62.02% | 642 |
| | Potri.010G090900.9 | | 77% | 0.0 | 63.59% | 716 |
| | Potri.010G090900.10 | | 72% | 0.0 | 62.94% | 648 |
| | Potri.010G090900.11 | | 72% | 0.0 | 62.94% | 648 |
| | Potri.010G090900.12 | | 74% | 0.0 | 64.13% | 694 |
| | Potri.010G090900.13 | | 67% | 0.0 | 63.55% | 609 |
| | Potri.010G090900.14 | | 70% | 0.0 | 67.76% | 590 |
| FUN_042764 | FUN_042764-T1 | Portulaca amilis | 87% | 0.0 | 61.55% | 723 |
| FUN_014427 | FUN_014427-T1 | | 94% | 0.0 | 55.92% | 690 |
| | FUN_014427-T2 | | 77% | 0.0 | 55.65% | 601 |
| Prupe.1G155800 | Prupe.1G155800.1 | Prunus persica | 94% | 0.0 | 59.71% | 704 |
| | Prupe.1G155800.2 | | 94% | 0.0 | 59.71% | 704 |
| Prupe.8G240700 | Prupe.8G240700.1 | | 95% | 0.0 | 60.25% | 710 |
| Qurub.06G239500 | Qurub.06G239500.1 | Quercus rubra | 89% | 0.0 | 63.73% | 732 |
| Qurub.12G088800 | Qurub.12G088800.1 | | 93% | 0.0 | 61.75% | 708 |
| Roisl.0210s0994 | Roisl.0210s0994.1 | Rorippa islandica | 99% | 0.0 | 84.98% | 739 |
| Roisl.0210s1650 | Roisl.0210s1650.1 | | 94% | 0.0 | 60.62% | 711 |
| | Roisl.0210s1650.2 | | 94% | 0.0 | 60.57% | 711 |
| Roisl.0089s0140 | Roisl.0089s0140.1 | | 87% | 0.0 | 57.52% | 701 |
| Sapur.005G165700 | Sapur.005G165700.1 | Salix purpurea | 88% | 0.0 | 62.33% | 701 |
| | Sapur.005G165700.2 | | 71% | 0.0 | 63.81% | 658 |
| Sapur.010G058700 | Sapur.010G058700.1 | | 87% | 0.0 | 62.08% | 713 |
| | Sapur.010G058700.2 | | 86% | 0.0 | 65.14% | 712 |
| | Sapur.010G058700.3 | | 61% | 0.0 | 63.39% | 605 |
| Sapur.002G041300 | Sapur.002G041300.1 | | 88% | 0.0 | 61.68% | 704 |
| | Sapur.002G041300.2 | | 79% | 0.0 | 64.29% | 701 |
| Sapur.008G121400 | Sapur.008G121400.1 | | 89% | 0.0 | 62.29% | 717 |
| | Sapur.008G121400.2 | | 86% | 0.0 | 67.12% | 711 |
| | Sapur.008G121400.3 | | 85% | 0.0 | 63.51% | 701 |

| | | | | | | |
|---|---|---|---|---|---|---|
| Sp6g30920 | Sp6g30920.1 | Schrenkiella parvula | 96% | 0.0 | 84.95% | 711 |
| Sp6g25080 | Sp6g25080.1 | | 94% | 0.0 | 58.92% | 710 |
| 164824 | 164824 | Selaginella moellendorffii | 87% | 0.0 | 58.09% | 701 |
| Sialb.0278s0087 | Sialb.0278s0087.1 | Sinapis alba | 96% | 0.0 | 84.79% | 710 |
| Sialb.0009s1174 | Sialb.0009s1174.1 | | 96% | 0.0 | 84.93% | 710 |
| Sialb.0193s0098 | Sialb.0193s0098.1 | | 94% | 0.0 | 59.69% | 711 |
| Sialb.0056s0376 | Sialb.0056s0376.1 | | 94% | 0.0 | 59.74% | 712 |
| Sialb.0056s0376 | Sialb.0056s0376.2 | | 94% | 0.0 | 59.74% | 712 |
| Sphfalx01G181900 | Sphfalx01G181900.1 | Sphagnum fallax | 89% | 0.0 | 57.51% | 714 |
| | Sphfalx01G181900.2 | | 89% | 0.0 | 57.77% | 711 |
| Sphfalx02G196900 | Sphfalx02G196900.1 | | 94% | 0.0 | 55.87% | 713 |
| | Sphfalx02G196900.2 | | 94% | 0.0 | 56.12% | 710 |
| Sphmag02G198100 | Sphmag02G198100.1 | Sphagnum magellanicum | 89% | 0.0 | 57.88% | 713 |
| | Sphmag02G198100.2 | | 89% | 0.0 | 58.14% | 710 |
| Sphmag01G172800 | Sphmag01G172800.1 | | 89% | 0.0 | 57.51% | 714 |
| | Sphmag01G172800.2 | | 89% | 0.0 | 57.77% | 711 |
| Spov3_chr4.02900 | Spov3_chr4.02900 | Spinacia oleracea | 94% | 0.0 | 58.40% | 709 |
| Spov3_chr3.03740 | Spov3_chr3.03740 | | 88% | 0.0 | 56.75% | 659 |
| Spipo22G0007800 | Spipo22G0007800 | Spirodela polyrhiza | 83% | 0.0 | 63.41% | 680 |
| Thint.S01G292400 | Thint.S01G292400.1 | Thinopyrum intermedium | 89% | 0.0 | 60.30% | 740 |
| Thint.J01G305100 | Thint.J01G305100.1 | | 89% | 0.0 | 60.30% | 742 |
| Thint.V01G294800 | Thint.V01G294800.1 | | 88% | 0.0 | 60.52% | 743 |
| Thupl.29378015s0033 | Thupl.29378015s0033.1 | Thuja plicata | 90% | 0.0 | 57.74% | 819 |
| Tp57577_TGAC_v2_gene9920 | Tp57577_TGAC_v2_mRNA10257 | Trifolium pratense | 80% | 0.0 | 62.12% | 578 |
| Tp57577_TGAC_v2_gene37872 | Tp57577_TGAC_v2_mRNA39151 | | 88% | 0.0 | 62.16% | 737 |
| Tp57577_TGAC_v2_gene5509 | Tp57577_TGAC_v2_mRNA5699 | | 92% | 0.0 | 60.35% | 704 |
| Urofu.3G293300 | Urofu.3G293300.1 | Urochloa fusca | 88% | 0.0 | 60.46% | 746 |
| | Urofu.3G293300.2 | | 88% | 0.0 | 60.46% | 744 |
| | Urofu.3G293300.3 | | 79% | 0.0 | 62.84% | 683 |
| | Urofu.3G293300.4 | | 87% | 0.0 | 64.68% | 517 |
| | Urofu.3G293300.5 | | 87% | 0.0 | 64.68% | 517 |
| | Urofu.3G293300.6 | | 61% | 0.0 | 63.50% | 465 |
| Vadar_g31390 | Vadar_g31390.t1 | Vaccinium darrowii | 94% | 0.0 | 61.80% | 706 |
| Vadar_g42337 | Vadar_g42337.t1 | | 94% | 0.0 | 60.66% | 703 |
| Vigun02g197000 | Vigun02g197000.1 | Vigna unguiculata | 89% | 0.0 | 62.31% | 778 |
| | Vigun02g197000.3 | | 89% | 0.0 | 62.31% | 778 |
| | Vigun02g197000.4 | | 89% | 0.0 | 62.31% | 778 |
| | Vigun02g197000.5 | | 89% | 0.0 | 62.31% | 778 |
| Vigun07g242500 | Vigun07g242500.1 | | 94% | 0.0 | 60.00% | 706 |
| | Vigun07g242500.2 | | 94% | 0.0 | 60.00% | 706 |
| | Vigun07g242500.3 | | 94% | 0.0 | 60.00% | 706 |
| Zosma02g23080 | Zosma02g23080.1 | Zostera marina | 84% | 0.0 | 57.91% | 698 |
| Zosma01g37860 | Zosma01g37860.1 | | 85% | 0.0 | 60.34% | 701 |
| Nitab4.5_0000458g0070 | Nitab4.5_0000458g0070 | Nicotiana tabacum | 95% | 0.0 | 57.28% | 729 |
| Nitab4.5_0000759g0190 | Nitab4.5_0000759g0190 | | 89% | 0.0 | 59.64% | 680 |
| Nitab4.5_0006749g0030 | Nitab4.5_0006749g0030 | | 80% | 0.0 | 58.71% | 617 |
| Nitab4.5_0008797g0060 | Nitab4.5_0008797g0060 | | 89% | 0.0 | 57.38% | 660 |
| NbS00014381g0003 | NbS00014381g0003.1 | Nicotiana benthamiana | 91% | 0.0 | 60.09% | 691 |
| NbS00024271g0016 | NbS00024271g0016.1 | | 91% | 0.0 | 60.56% | 677 |
| NbS00032920g0003 | NbS00032920g0003.1 | | 85% | 0.0 | 60.88% | 651 |
| NbS00022240g0022 | NbS00022240g0022.1 | | 86% | 0.0 | 57.50% | 697 |

| | | | | | | |
|---|---|---|---|---|---|---|
| PtXaAlbH.03G148200 | PtXaAlbH.03G148200.1 | *Populus tremula x Populus alba* | 95% | 6E-89 | 68.78% | 220 |
| | PtXaAlbH.03G148200.2 | | 90% | 1E-77 | 65.28% | 207 |
| | PtXaAlbH.03G148200.3 | | 63% | 5E-72 | 80.74% | 179 |
| PtXaAlbH.01G035600 | PtXaAlbH.01G035600.1 | | 95% | 6E-85 | 65.85% | 221 |
| Potri.003G184925 | Potri.003G184925.1 | *Populus trichocarpa* | 95% | 8E-89 | 68.78% | 220 |
| | Potri.003G184925.2 | | 63% | 2E-72 | 80.74% | 152 |
| Potri.001G041900 | Potri.001G041900.1 | | 95% | 5E-87 | 67.80% | 221 |
| | Potri.001G041900.5 | | 64% | 7E-73 | 78.99% | 152 |
| FUN_040438 | FUN_040438-T1 | *Portulaca amilis* | 93% | 2E-64 | 57.28% | 211 |
| Prupe.3G103600 | Prupe.3G103600.1 | *Prunus persica* | 93% | 2E-86 | 67.98% | 210 |
| | Prupe.3G103600.2 | | 66% | 9E-74 | 77.93% | 147 |
| | Prupe.3G103600.3 | | 66% | 9E-74 | 77.93% | 147 |
| Qurub.05G179800 | Qurub.05G179800.1 | *Quercus rubra* | 96% | 4E-86 | 62.98% | 220 |
| | Qurub.05G179800.2 | | 81% | 1E-74 | 65.90% | 177 |
| | Qurub.05G179800.3 | | 81% | 1E-74 | 65.90% | 177 |
| 30131.t000042 | 30131.m006891 | *Ricinus communis* | 100% | 1E-92 | 62.33% | 230 |
| Roisl.0115s0146 | Roisl.0115s0146.1 | *Rorippa islandica* | 95% | 1E-126 | 89.37% | 216 |
| Sspon.04G0001710-2C | Sspon.04G0001710-2C | *Saccharum spontaneum* | 89% | 3E-66 | 59.61% | 319 |
| Sspon.04G0001710-3D | Sspon.04G0001710-3D | | 89% | 6E-66 | 59.11% | 313 |
| Sapur.003G140400 | Sapur.003G140400.1 | *Salix purpurea* | 95% | 2E-85 | 66.34% | 220 |
| Sapur.001G032000 | Sapur.001G032000.1 | | 95% | 4E-82 | 62.93% | 219 |
| Sp5g34800 | Sp5g34800.1 | *Schrenkiella parvula* | 100% | 3E-130 | 90.14% | 209 |
| GWHGASIY033783 | GWHGASIY033783 | *Secale cereale* | 86% | 1E-65 | 57.62% | 220 |
| GWHGASIY033046 | GWHGASIY033046 | | 86% | 2E-65 | 57.14% | 220 |
| 5289 | 5289 | *Selaginella moellendorffii* | 62% | 4E-61 | 65.67% | 136 |
| Sevir.1G355800 | Sevir.1G355800.1 | *Setaria viridis* | 89% | 8E-68 | 58.82% | 207 |
| | Sevir.1G355800.2 | | 89% | 8E-68 | 58.82% | 207 |
| Sialb.0005s0947 | Sialb.0005s0947.1 | *Sinapis alba* | 100% | 2E-120 | 84.23% | 221 |
| | Sialb.0005s0947.2 | | 100% | 2E-121 | 84.16% | 220 |
| Sialb.0005s0646 | Sialb.0005s0646.1 | | 96% | 2E-119 | 85.71% | 213 |
| | Sialb.0005s0646.2 | | 96% | 1E-120 | 86.12% | 212 |
| Solyc03g117670 | Solyc03g117670.4 | *Solanum lycopersicum* | 88% | 2E-83 | 64.06% | 241 |
| PGSC0003DMG400014155 | PGSC0003DMT400036704 | *Solanum tuberosum* | 73% | 1,00E-73 | 66.67% | 270 |
| | PGSC0003DMT400036701 | | 47% | 8,00E-43 | 66.67% | 225 |
| | PGSC0003DMT400036702 | | 73% | 7,00E-74 | 66.67% | 264 |
| | PGSC0003DMT400036703 | | 47% | 1,00E-42 | 66.67% | 223 |
| Sobic.004G328000 | Sobic.004G328000.3 | *Sorghum bicolor* | 89% | 1E-67 | 59.90% | 207 |
| Sphfalx03G101000 | Sphfalx03G101000.1 | *Sphagnum fallax* | 63% | 5E-60 | 67.41% | 322 |
| Sphfalx18G098200 | Sphfalx18G098200.1 | | 61% | 1E-55 | 66.15% | 344 |
| | Sphfalx18G098200.2 | | 61% | 2,00E-57 | 66.15% | 316 |
| | Sphfalx18G098200.3 | | 33% | 1,00E-30 | 68.06% | 253 |
| | Sphfalx18G098200.4 | | 33% | 1,00E-30 | 68.06% | 282 |
| Sphmag03G095600 | Sphmag03G095600.1 | *Sphagnum magellanicum* | 63% | 6E-60 | 67.41% | 322 |
| Sphmag18G097300 | Sphmag18G097300.1 | | 62% | 2E-55 | 66.92% | 344 |
| Spov3_chr5.02035 | Spov3_chr5.02035 | *Spinacia oleracea* | 86% | 1E-74 | 61.38% | 208 |
| Spipo28G0013100 | Spipo28G0013100 | *Spirodela polyrhiza* | 90% | 8,00E-68 | 53.59% | 244 |
| Thecc.06G046600 | Thecc.06G046600.1 | *Theobroma cacao* | 97% | 9E-91 | 68.75% | 209 |
| | Thecc.06G046600.2 | | 75% | 8,00E-78 | 73.91% | 163 |
| | Thecc.06G046600.3 | | 70% | 1,00E-75 | 76.67% | 181 |
| Thint.S06G379700 | Thint.S06G379700.1 | *Thinopyrum intermedium* | 88% | 7E-67 | 56.28% | 220 |
| | Thint.S06G379700.2 | | 88% | 7E-67 | 56.28% | 220 |
| | Thint.S06G379700.3 | | 71% | 2E-67 | 69.87% | 166 |
| Thint.J06G419300 | Thint.J06G419300.1 | | 71% | 8E-66 | 69.23% | 219 |
| | Thint.J06G419300.2 | | 71% | 2E-66 | 69.23% | 165 |
| Thlar.0008s0131 | Thlar.0008s0131.1 | *Thlaspi arvense* | 95% | 7E-132 | 90.87% | 217 |
| Thupl.29377465s0003 | Thupl.29377465s0003.1 | *Thuja plicata* | 61% | 5E-60 | 67.69% | 191 |
| | Thupl.29377465s0003.2 | | 61% | 5E-60 | 67.69% | 191 |
| | Thupl.29377465s0003.4 | | 61% | 5E-60 | 67.69% | 192 |
| | Thupl.29377465s0003.5 | | 61% | 5E-60 | 67.69% | 191 |
| | Thupl.29377465s0003.6 | | 61% | 5E-60 | 67.69% | 191 |
| | Thupl.29377465s0003.8 | | 61% | 5E-60 | 67.69% | 192 |
| | Thupl.29377465s0003.11 | | 61% | 5E-60 | 67.69% | 192 |
| Tp57577_TGAC_v2_gene3367 | Tp57577_TGAC_v2_mRNA3464 | *Trifolium pratense* | 94% | 3E-90 | 63.59% | 210 |
| TraesCS6D03G0782500 | TraesCS6D03G0782500.1 | *Triticum aestivum* | 71% | 1E-65 | 69.23% | 219 |
| TraesCS6A03G0911900 | TraesCS6A03G0911900.1 | | 71% | 1E-65 | 69.87% | 292 |
| TraesCS6B03G1096200 | TraesCS6B03G1096200.1 | | 71% | 2E-65 | 69.23% | 219 |
| Urofu.1G327800 | Urofu.1G327800.1 | *Urochloa fusca* | 71% | 1E-66 | 70.06% | 211 |
| | Urofu.1G327800.2 | | 89% | 2E-70 | 60.40% | 207 |
| | Urofu.1G327800.3 | | 59% | 2E-39 | 57.04% | 183 |
| | Urofu.1G327800.4 | | 89% | 2E-70 | 60.40% | 207 |
| | Urofu.1G327800.5 | | 41% | 1E-36 | 72.22% | 187 |
| Vadar_g14679 | Vadar_g14679.t1 | *Vaccinium darrowii* | 94% | 3E-86 | 58.41% | 228 |
| Vigun08g128600 | Vigun08g128600.1 | *Vigna unguiculata* | 95% | 2E-84 | 61.26% | 225 |
| | Vigun08g128600.2 | | 95% | 7E-86 | 63.51% | 214 |
| GSVIVG01016103001 | GSVIVG01016103001 | *Vitis vinifera* | 91% | 4E-81 | 63.45% | 201 |
| Zm00001d018364 | Zm00001d018364_T001 | *Zea mays B73* | 89% | 4E-67 | 58.25% | 211 |
| Zm00007a00050657 | Zm00007a00050657 | *Zea mays B104* | 89% | 1E-65 | 57.21% | 291 |
| Zosma03g12720 | Zosma03g12720.1 | *Zostera marina* | 59% | 5E-55 | 67.72% | 585 |

**Supplemental Table S4.** Attributions for the main vibrational bands identified in the bio-fingerprint region.

| Band ($cm^{-1}$) | Stretch vibrations | Assignment | | Reference |
|---|---|---|---|---|
| **1744** | C=O | Lipids | | Durak & Depciuch, 2020 |
| **1636** | C=O | Protein (Amide I) | | Tessaro et al., 2022 |
| **1535** | N-H | Lipids and protein (Amide II) | | Durak & Depciuch, 2020 |
| **1439** | C-H | Lipids and protein | | Durak & Depciuch, 2020 |
| **1242** | C-O | Protein | | Durak & Depciuch, 2020 |
| **1230-1235** | O-H | Carbohydrate | Lignin | Dinant et al., 2019 |
| **1090 - 1100** | C-O and C-C | | Pectin | Liu et al., 2021 |
| **1160** | C-H | | Lignin | Dinant et al., 2019 |
| **1065** | C-O | | Sucrose | Hashimoto et al., 2005 |
| **1055** | C-OH | | Sucrose | Hashimoto et al 2005 |
| **1052** | C-H | | Crystalline Cellulose | Liu et al., 2021 |
| **1024** | C-O | | Cellulose | López-Malvar et al., 2021 |
| **1064** | C=C | | Hemicellulose | Liu et al., 2021 |
| **991** | α,α-1,1 linkage | | trehalose | San-Blas et al., 2011 |
| **972** | $O-CH_3$ | | Pectin | Schulz & Baranska, 2007 |
| **954** | C-O-C | | Amylose | Liu et al., 2004 |
